# An updated catalogue of split-GAL4 driver lines for descending neurons in *Drosophila melanogaster*

**DOI:** 10.1101/2025.02.22.639679

**Authors:** Jessica L. Zung, Shigehiro Namiki, Geoffrey W. Meissner, Han S.J. Cheong, Marta Costa, Katharina Eichler, Tomke Stürner, Gregory S.X.E. Jefferis, Claire Managan, FlyLight Project Team, Wyatt Korff, Gwyneth M. Card

**Affiliations:** Janelia Research Campus, Howard Hughes Medical Institute, Ashburn, VA, USA; Howard Hughes Medical Institute and Zuckerman Mind Brain Behavior Institute, Columbia University, New York, NY, USA; Department of Biology, West Virginia University, Morgantown, WV, USA; Drosophila Connectomics Group, Department of Zoology, University of Cambridge, Cambridge, UK; Genetics Department, Leipzig University, Leipzig, Germany; Neurobiology Division, MRC Laboratory of Molecular Biology, Cambridge, UK

**Author notes:** Corresponding authors; WK; GMC. Present address: Research Center for Advanced Science and Technology, University of Tokyo, Tokyo, Japan (Shigehiro Namiki). Equal contribution (Jessica L. Zung, Shigehiro Namiki).

## Abstract

Descending neurons (DNs) occupy a key position in the sensorimotor hierarchy, conveying signals from the brain to the rest of the body below the neck. In *Drosophila melanogaster* flies, approximately 480 DN cell types have been described from electron-microscopy image datasets. Genetic access to these cell types is crucial for further investigation of their role in generating behaviour. We previously conducted the first large-scale survey of *Drosophila melanogaster* DNs, describing 98 unique cell types from light microscopy and generating cell-type-specific split-Gal4 driver lines for 65 of them. Here, we extend our previous work, describing the morphology of 146 additional DN types from light microscopy, bringing the total number DN types identified in light microscopy datasets to 244, or roughly 50% of all DN types. In addition, we produced 500 new sparse split-Gal4 driver lines and compiled a list of previously published DN lines from the literature for a combined list of 806 split-Gal4 driver lines targeting 190 DN types.

## Introduction

A common feature of nervous systems across most of the animal kingdom is the presence of a brain, which coordinates most sensory processing and decision making, and a separate nerve cord—a ventral nerve cord or spinal cord—that performs final coordination of motor output. Communication from the brain to the nerve cord is relayed via a relatively small population of descending neurons (DNs). The privileged position of these neurons in the sensorimotor hierarchy makes them prime targets for the study of behaviour and motor control.

In *Drosophila melanogaster*, genetic resources and stereotypy of neurons across individuals allow us to generate cell-type-specific driver lines that help reveal the morphology of individual neurons and enable targeted manipulation or measurement of neurons. We previously created 178 sparse split-Gal4 driver lines targeting 65 DN types and described an additional 33 DN types labelled by more broadly expressing Gal4 lines (Namiki et al. 2018). This covered roughly half of the ∼700 total DNs we estimated at the time based on counting DN cell bodies using a pan-neuronally expressed photoactivatable GFP.

Recent analyses based on electron-microscopy (EM) images have since revealed that there are in fact ∼1300 DNs split across ∼480 types in *Drosophila melanogaster* males (Cheong et al. 2024) and a similar number in females (Stuerner et al. 2024). These exhaustive catalogues underscore the value of EM-based connectomes for delivering a more complete picture of animal nervous systems. However, genetic driver lines are still required for *in vivo* manipulation or measurement of neurons. Indeed, our previous collection of descending-neuron lines has already been used to study *Drosophila* escape behaviour (Dombrovski et al. 2023), saccades during flight (Ros et al. 2024), walking initiation (Bidaye et al. 2020), steering during flight (Namiki et al. 2022) and walking (Chen et al. 2018; Rayshubskiy et al. 2024), freezing (Zacarias et al. 2018), landing (Ache et al. 2019), grooming (Guo et al. 2022), and diverse other behaviours (Cande et al. 2018). Here, we extend our previous work, describing an additional 146 DN types from light-level microscopy data and producing 500 additional DN driver lines. We also survey the literature for previously published DN lines to compile a combined list of 806 split- Gal4 driver lines targeting 190 DN types.

## Results

### Identification of new DN types

Our first goal was to individually characterize the morphology of descending neurons using a large database of light-microscopy (LM) brain and ventral nerve cord (VNC) images (Meissner et al. 2023). Although new EM datasets show neuron morphology with greater spatial resolution, all EM datasets published to date terminate at the neck (Stuerner et al. 2024), limiting their utility for describing complete DN morphology. We previously described 98 DN types from an initial survey of this LM database (Namiki et al. 2018) and now describe the results of our completed search below.

We manually searched publicly available images in Janelia’s “Gen1” collection of broadly expressing Gal4 driver lines (Jenett et al. 2012; Tirian and Dickson 2017; Meissner et al. 2023). We specifically searched for lines that showed expression in the brain and VNC, using the MultiColor FlpOut (MCFO) technique to label neurons sparsely enough for easy visualization (Nern et al. 2015; Meissner et al. 2023). We identified DNs from these images, operationally defining them as in Namiki et al. (2018) as neurons with a soma in the brain and processes extending to the VNC (Fig. 1a,b). Using these criteria, we characterized the morphology of 122 new DN types from light-level microscopy and named them following previous convention (Namiki et al. 2018). We also documented 24 types that had been described previously in other work (see Supplementary Table 1), for a total of 146 new types that were not covered in our 2018 survey.

**Figure 1.**
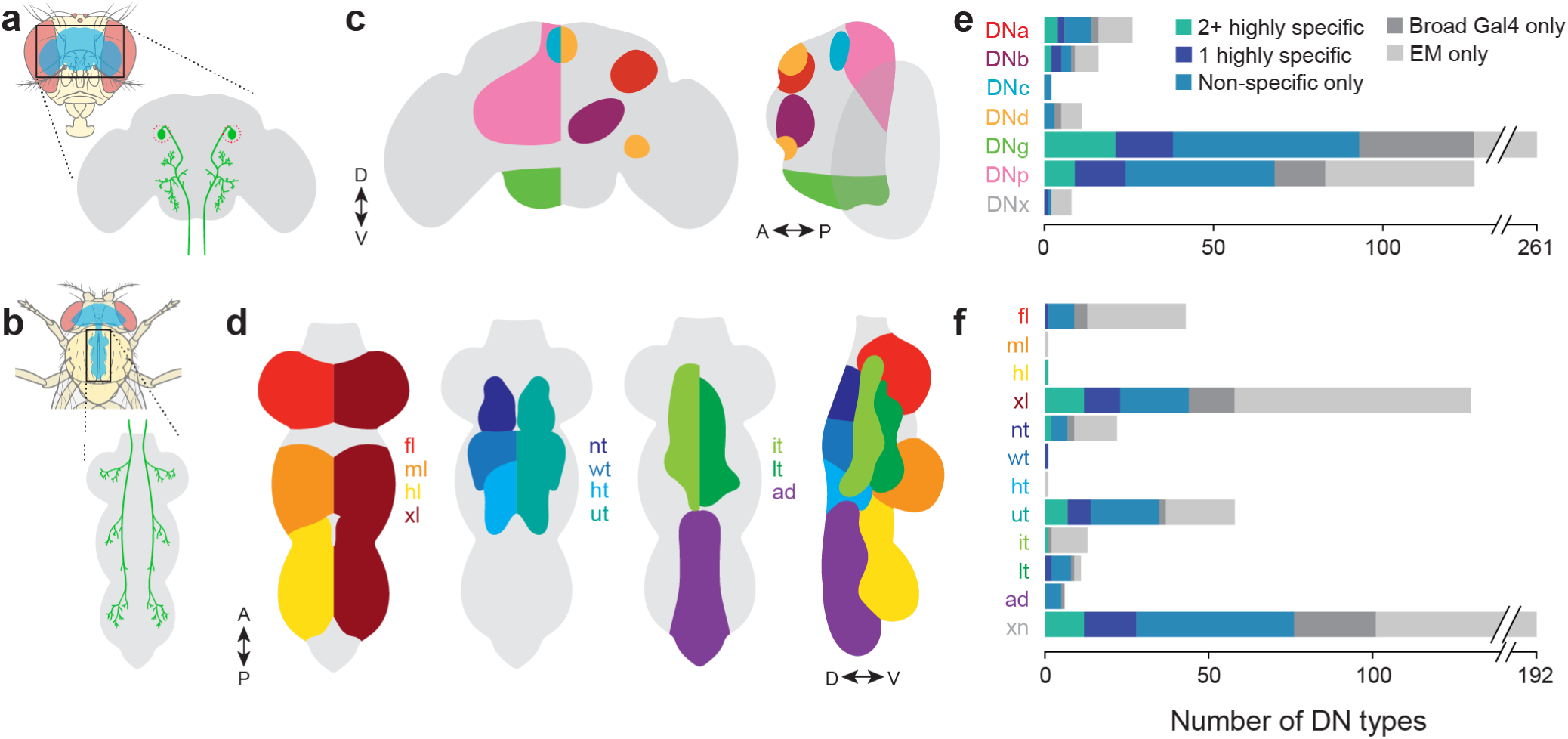
(**a,b**) Upper right schematics show in blue the locations of the brain (a; frontal head view) and VNC (b; top view of head and thorax) within a fly body. Lower right schematics show the general anatomy of a DN, including a soma in the brain (red dotted circle) (a) and processes in the VNC (b). Fly cartoons from Wikimedia Commons. (**c**) Schematic showing the approximate location of DN somas according to the naming scheme used in Namiki et al. (2018). For clarity, the left-hand illustration shows locations of each soma group on only one side of the bilaterally symmetric brain. DNx somas (not shown) are outside the brain. (**d**) Schematic showing the approximate location of VNC neuropils by which DNs were categorized systematically in Cheong et al. (2024). As in (b), the dorsal/ventral views often show neuropil location on only one side of the bilaterally symmetric VNC. ‘xn’ DN types (not shown) innervate multiple neuropils. (**e**) Bar plot showing the number of DN types in each soma- location category. Coloured bars indicate the number of types for which there exist at least 2 highly specific (‘A’) lines (teal), 1 highly specific line (dark blue), only non-specific or inconsistently expressing lines (‘B’ or ‘C’) (light blue), or no split-Gal4 lines (grey). DN types with no corresponding split-Gal4 lines are further separated by whether they were ever identified from light microscopy images in a very broadly expressing Gal4 line (dark grey) or identified only from electron microscopy images (light grey). (**f**) Bar plot showing the number of DN types innervating each VNC neuropil. As in (c), coloured bars indicate the number of types for which there exist a given number of specific or non-specific lines.

Systematic names were generated based on the location of the soma in the central brain (Fig. 1c) (DNa: anterior dorsal, DNb: anterior ventral, DNc: pars intercerebralis, DNd: outside cell cluster on the anterior surface, DNg: gnathal ganglion, DNp: posterior surface of the brain, DNx: outside the brain) and assigned sequential numbers within these groups. We describe 6 new ‘DNa’ types, 3 new ‘DNb’ types, 2 new ‘DNd’ types, 86 new ‘DNg’ types, 48 new ‘DNp’ types, and 1 new ‘DNx’ type. DNs were also categorized by the VNC neuropils they innervated, following the convention from analysis of an EM VNC dataset (Cheong et al. 2024) (Fig. 1d) (fl: foreleg, ml: midleg, hl: hindleg, xl: multiple leg neuropils, nt: neck tectulum, wt: wing tectulum, ht: haltere tectulum, ut: upper tectulum, it: intermediate tectulum, lt: lower tectulum, ad: abdomen, xn: multiple neuropils).

Our initial survey of 98 DN types documented 78 “unique” types consisting of a single bilateral pair of neurons and 20 “population” types comprising groups of up to 30 cells with very similar morphology (Namiki et al. 2018). Among the 146 new DN types described in this study, the vast majority (122) consisted of unique bilateral pairs. Three types each consisted of a single neuron with symmetrical, bilaterally projecting neurites: DNg55, DNg66, and DNg114. The remaining 21 types formed larger populations of neurons with nearly identical morphology within a type. However, further investigation may reveal consistent differences in their connectivity, morphology, or function that merit splitting of types.

We have now described a total of 244 DN types from LM images—approximately 50% of the types known from EM datasets (Fig. 1e,f). The morphologies of these neurons are shown in Figures 2 through 26. (For three-dimensional stacks, see Supplementary Table 2 for a list of publicly available slides.) Nearly all of these have been matched to neurons in existing EM datasets (FAFB, MANC, FANC) by Stuerner et al. (2024).

**Figures 2–26.**
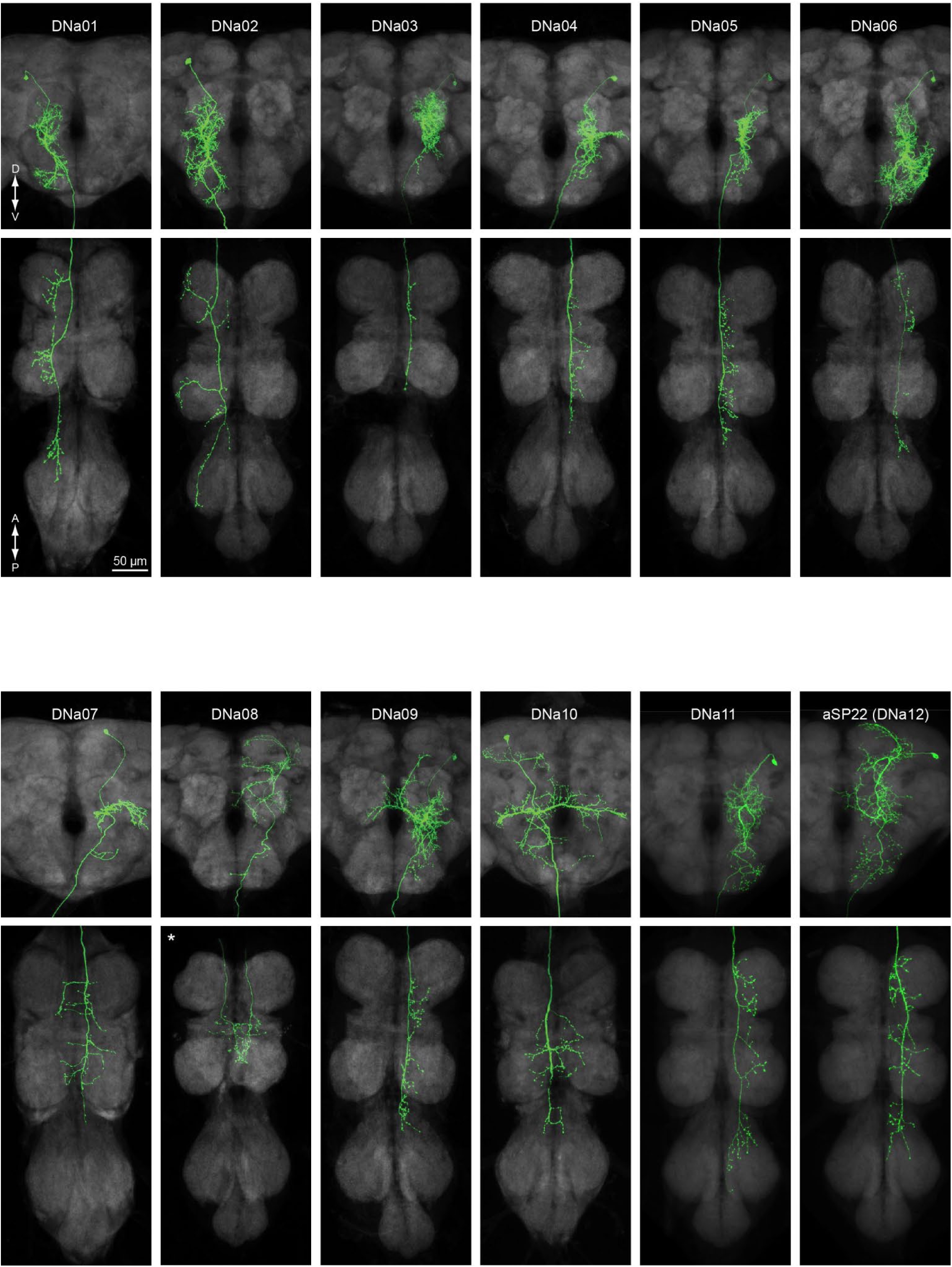
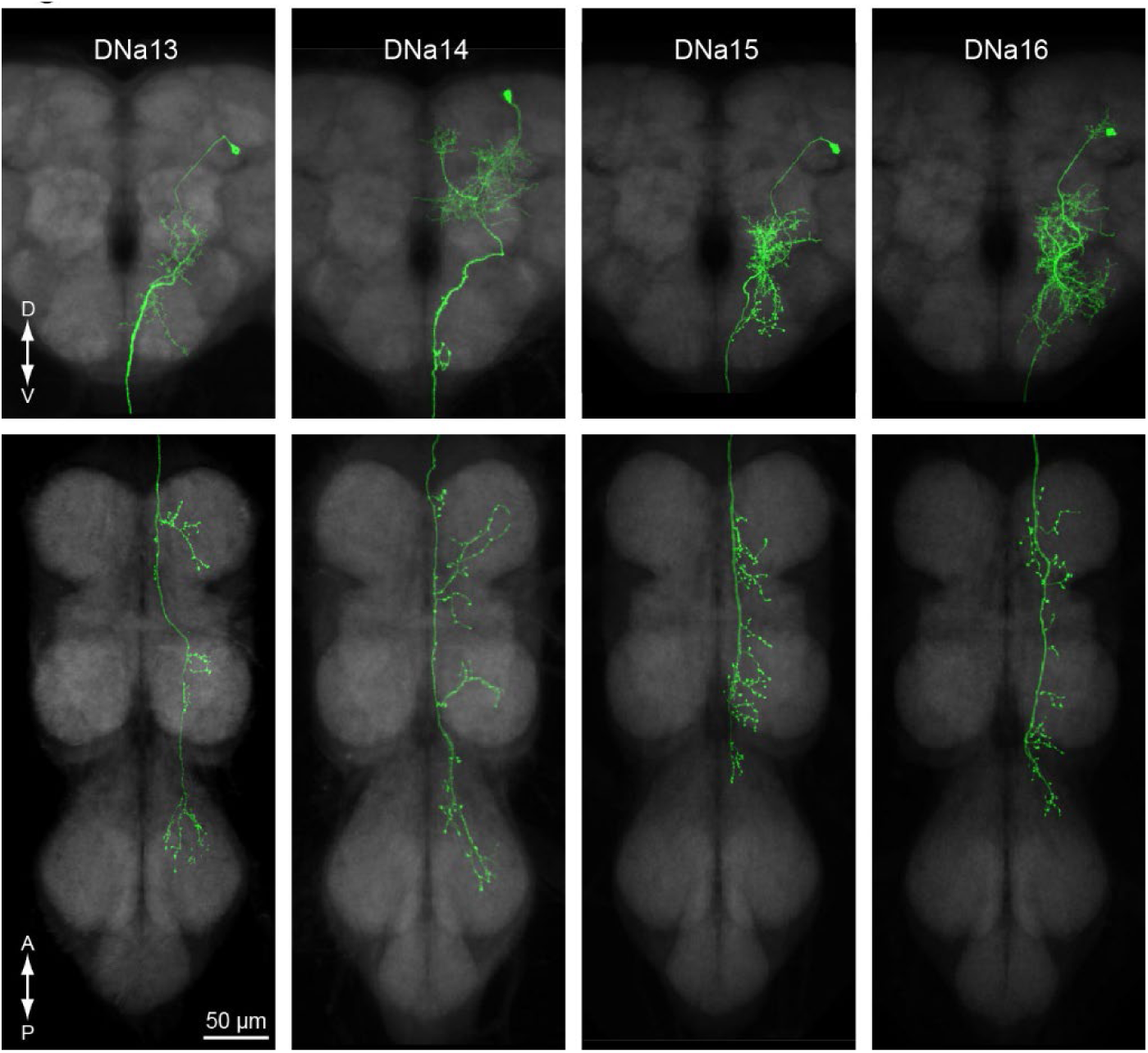
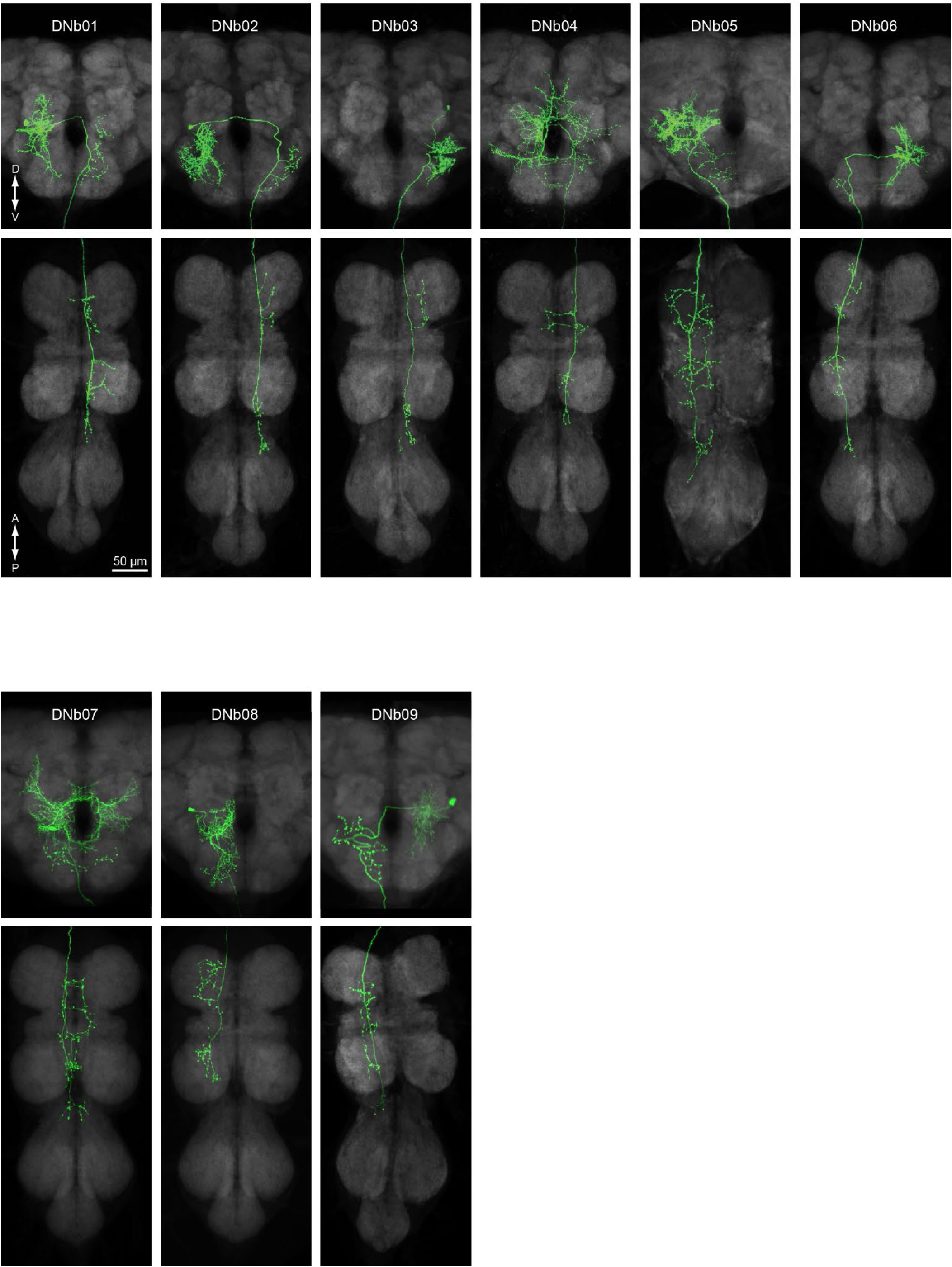
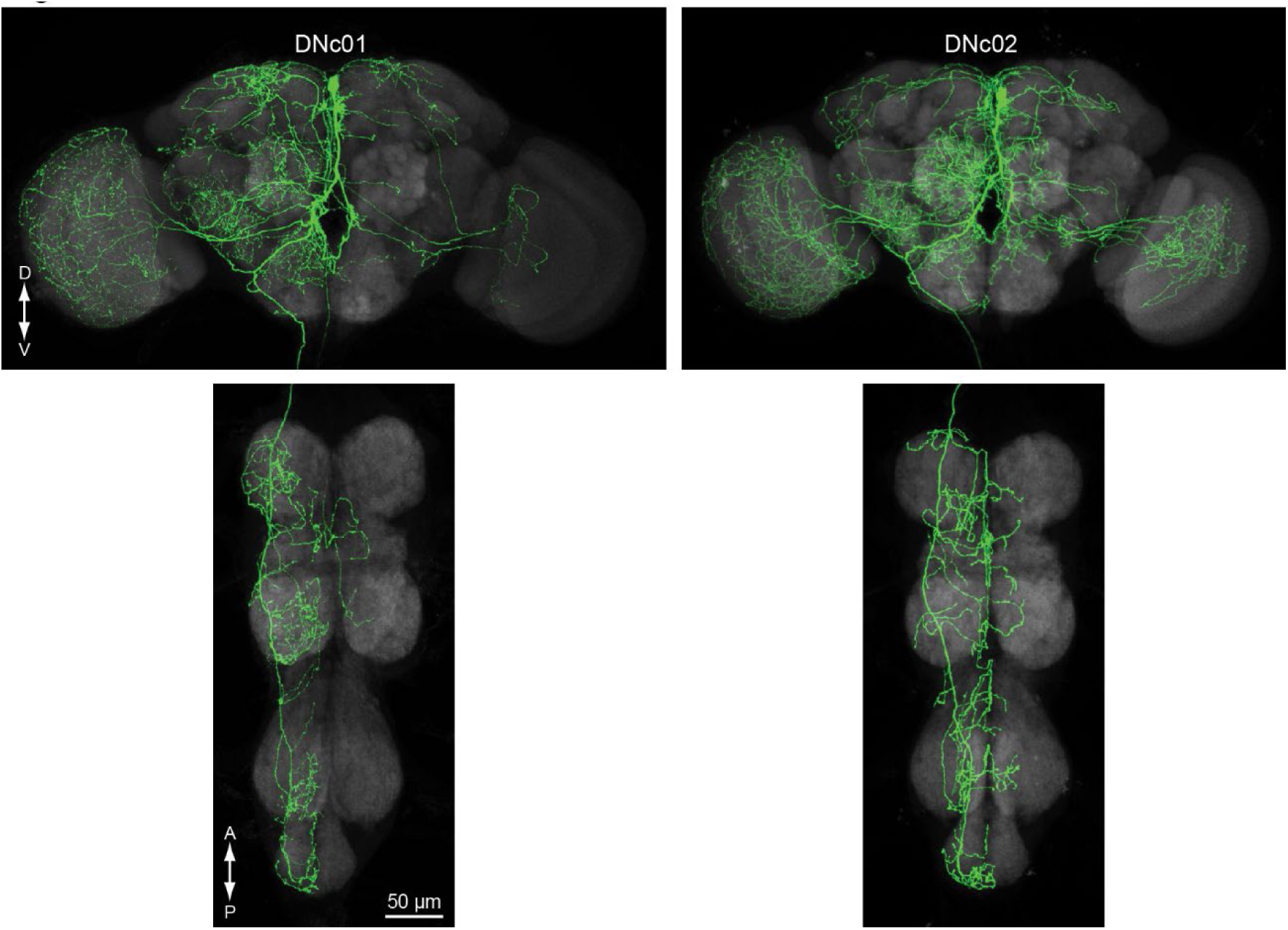
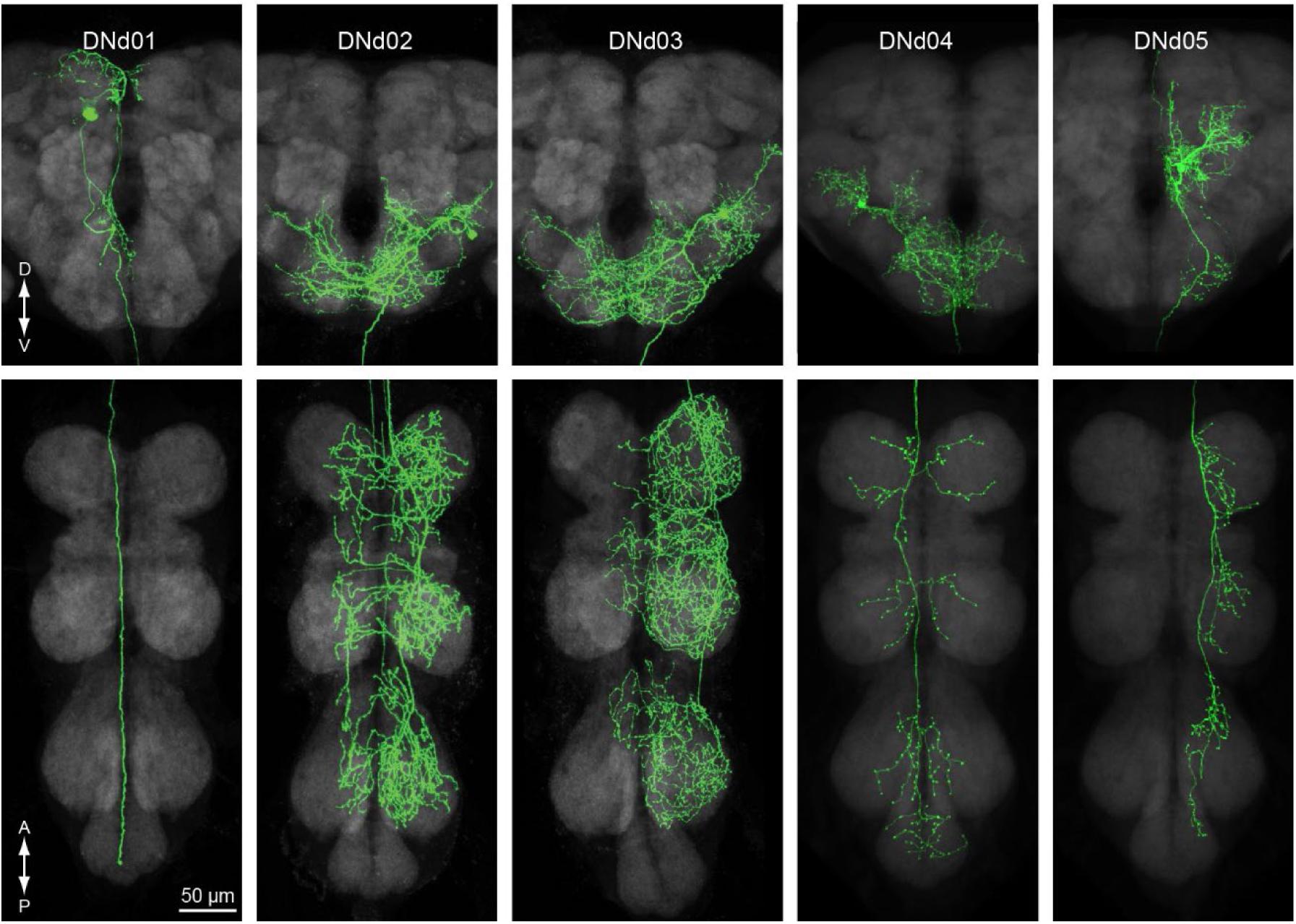
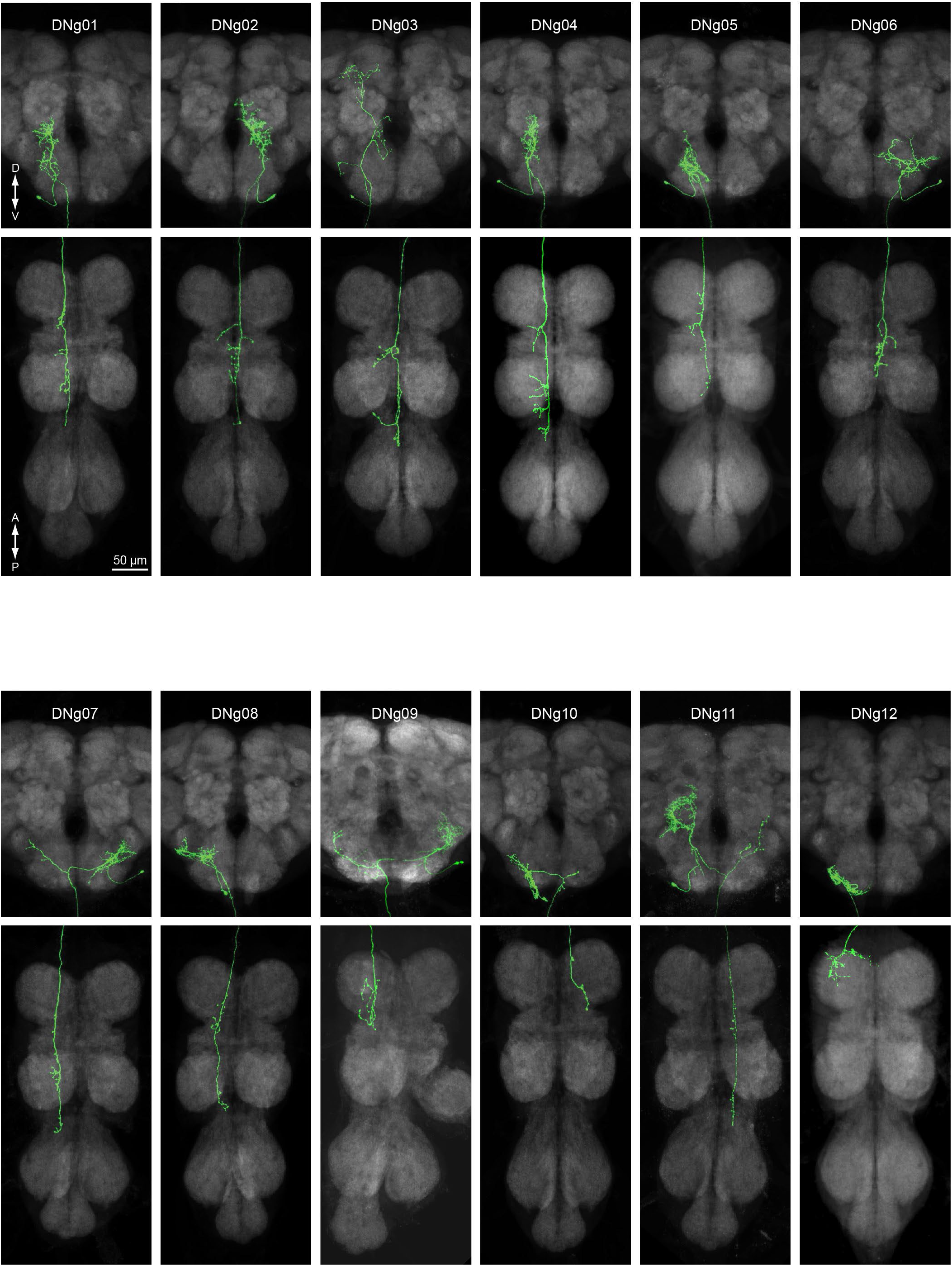
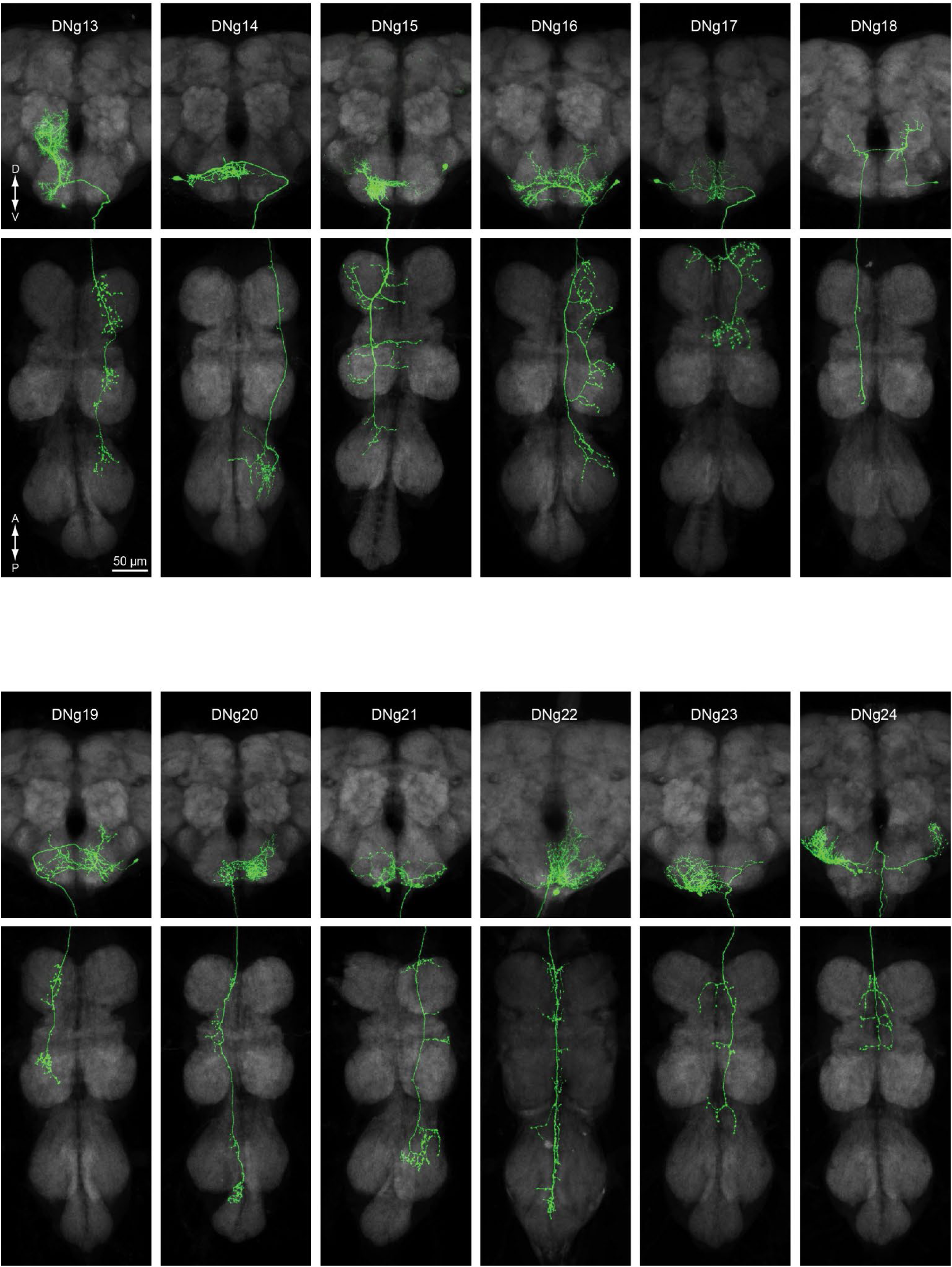
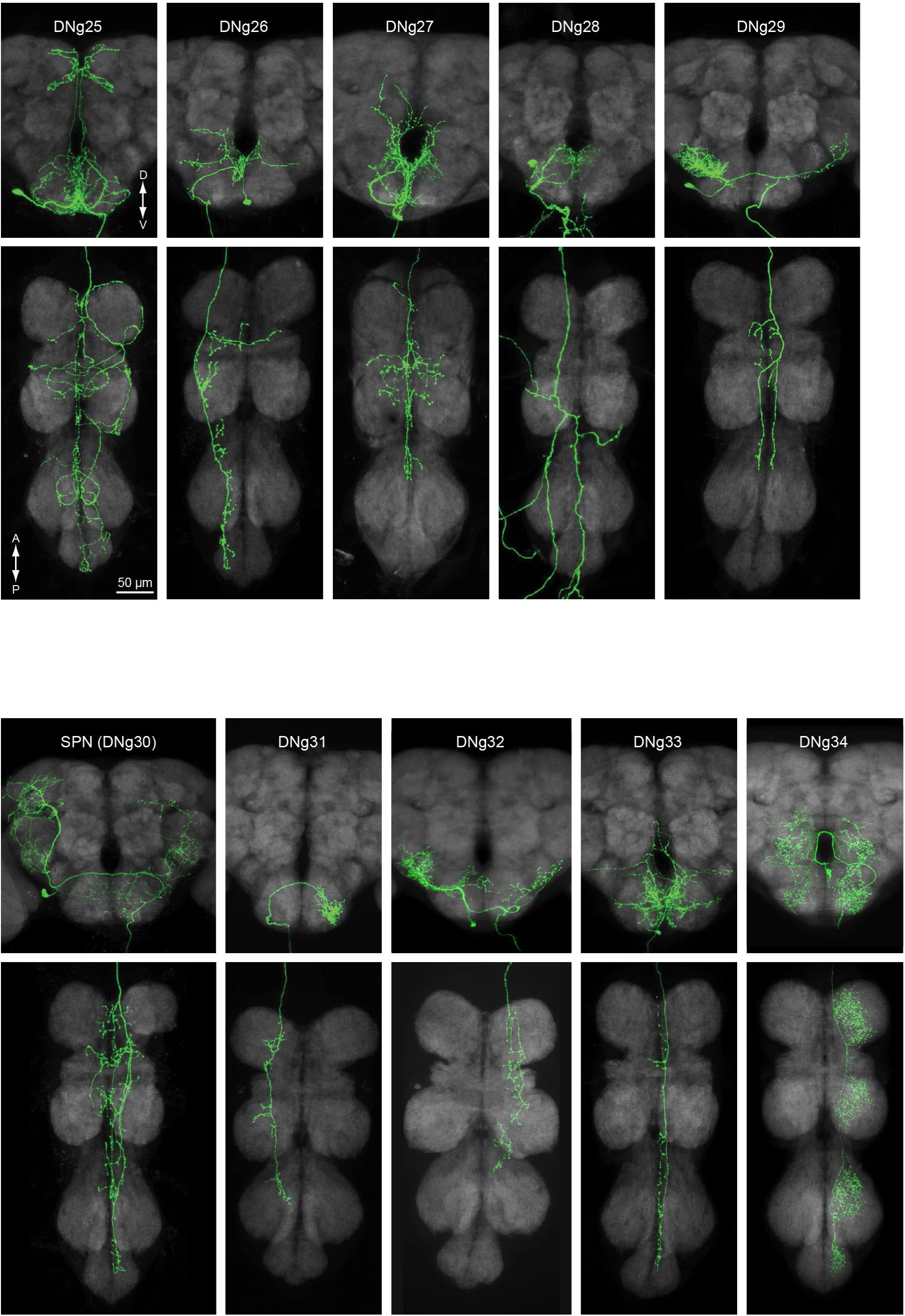
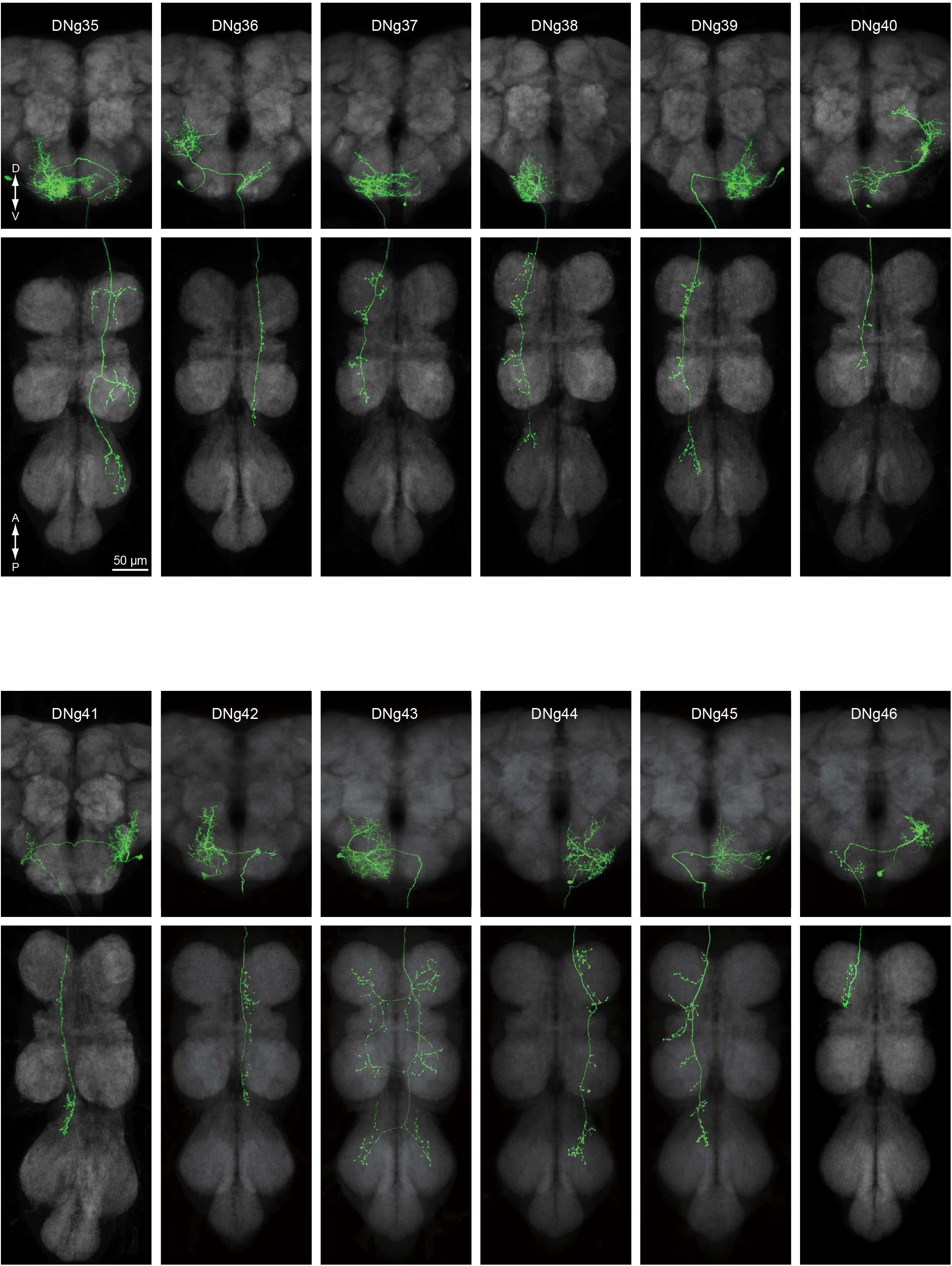
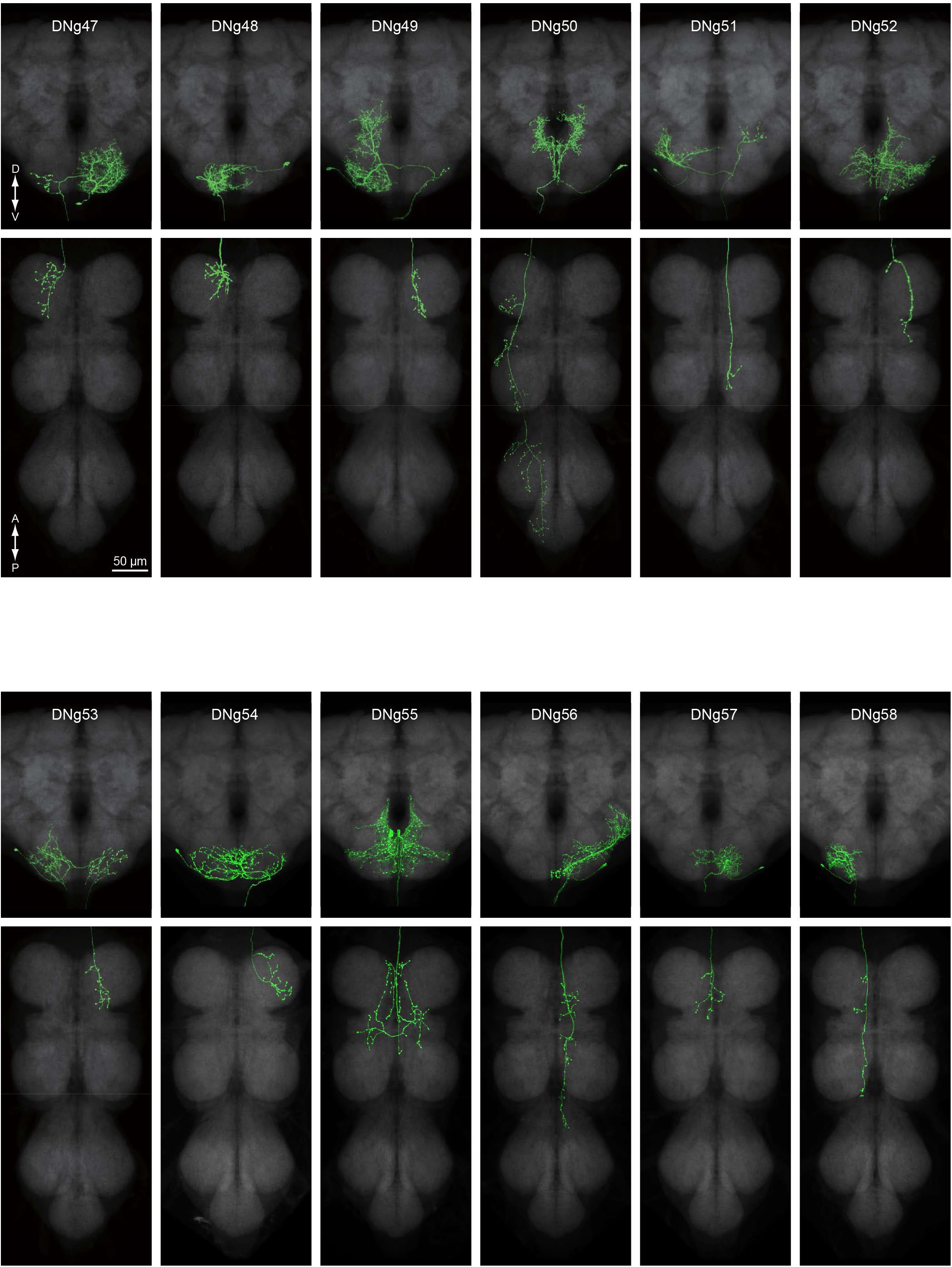
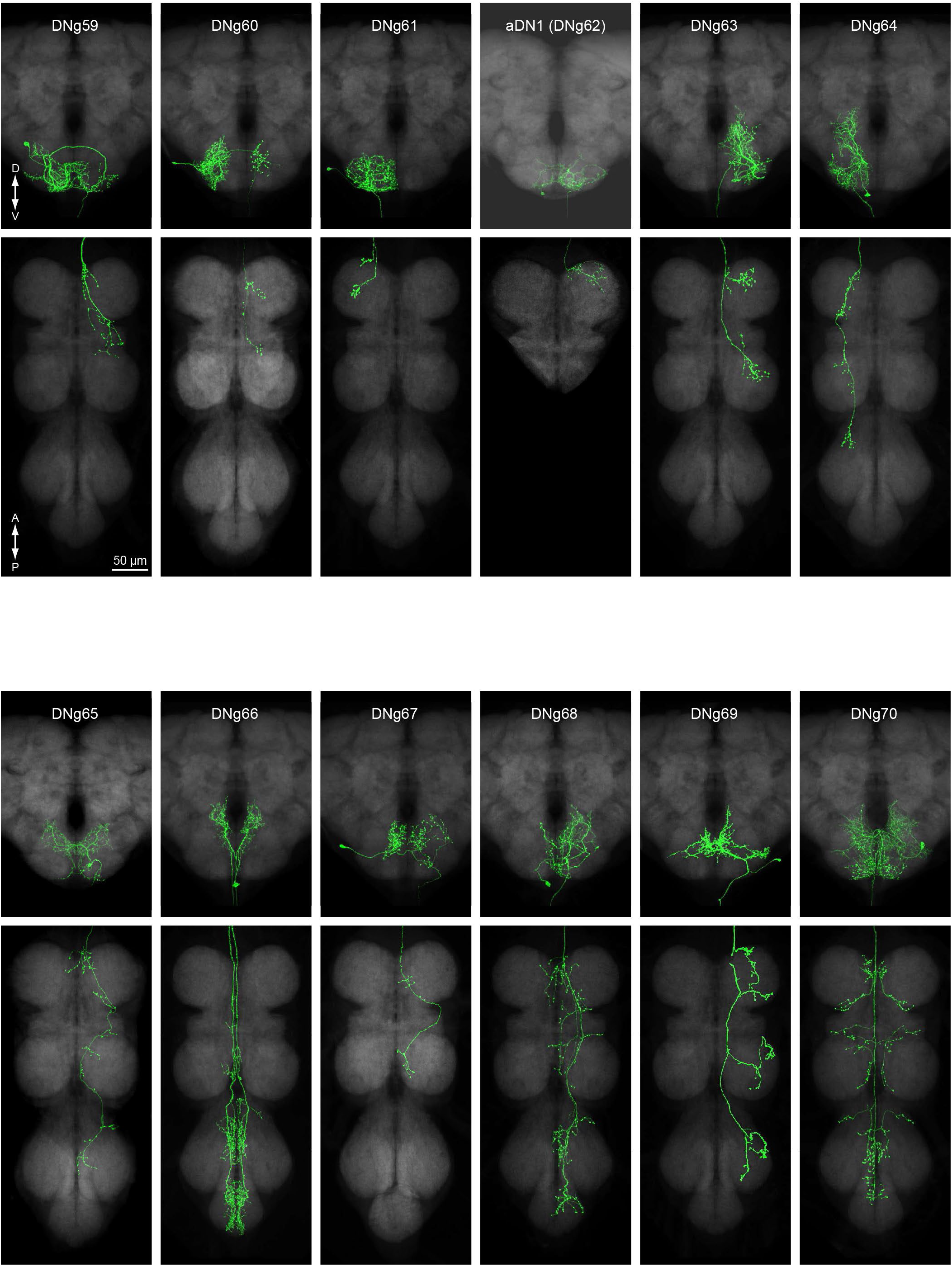
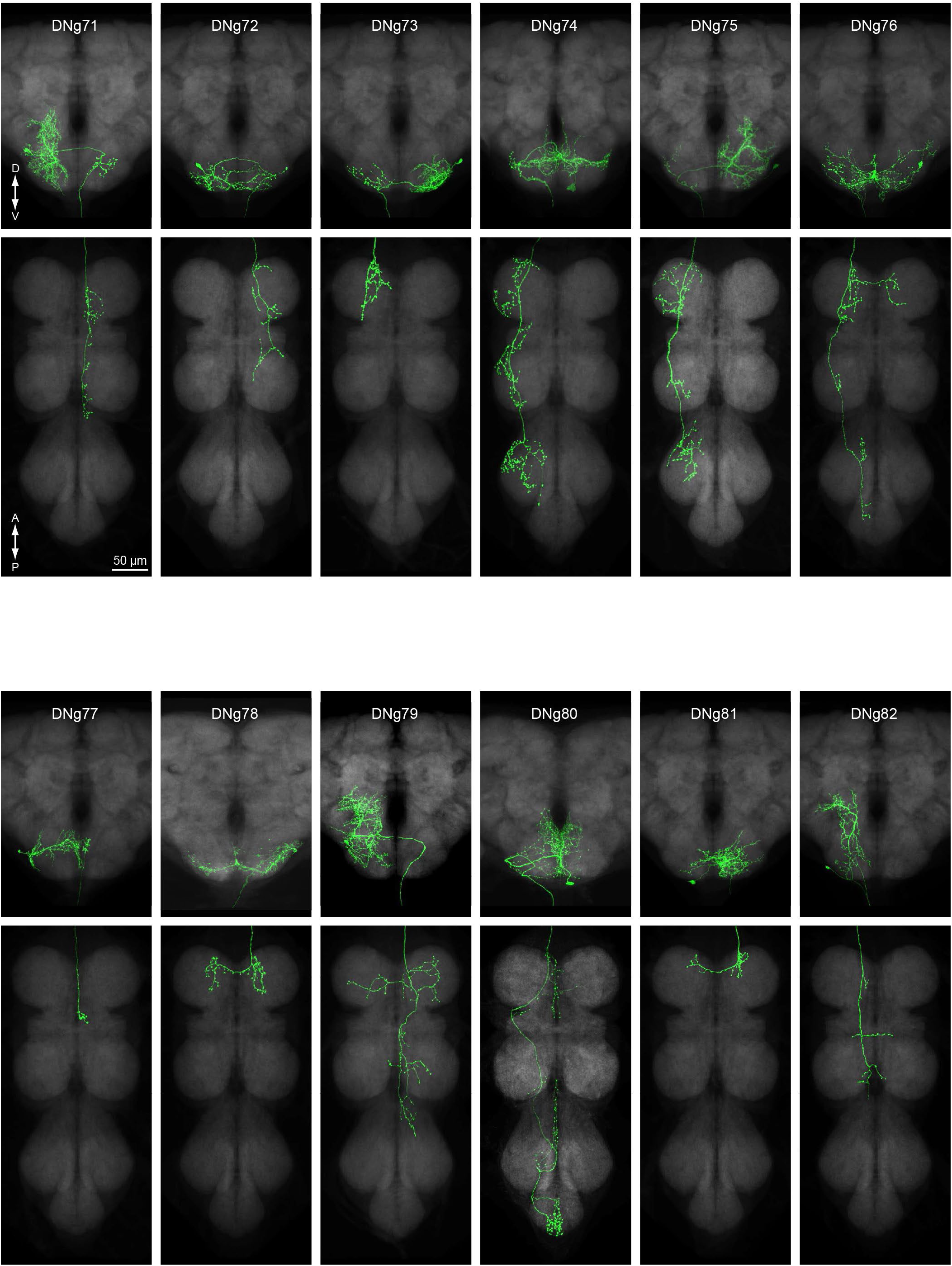
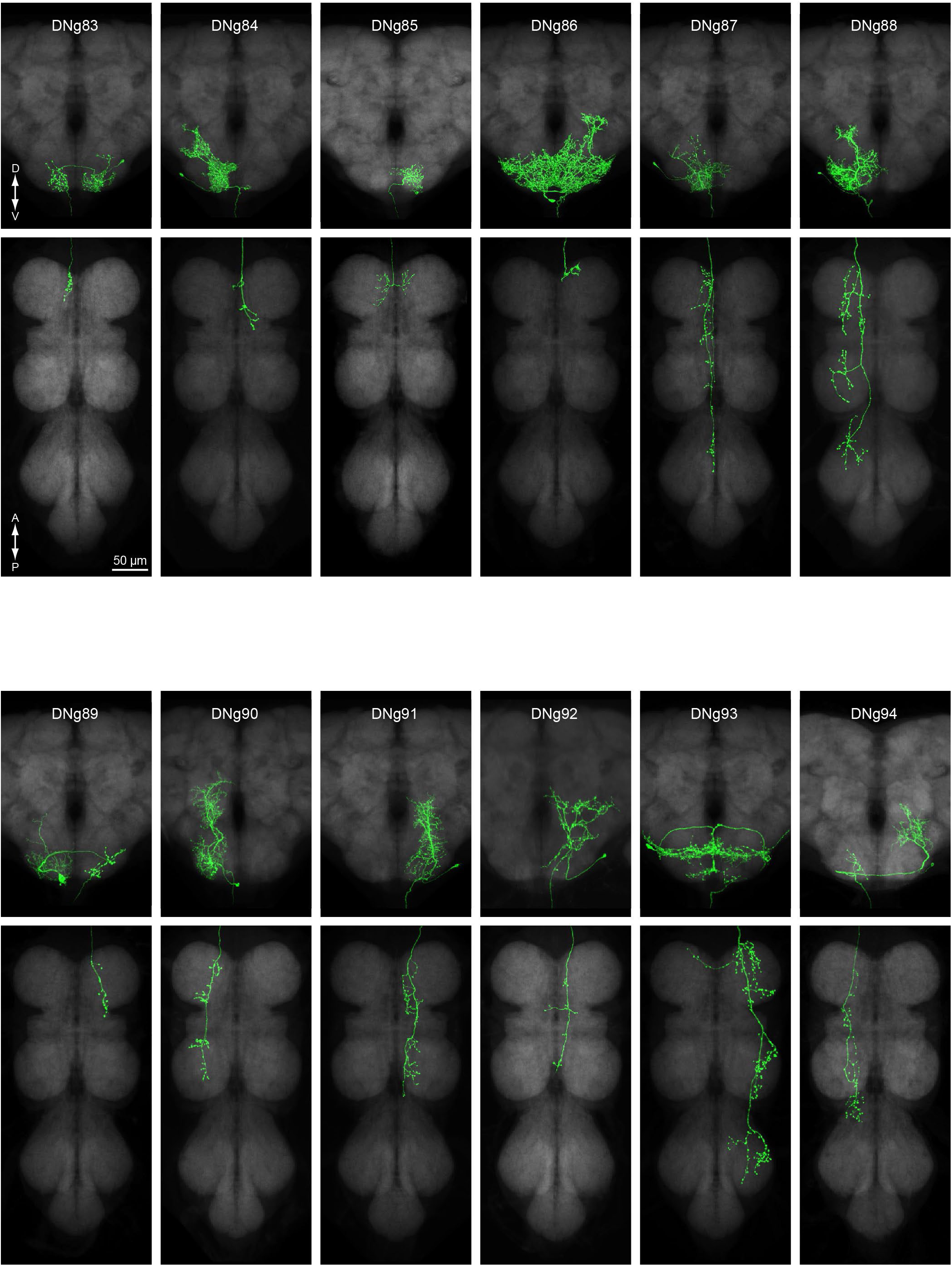
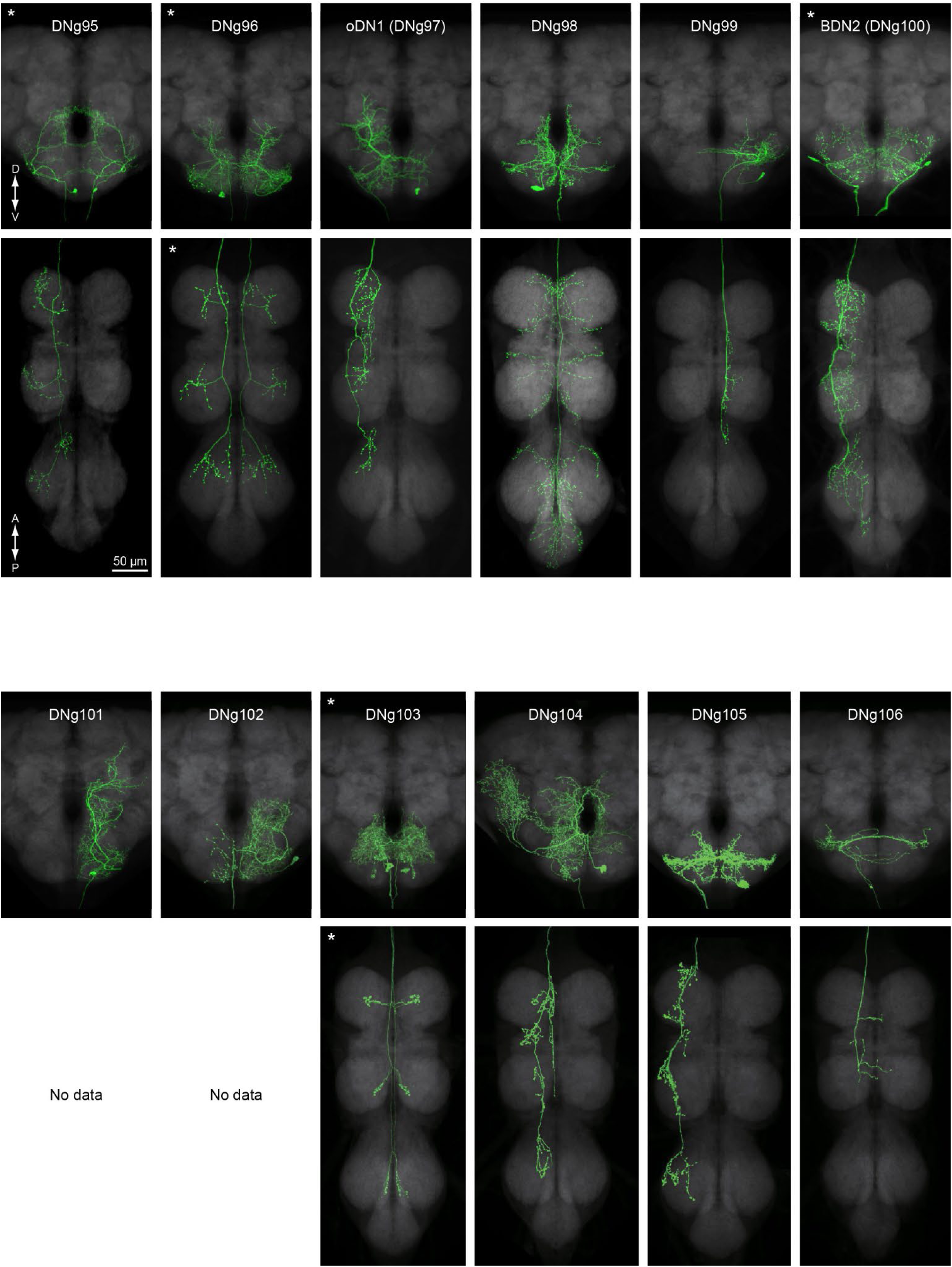
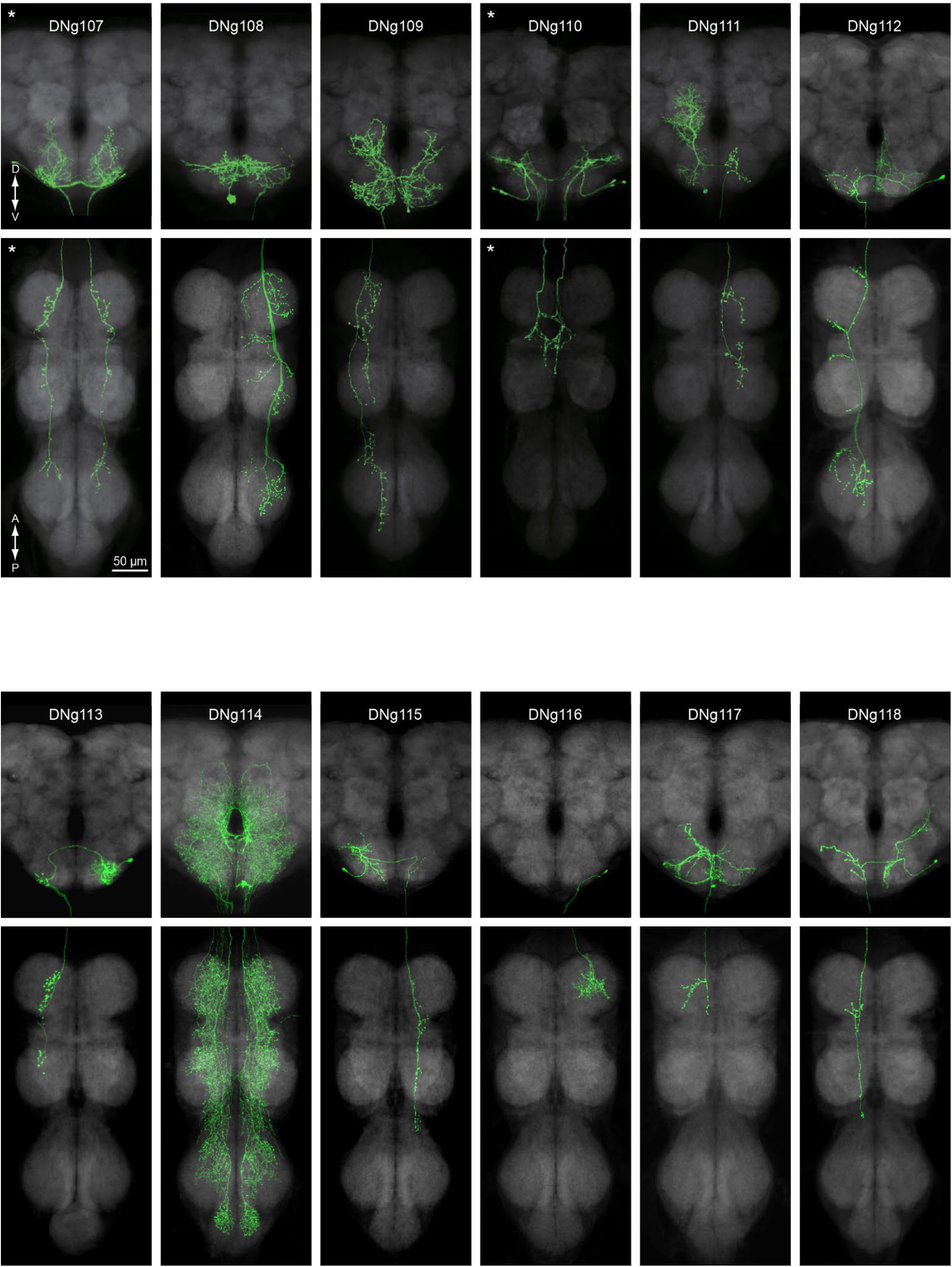
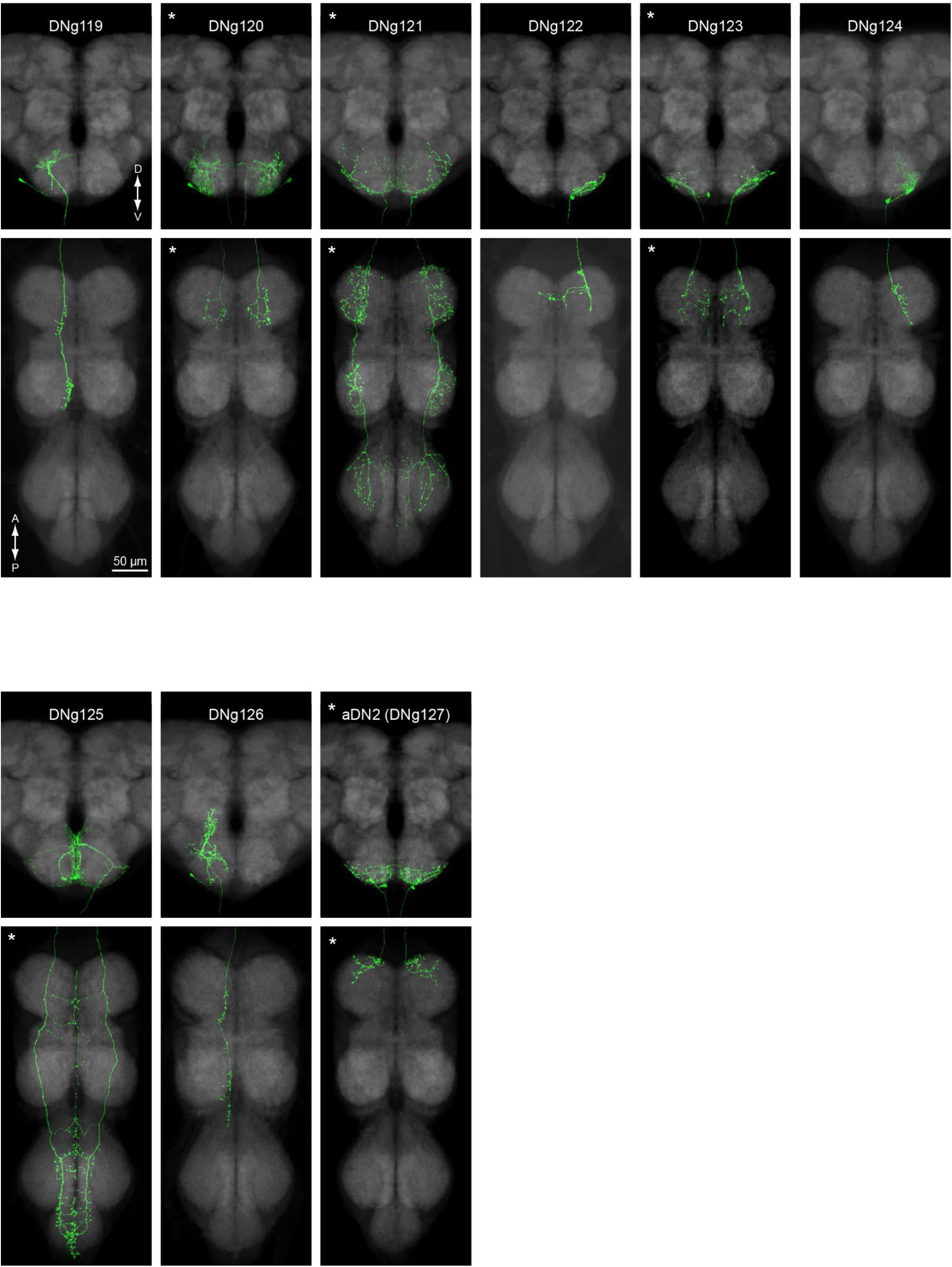
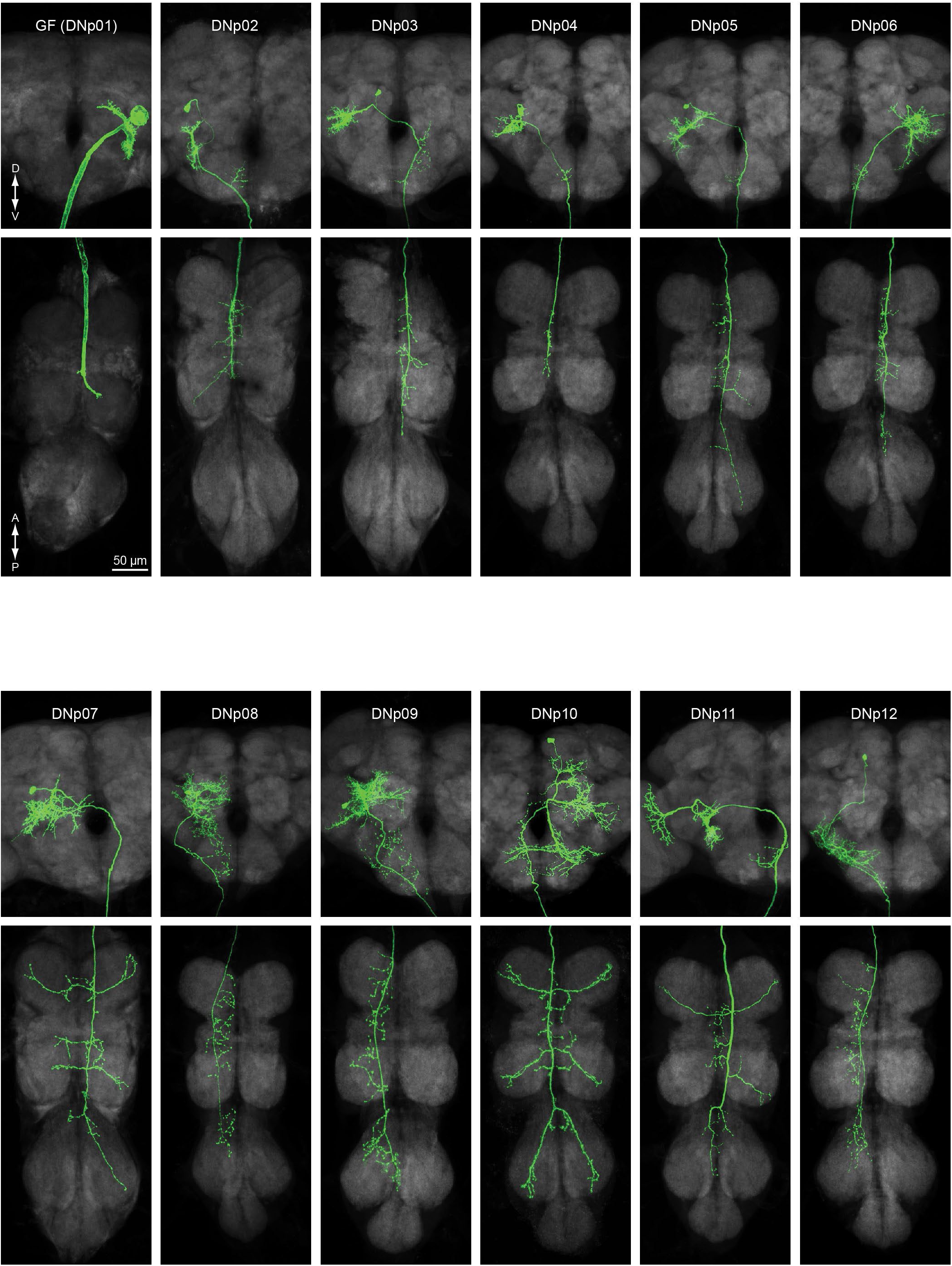
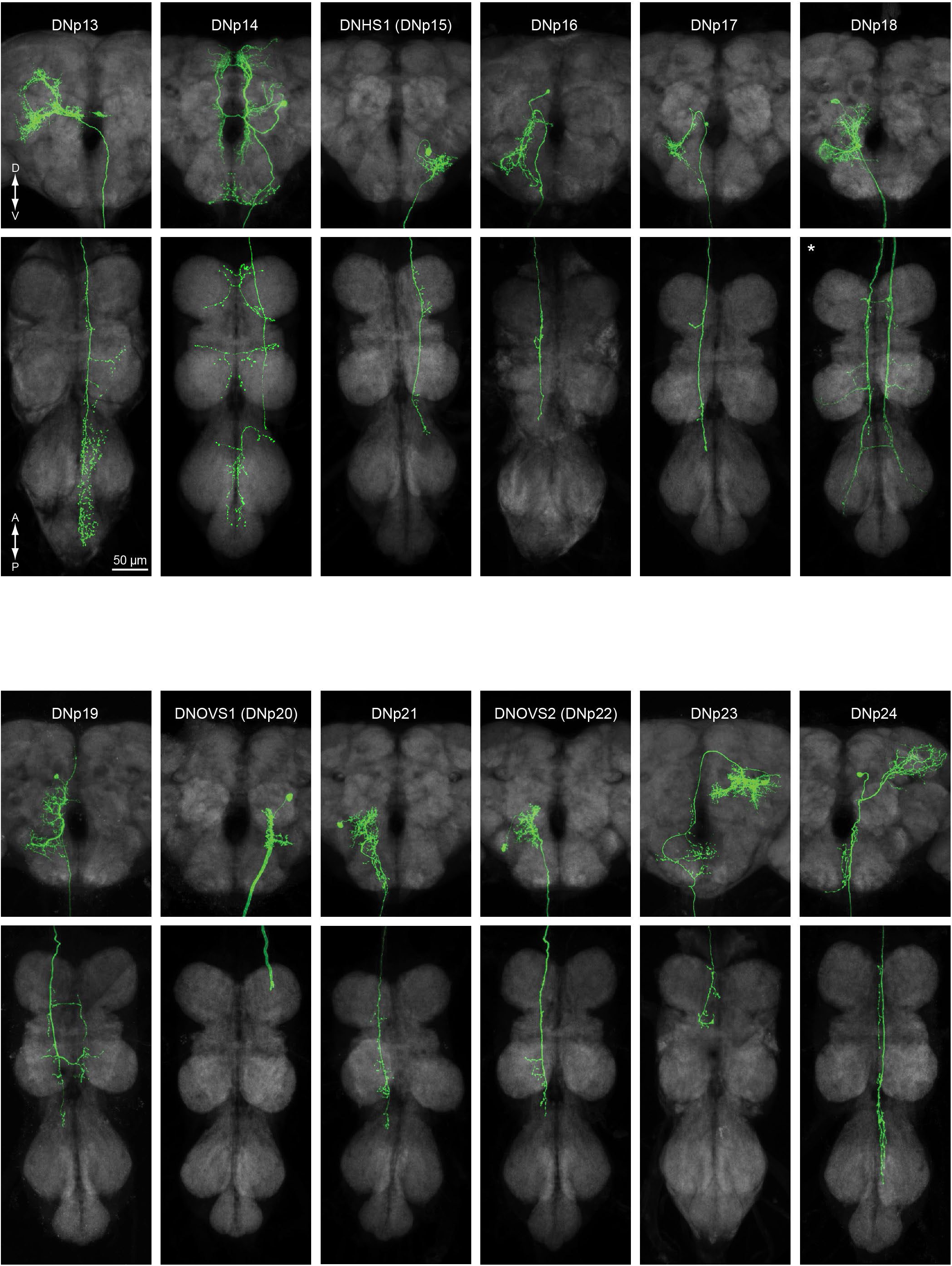
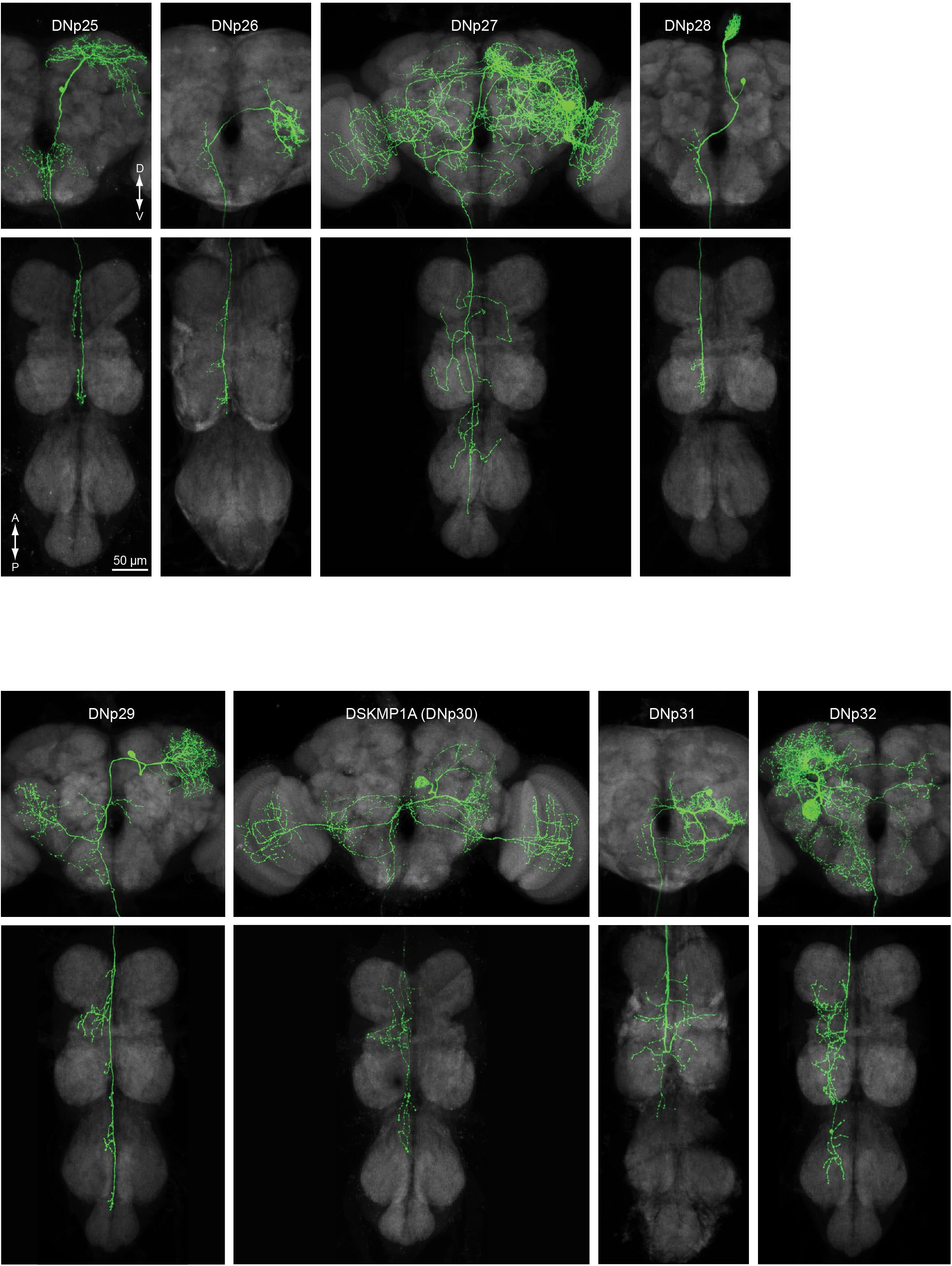
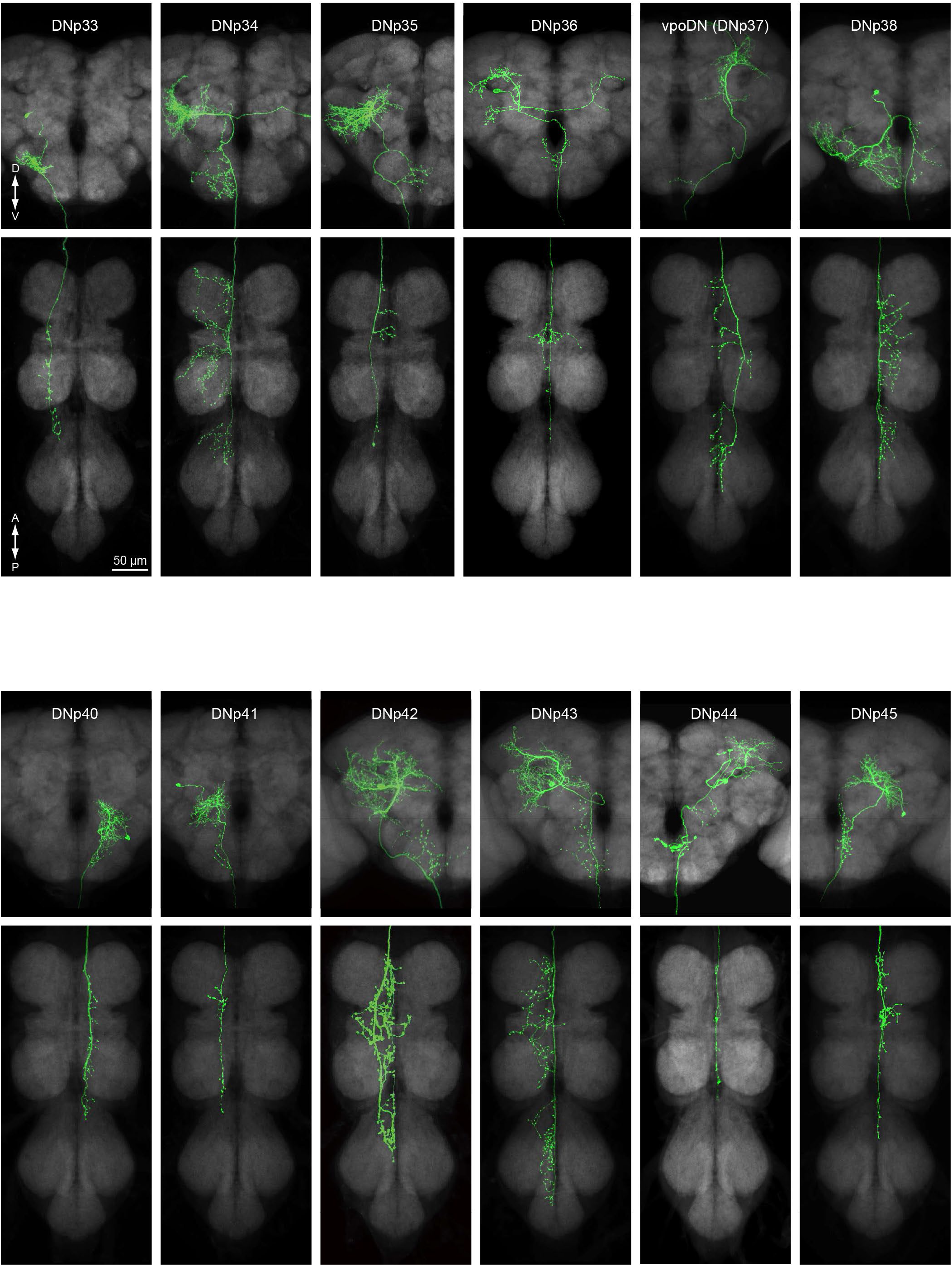
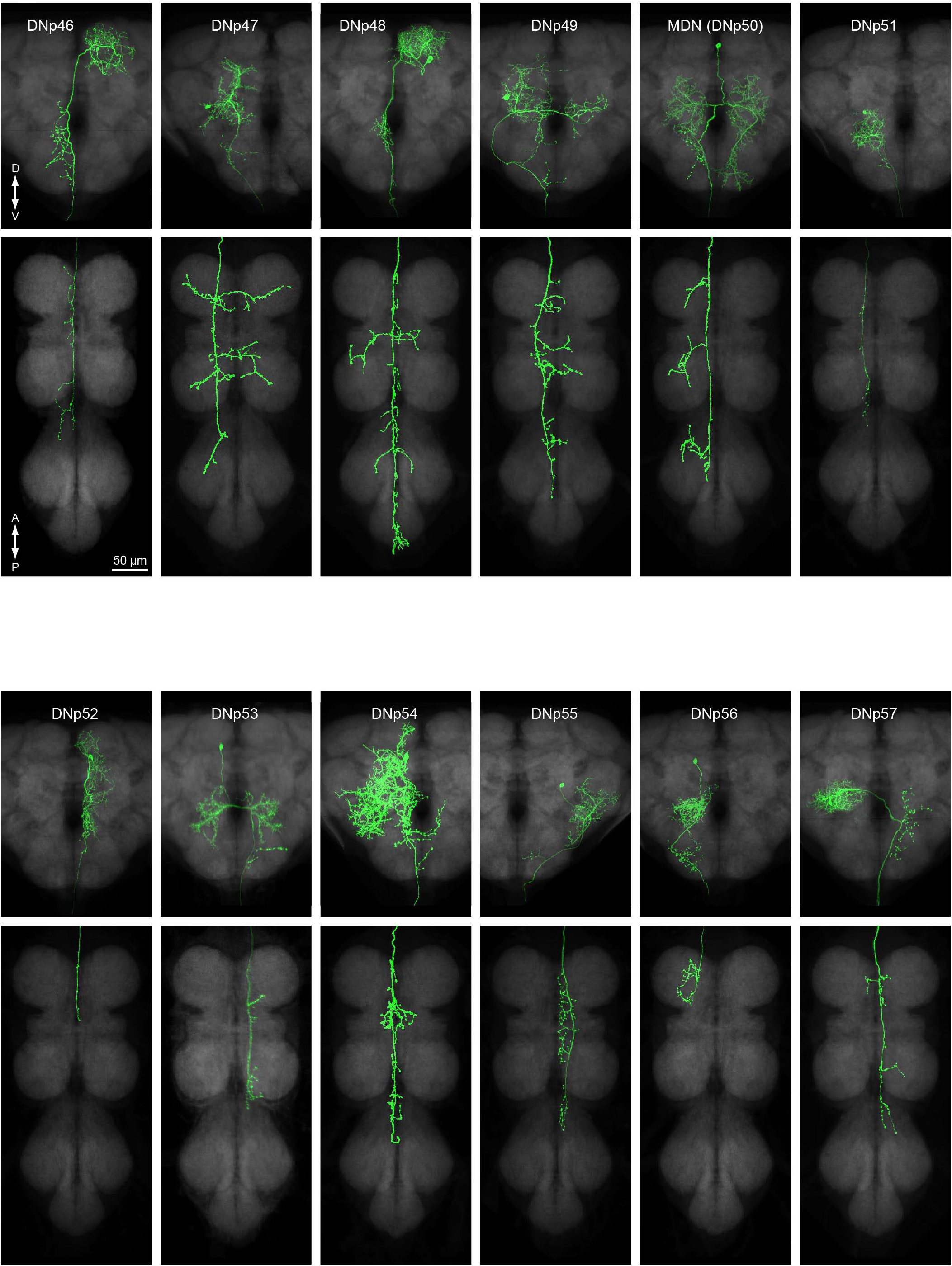
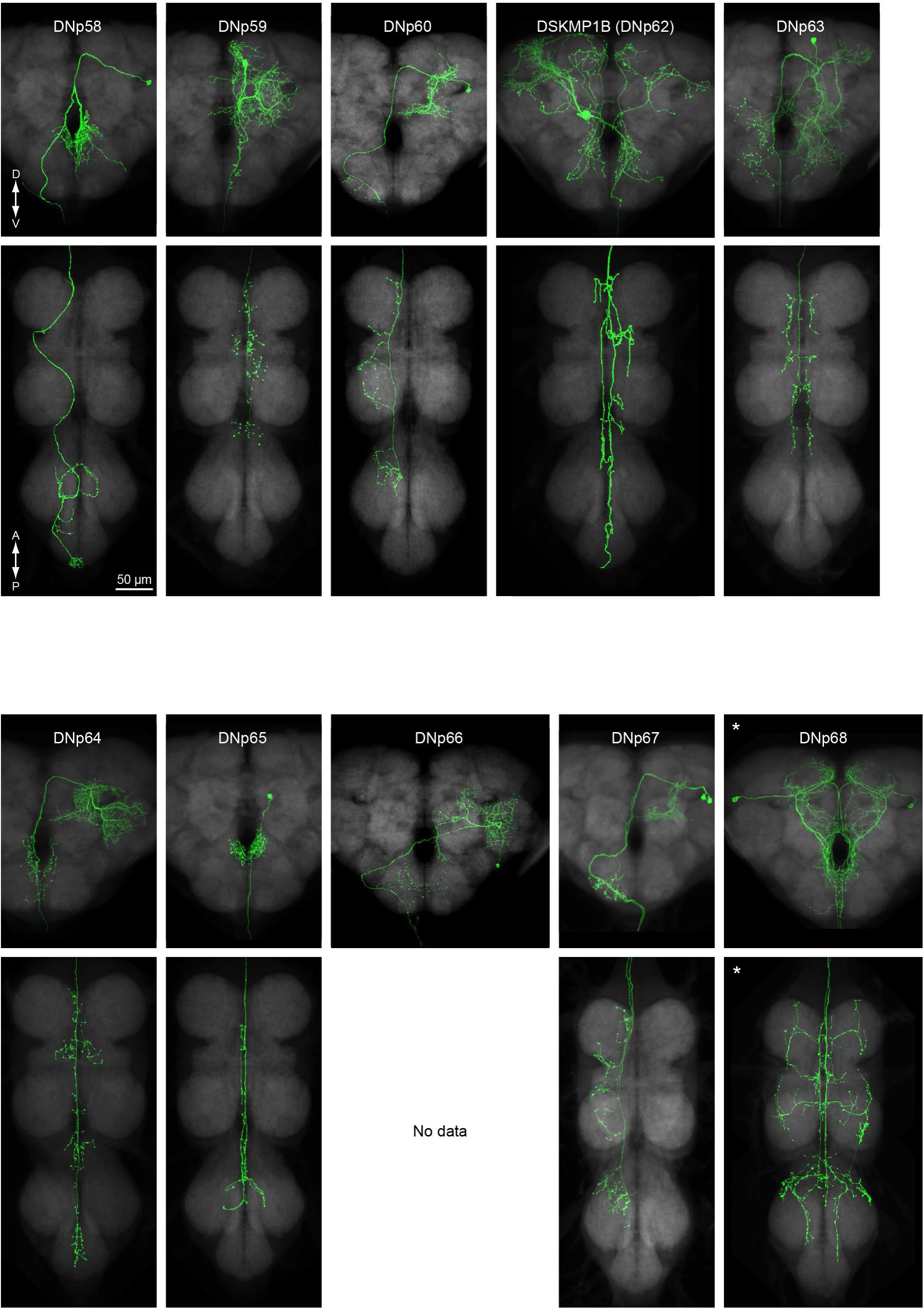
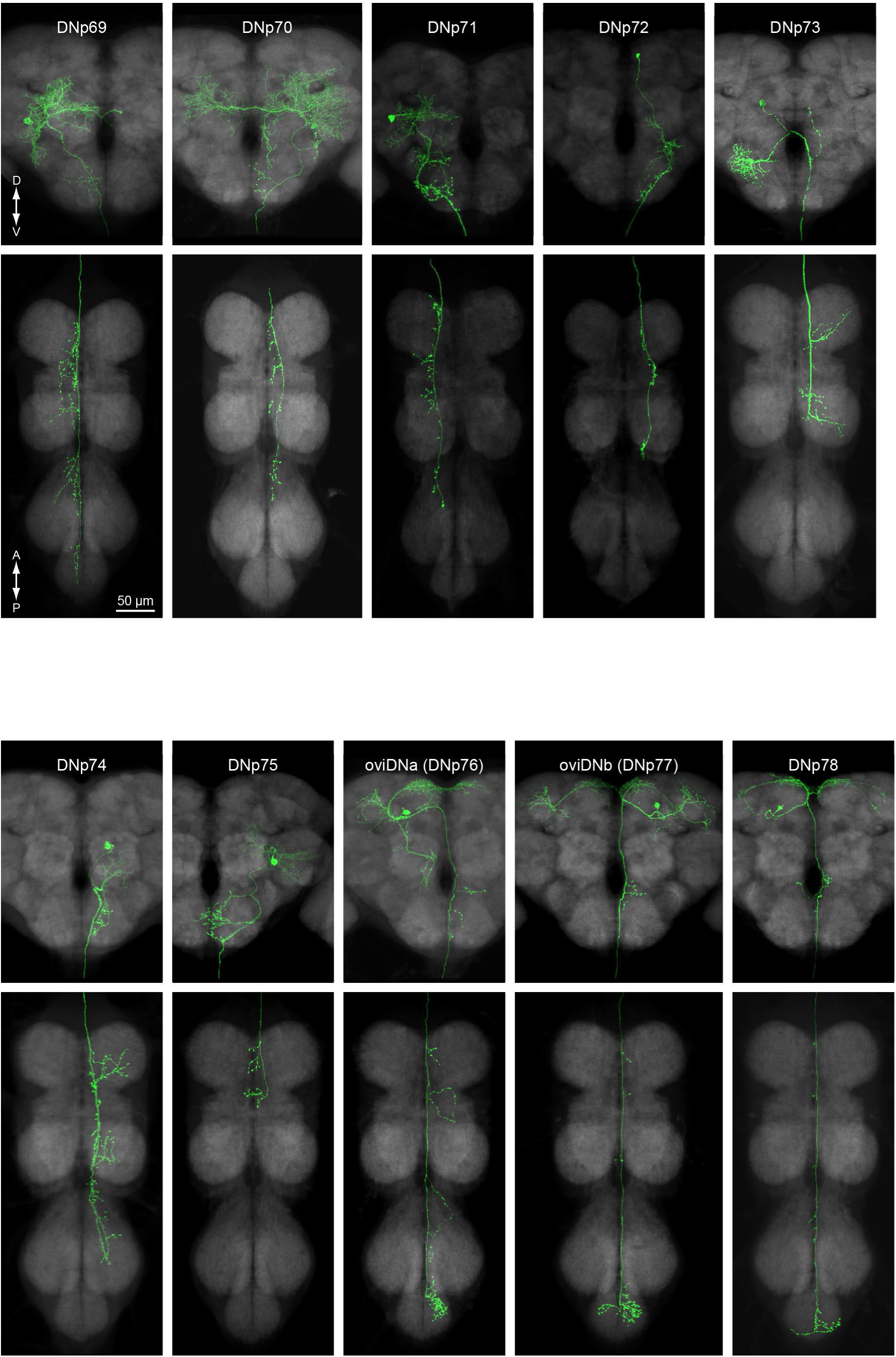
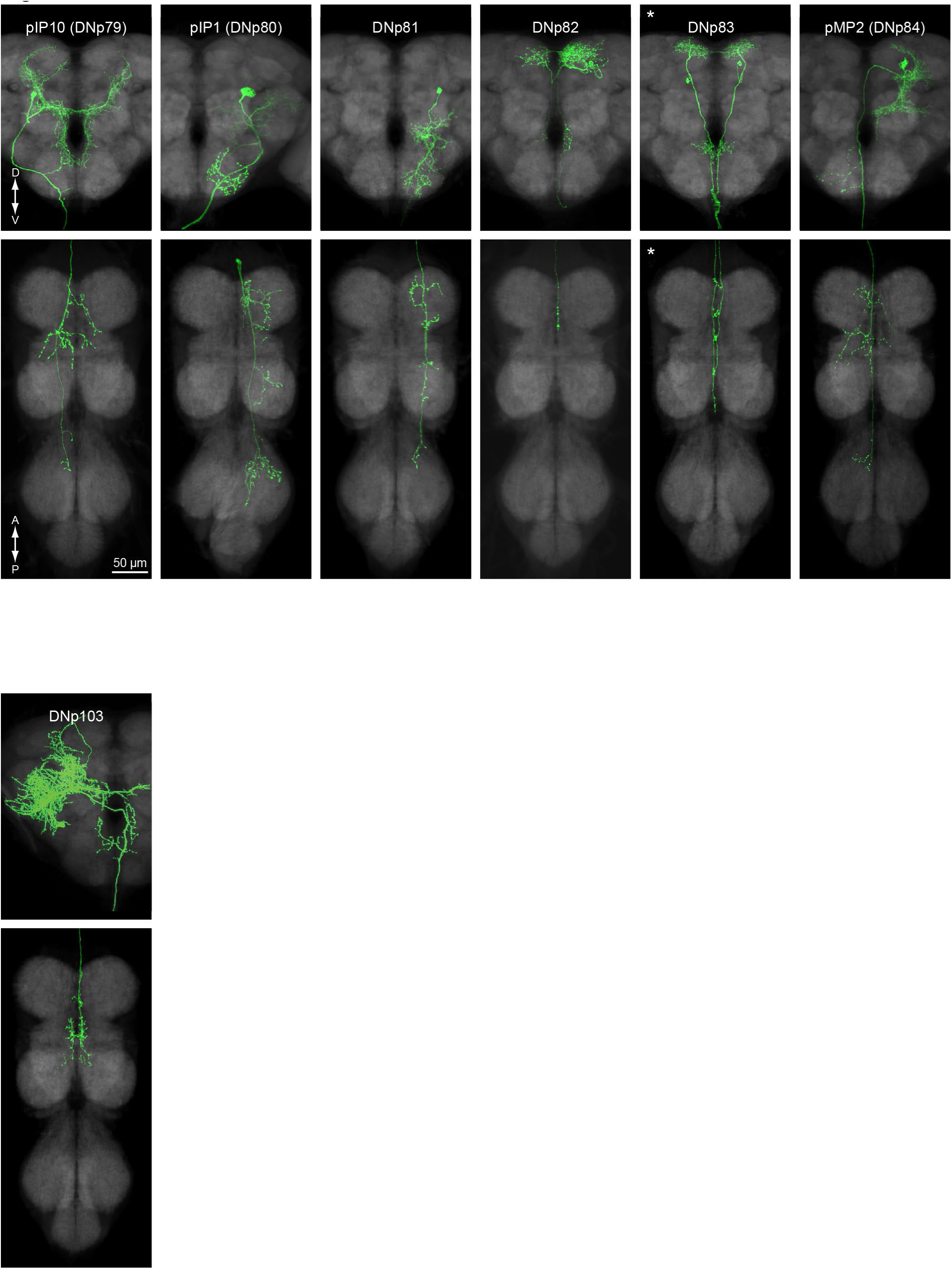
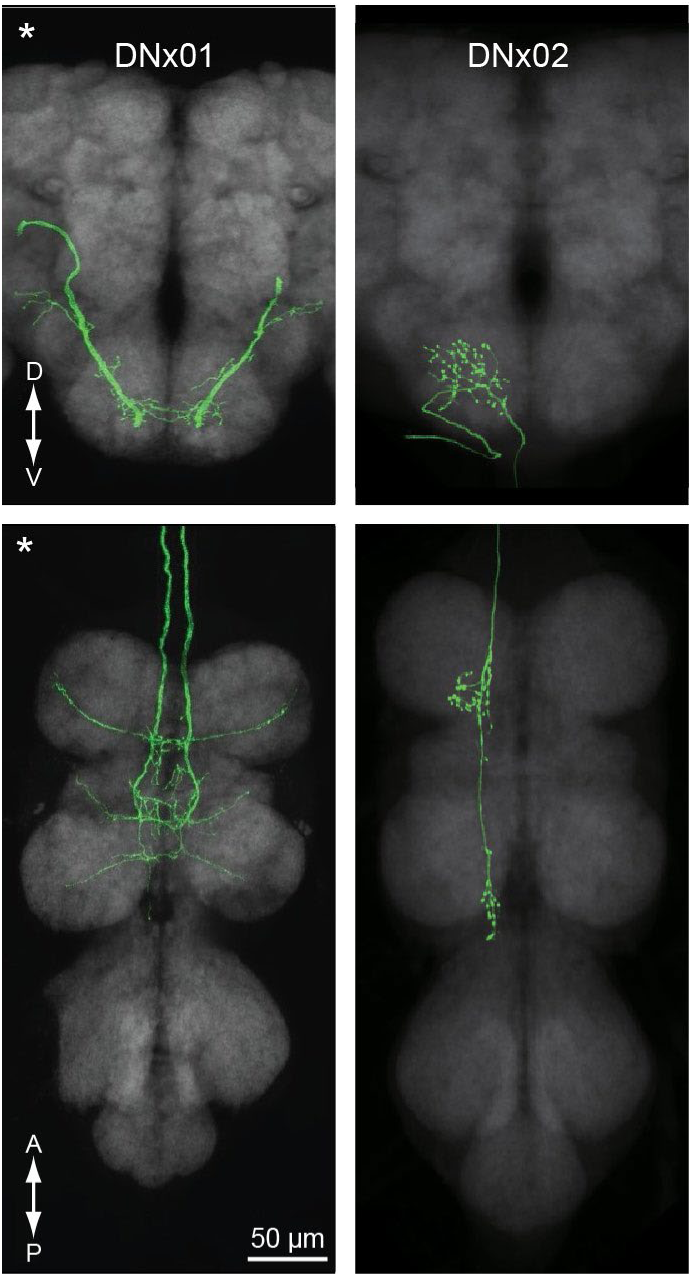
Morphology of descending neurons identified in Namiki et al. (2018) and in the present study. Images of neurons have been manually segmented (see Methods for details). Asterisks indicate images showing two or more neurons of the same type (generally both instances of a bilateral pair).

### Updates to 2018 eLife paper

While characterizing new DN types, we discovered subtle but consistent morphological variation among neurons within four DN types we previously defined in Namiki et al. (2018): DNb01, DNg05, DNg16, and DNp13. These variants have now been defined as new types. In each case that follows, the first name of each pair formerly encompassed both variants, but both names now describe distinct types: DNb01/DNb09, DNg05/DNg126, DNg16/DNg69, and DNp13/DNp62.

In addition, we note that the morphologies of several DN types described in Namiki et al. (2018) were atypical, due to developmental abnormalities or misidentifications in the image examples used to describe them. Figures 2 through 26 include revised images for all these cases. Our ground truth was the split-Gal4 lines annotated with a given DN type (if available) or the images found in the figure supplements to Figure 2 in Namiki et al. (2018). A full list of the revisions may be found in Supplementary Table 3.

We also update annotations for split-Gal4 lines produced by Namiki et al. (2018), generally adding unannotated types and in a few cases removing annotations for types where we no longer see convincing evidence that they are labelled (see Supplementary Table 4).

### New DN split-Gal4 lines

Our next goal was to create new cell-type-specific driver lines for as many DN types as possible using the intersectional split-Gal4 technique (Luan et al. 2006). Briefly, this method involves choosing pairs of enhancers whose expression pattern overlaps in the target cell type and using them to drive expression of the two halves of the GAL4 transcription factor, such that a functional GAL4 protein is expressed only in cell types labelled by both enhancers. Using the Janelia FlyLight image database (Meissner et al. 2023) and “split half” hemidriver lines (Tirian and Dickson 2017; Dionne et al. 2018), we identified candidate enhancer pairs and crossed them to produce 500 new split-Gal4 driver lines that sparsely label DNs (Supplementary File 1, Supplementary Table 5). 131 of these lines were previously released with tentative annotations as part of Janelia’s “omnibus” list of top cell-type-specific lines (Meissner et al. 2025), but are now more thoroughly curated with updated annotations.

We gave each line an overall quality score from ‘A’ (best) to ‘C’ (worst) based on their specificity and consistency in labelling the annotated DN type(s). We used stringent criteria for lines given an ‘A’ score (see Methods for details), stricter than we used previously (Namiki et al. 2018). For these lines, optogenetic activation induced by illuminating the entire animal is expected to reliably activate the target neuron(s) with virtually no off-target effects in the rest of the central nervous system. For many other applications such as imaging or targeted optogenetic illumination, ‘B’ or even ‘C’ lines should be more than suitable. Examples of lines given different scores are shown in Supplementary Figure 1. In total, we scored 97 lines ‘A’, 325 ‘B’, and 78 ‘C’. Supplementary Table 1 lists lines by DN type and quality score.

### Compilation of previously published DN split-Gal4 lines

Many other research groups have created split-Gal4 driver lines that label descending neurons. We have compiled a list of the 128 lines we found in the literature and collated it with our own collection (Supplementary Table 5; lines also included in Supplementary Table 1). While our search was by no means exhaustive, our list now represents the most comprehensive collection to date of DN driver lines, comprising 806 lines targeting 190 DN types.

### Line availability

382 of these 806 lines are available at the Bloomington *Drosophila* Stock Center at the time of publication (Supplementary Table 5). New lines released in this paper and those from Namiki et al. (2018) will be maintained until at least December 31, 2026 by the authors and are available upon request. Readers may also reconstruct the lines using split hemidrivers, which are currently maintained at the Bloomington *Drosophila* Stock Center.

## Discussion

Genetic access is often a key prerequisite to understanding the function of a target cell type. Here, we provide a significantly expanded catalogue of driver lines that enable genetic access to DNs in *Drosophila melanogaster*. DNs are relatively few in number yet are the sole conduits of nervous signals from the brain to the ventral nerve cord and its downstream motor circuits. Thus, DNs are well poised to provide insight into the neural control of behaviour.

Optogenetic activation of DNs can be especially informative, since these neurons transmit information to motor neurons via a relatively short path, in some cases forming direct synapses. Indeed, activation of single bilateral DN pairs has been shown to drive whole coordinated motor programs such as forwards walking (Bidaye et al. 2020; Rayshubskiy et al. 2024), backwards walking (Bidaye et al. 2014), takeoff (von Reyn et al. 2014), landing (Ache et al. 2019), courtship song (Ding et al. 2019; Lillvis et al. 2024), and grooming (Hampel et al. 2015; Guo et al. 2022), reinforcing the idea that DNs convey high-level commands driving complex behaviours rather than lower-level coordination of individual muscle movements. There is also mounting evidence that subsets of DNs coordinate behaviour as a population. Namiki et al. (2018) noted that the number of individual neurons in a DN type, as defined by neurons with nearly identical morphology, typically ranges from 2 (a single left–right pair) to 30. Those found to activate coordinated motor programs thus far have mostly been types with only a single left– right pair, or “unique” DNs. In contrast, a “population” DN type with ∼13 neurons per hemisphere represents the gain of the fly’s wingstroke as the fraction of the population activated (Namiki et al. 2022). Further supporting population representation, activation of different DN types can evoke similar behaviours (Cande et al. 2018), and calcium-imaging experiments reveal that many DNs are active during various behaviours (Aymanns et al. 2022). Activation of a single DN type has also been shown to directly recruit other DNs through collateral connections (Braun et al. 2024). These observations suggest that many DNs may not act as “command neurons” that uniquely signal a specific behaviour, but instead encode activity as a population. In any case, results from optogenetic experiments must be interpreted with caution. While optogenetic activation of single types may indicate involvement in a particular behaviour, it is unlikely to recapitulate natural endogenous activation.

Our driver lines also enable recording of neural activity from the same identified DNs across multiple animals. Imaging using genetically encoded calcium indicators has revealed important insights into DN activity (Chen et al. 2018; Namiki et al. 2022; Aymanns et al. 2022). However, several studies have shown that precise spike timing is important for DN signalling (von Reyn et al. 2014; Rayshubskiy et al. 2024). The relatively low temporal resolution of calcium imaging may therefore obscure important biological details. Electrophysiological recordings allow much higher temporal resolution, and *in vivo* whole-cell patch clamp recordings in behaving flies is a well-established technique (Wilson et al. 2004; Maimon et al. 2010). DNs often have large cell bodies that are amenable to electrophysiology, and our collection of driver lines now enable many of them to be targeted effectively by expressing fluorescent markers. We also anticipate that our driver lines will be useful for voltage imaging as new improvements in genetically encoded voltage sensors and optical methods make this method more feasible.

For all experiments using these lines, researchers are strongly encouraged to examine expression patterns themselves to evaluate line cleanliness and consistency. Ideally, the expression pattern should be visualized using the actual effector gene(s) used in experiments, as the pattern may be effector-specific (Meissner et al. 2025). We have also occasionally observed changes in expression pattern after many generations of stock maintenance, perhaps due to contamination, epigenetic changes, or mutation. Furthermore, it is good practice to replicate results using two or more lines to reduce or eliminate the confounding effects of off-target expression. Our collection includes two or more high-quality ‘A’ lines for only 36 DN types (Fig. 1e,f), but some lines with lower specificity may still provide useful validation if lines do not contain overlapping off-target expression.

Rough estimates suggested that Janelia’s Gen1 MCFO database should enable the creation of cell-type-specific lines for 50–80% of cell types (Meissner et al. 2023). However, after screening this database and surveying the literature for additional DN split-Gal4 driver lines, we report lines for only 190 (40%) of all DN types. We estimate this corresponds to ∼600 (45%) of the roughly 1300 DNs counted from EM data. This lower-than-expected rate may partly reflect the database’s slightly lesser focus on the VNC compared to the central brain. It may also suggest that cell-type-specific lines labelling new DNs can still be produced using existing images and split hemidrivers. We expect this may be feasible for some of the 54 (roughly 11% of) DN types that we have identified from broadly expressing Gal4 lines but that still lack specific split-Gal4 drivers. However, many of these were identified from only a single image. Using the split-Gal4 technique to generate additional cell-type-specific lines would require LM identification in at least one other driver line with expression in the desired DN type.

Approximately 240 DN types, or 50% of the total, have only been identified from EM data. While an intense, targeted effort may still uncover instances of these neurons in the Gen1 MCFO database, we believe our search was reasonably thorough. Further efforts using the methods we employ here are likely to yield diminishing returns, barring large-scale generation of new enhancer lines or at least new MCFO images of existing lines.

With the advent of connectomic data that enable precise identification of neurons involved in a given circuit, it is becoming increasingly urgent to gain genetic access to specific neurons of interest. New methods may help increase the efficiency of successfully generating desired split-GAL4 lines. For example, single-cell transcriptomic data enable the selection of gene pairs that often uniquely specify a single cell type. Knocking the two split-Gal4 halves into these loci are likely to produce a cell-type-specific line—a method successfully demonstrated by Chen et al. (2023). Spatial transcriptomics (Janssens et al. 2024) may be an important tool in facilitating this approach, allowing researchers to discover genes expressed in cell types that are identified morphologically from connectome data.

In summary, we complete our survey of existing resources for producing cell-type-specific DN lines and report a total of 806 lines covering roughly 40% of DN types. We anticipate that these lines will be useful for understanding the neural control of behaviour in *Drosophila melanogaster*.

## Acknowledgements

During this effort, the FlyLight Project Team consisted of: Megan Atkins, Kari Close, Gina DePasquale, Zachary Dorman, Kaitlyn Forster, Theresa Gibney, Joanna Hausenfluck, Yisheng He, Kristin Henderson, Nirmala Iyer, Jennifer Jeter, Lauren Johnson, Rebecca Johnston, Rachel Lazarus, Kelley Lee, Hsing Hsi Li, Hua-Peng Liaw, Oz Malkesman, Geoffrey Meissner, Brian Melton, Scott Miller, Alexandra Novak, Omotara Ogundeyi, Alyson Petruncio, Jacquelyn Price, Sophia Protopapas, Susana Tae, Allison Vannan, Rebecca Vorimo, Brianna Yarbrough, and Kevin Xiankun Zeng, and the Steering Committee consisted of: Gerry Rubin, Barry Dickson, Reed George, Gwyneth Card, Wyatt Korff, Geoffrey Meissner. We also thank Gudrun Ihrke, Manager of Project Technical Resources, for helping to coordinate additional imaging and supervision of Claire Managan’s efforts in performing neuron segmentations from light-level data.

This work was part of the Descending Interneuron Project Team at Janelia Research Campus, part of the Howard Hughes Medical Institute. JLZ was partly supported by an Alan Kanzer Zuckerman Institute Postdoctoral Fellowship. HSJC was supported by a Howard Hughes Medical Institute Janelia Research Campus Visiting Scientist Program project awarded to Dr. Kevin C. Daly, Dr. Andrew M. Dacks, HSJC, and GMC. MC, KE, TS, and GSXEJ were supported by Wellcome Trust Collaborative Award 220343/Z/20/Z. GMC is an Investigator of the Howard Hughes Medical Institute.

## Methods

### Identification of DN types

We generally employed the same methods as in Namiki et al. (2018), except that we occasionally used DN annotations from newly available EM datasets (Cheong et al. 2024; Stuerner et al. 2024; Schlegel et al. 2024) to help clarify morphology and aid in segmentation.

In brief: We manually searched the Janelia Gen1 MCFO database (Meissner et al. 2023) to identify distinct descending neuron types. Some types were also identified from lines found in our literature search (see section “Literature search for previously published lines”). For newly identified types, we selected a representative image from either a Gen1 Gal4 line or a sparsely labelled split-Gal4 line and semi-manually produced the segmentations shown in Fig. 2–26. Image signal was thresholded to create a mask using the Amira ‘Interactive Thresholding Function’ or the Fiji ‘Threshold’ function, then manually corrected, often using other LM images and/or matching EM neurons as a reference. We then applied this mask to the original images to produce the final images shown in Fig. 2–26.

### Creation of split-Gal4 lines

We employed the same methods as in Namiki et al. (2018). We created split-Gal4 driver lines by crossing hemidriver lines (containing either the GAL4 activation domain (AD) or DNA binding domain (DBD)) from the collections described by Tirian and Dickson (2017) and Dionne et al. (2018). Candidate combinations of split-Gal4 halves were chosen based on images in the Janelia Gen1 MCFO database (Meissner et al. 2023). We searched this database semi-manually, using the Color Depth MIP Mask Search tool (Otsuna et al. 2018) on Janelia Workstation software (Rokicki et al. 2019) to identify candidate split halves that contained the target neuron.

Combinations producing a line that sparsely labelled the target cell type were made into stable stocks (i.e., homozygous or carrying balancer chromosomes). Final expression patterns were evaluated by crossing stable driver lines to a line containing 20XUAS-CsChrimson-mVenus in attP18. Select lines were also crossed to a reporter line containing 5XUAS-IVS-myr::smFLAG in VK00005, pJFRC51- 3XUAS-IVS-Syt::smHA in su(Hw)attP1 to obtain polarity information. For lines labelling multiple cell types, we often also used the MCFO technique (Nern et al. 2015) to discern the morphology and identity of individual neurons.

### Naming of DN types

DN types were given systematic names consisting of a prefix indicating the location of the soma in the central brain and a sequential number. Occasionally, a number was skipped when we originally assigned a name to a neuron, but it was later found to be a duplicate of another type. Synonyms are shown in Fig. 2–26 if we are aware of functional work using that synonym. A more comprehensive list of synonyms (including prior names assigned in purely anatomical studies) is shown in Supplementary Table 1.

### Line annotation

For the 500 lines newly released in this paper, we manually examined all available images of a given line (in colour-depth maximum-intensity-projection format) and compared them against our ground-truth light-microscopy (LM) images that are listed in Supplementary Table 2 and shown in Fig. 2–26. We also used NeuronBridge (Clements et al. 2024) to discover additional candidate matches in LM and EM datasets. In cases where our ground-truth LM images could be confidently matched to neurons in FAFB, FANC, and/or MANC EM datasets (Cheong et al. 2024; Stuerner et al. 2024; Schlegel et al. 2024), we used the clearer EM images to aid in identifying differences between similar-looking neurons. Scores of “Confident”, “Probable”, and “Candidate” respectively correspond to confidence levels of >95%, 70–95%, and 30–70% that the line labels the DN type(s) shown in Fig. 2–26 and in the example slides listed in Supplementary Table 2. However, we have not exhaustively compared all images with DN types known only from EM data. Therefore, it is possible that some lines in fact label a neuron outside our list of 244 defined types but were misannotated with a morphologically similar DN type. All images of our 500 lines have been deposited at https://splitgal4.janelia.org/ or https://flylight-raw.janelia.org/.

For previously published lines, we relied primarily on published annotations. For lines given “Confident”, “Probable”, or “Candidate” scores, we checked published annotations and updated them when necessary using the methods described in the previous paragraph and images available at https://splitgal4.janelia.org/ and https://flylight-raw.janelia.org/. Annotations for lines marked “From literature” in the “Annotation_confidence” column in Supplementary Table 5 rely solely on published annotations.

### Scoring of line specificity and consistency

Line specificity and consistency was evaluated using all available Janelia images that did not use stochastic effectors (i.e., all non-MCFO images). All images have been deposited at https://splitgal4.janelia.org/ or https://flylight-raw.janelia.org/.

We used the following criteria for line quality scores:

A - Target cell type is consistently expressed bilaterally. Background expression is zero or in very few cell types. Any background is extremely faint and typically also stochastic relative to expression of the target cell type.

B - Fails one of the three criteria for A. E.g., may have low/no background, but stochastic expression; sparse but bright background expression; or faint, diffuse background expression.

C - Target cell type is very inconsistently expressed and/or there is high background expression, especially background that overlaps spatially with the target cell type.

### Literature search for previously published lines

We searched the following query using Google Scholar: *drosophila “split-gal4” “descending neuron”*. This search returned 165 results. We skimmed all primary research articles for indications that the authors had produced new split-Gal4 lines sparsely labelling DNs in *Drosophila melanogaster* adults. Any such lines were added to our list if we could confidently match them to any of our described cell types.

We also manually checked expression patterns for all lines in the Janelia omnibus database (Meissner et al. 2025) that met both of the following two criteria: (1) Annotations were missing or ambiguous or included the string “DN”. (2) Expression pattern was annotated as not being restricted to only the brain or only the VNC. All lines that labelled identifiable DNs were added to our list.

---

**Supplementary Figure 1.**
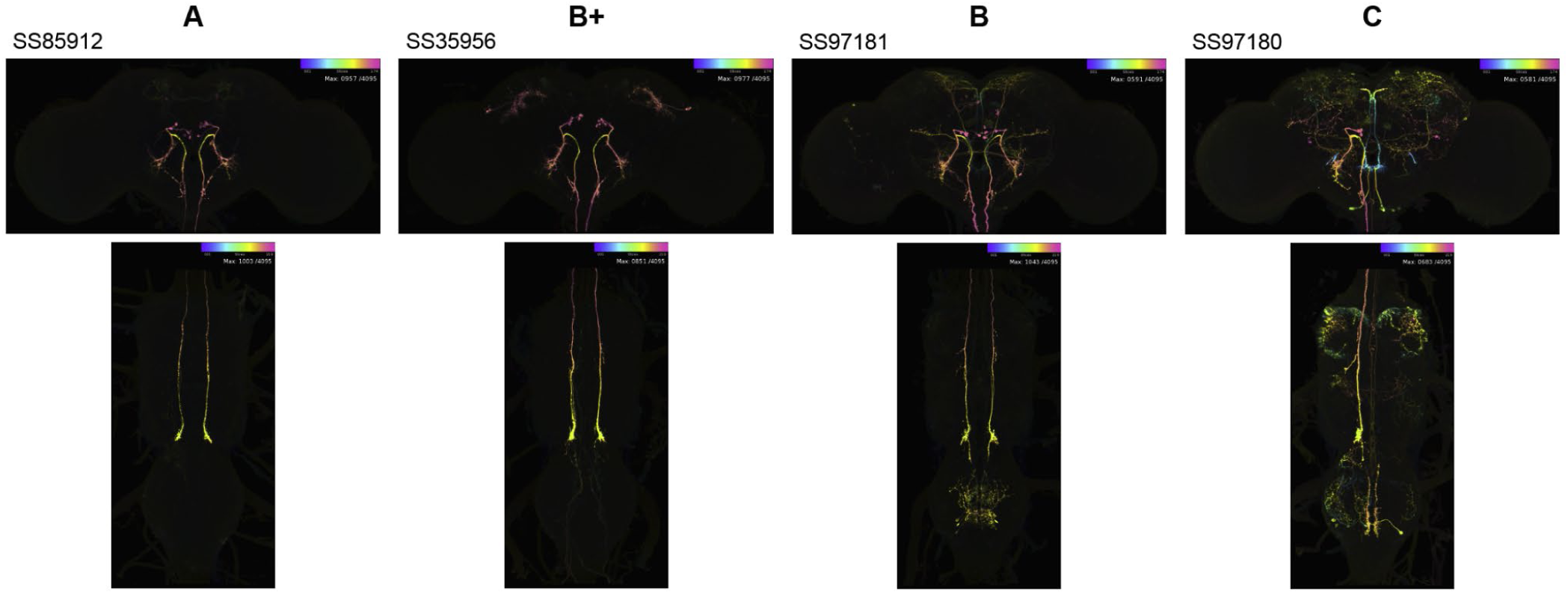
Maximum-intensity projections showing DNp17 lines representative of ‘A’, ‘B+’, ‘B’, and ‘C’ scores for cleanliness/consistency. Colour indicates depth in the Z axis (violet→pink = anterior→posterior in the brain and ventral→dorsal for the VNC).

**Supplementary Table 1.** List of all DN types identified from light-level microscopy and split-Gal4 lines that label them (from this study or any previously published source we could find). Known synonyms and their sources are also listed. Lines were given quality scores from ‘A’ (best) to ‘C’ (worst) for their specificity and consistency (see Methods for additional details). If any type had fewer than two ‘A’- quality lines, we bumped ‘B’ lines to a ‘B+’ category if they met a certain minimum standard. Asterisks mark lines with tentative identifications for a given DN type (see Supplementary Table 5).

**Supplementary Table 2.** Example slide codes for 3D stacks that show the morphology of each DN type. Images may be found at https://splitgal4.janelia.org/ and https://flylight-raw.janelia.org/.

**Supplementary Table 3.** List of updates to the morphology of DNs described in Namiki et al. (2018).

**Supplementary Table 4.** List of annotation updates for split-Gal4 lines described in Namiki et al. (2018). Cell-type annotations were removed if images available online (at https://splitgal4.janelia.org/ and https://flylight-raw.janelia.org/) did not show the cell type listed or showed it was extremely faintly expressed.

**Supplementary Table 5.** List of all split-Gal4 lines labelling descending neurons that were produced in this study, produced in Namiki et al. (2018), or found in other previously published work. “Bloomington_ID” lists stock IDs for lines maintained at the Bloomington *Drosophila* Stock Center at the time of publication. The “Previous_annotation” column lists published annotations for previously published lines. “Cell_type” lists clean annotations using systematic names, updated from published annotations where appropriate. An ‘x’ in the “Unannotated_DN” column denotes lines where there is clear evidence of one or more unannotated but consistently expressed DN types, but available images do not show the morphology clearly enough for identification. See Methods for further details on “Annotation_confidence” and “Quality_score”.

**Supplementary File 1.** One representative maximum-intensity projection showing the expression pattern in the brain and VNC for each new split-Gal4 line produced in this study. Colour indicates depth in the Z axis (violet→pink = anterior→posterior in the brain and ventral→dorsal for the VNC). Images are aligned to a unisex template. The complete set of images for all lines is available online at https://splitgal4.janelia.org/ and https://flylight-raw.janelia.org/.

