## Supplementary File 1 for "An updated catalogue of split-GAL4 driver lines for descending neurons in *Drosophila melanogaster*"

SS52394 - DNa01, DNa13

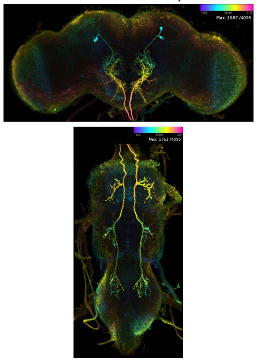

SS01591 - DNa02

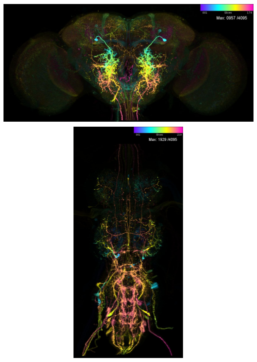

SS66179 - DNa02

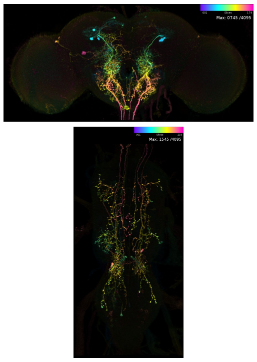

SS89717 - DNa02

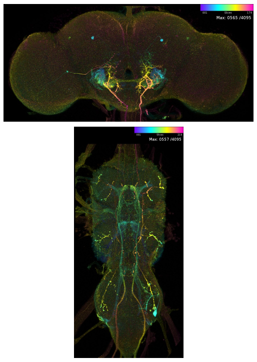

SS65478 - DNa02, DNa04, DNa09, DNa10

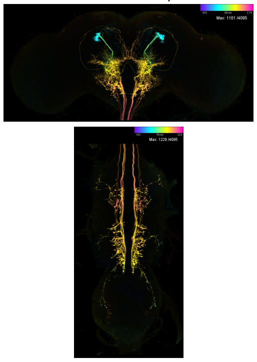

SS65543 - DNa02, DNa16, DNp15

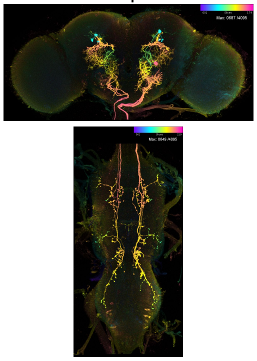

SS13529 - DNa04

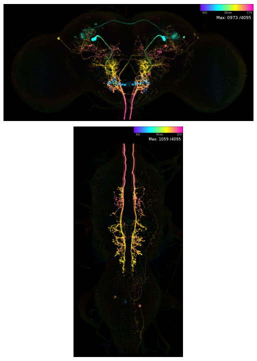

SS53394 - DNa04

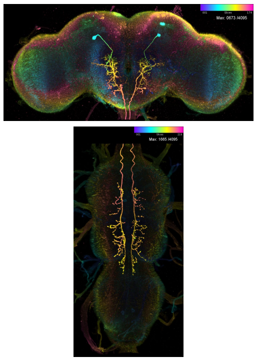

SS55587 - DNa04

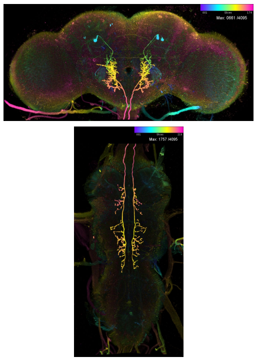

SS68367 - DNa04

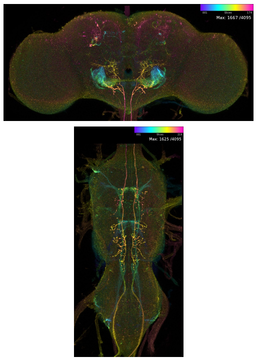

SS94762 - DNa04

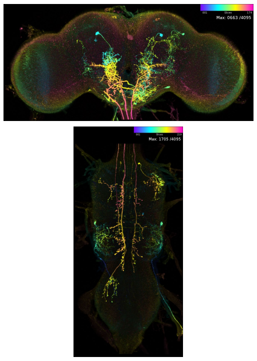

SS55594 - DNa04, DNa05

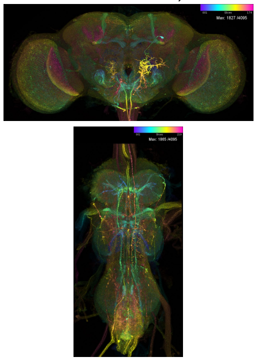

SS69285 - DNa04, DNa05

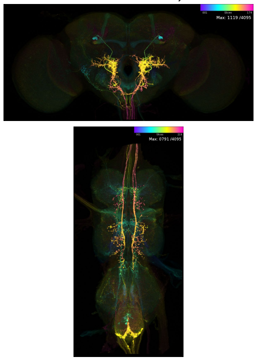

SS64864 - DNa04, DNa05, DNa06, DNa09

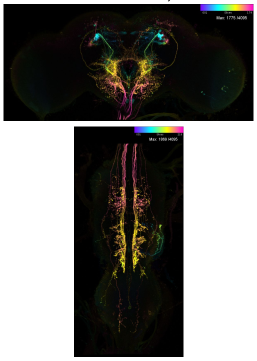

SS55580 - DNa04, DNa05, DNa07

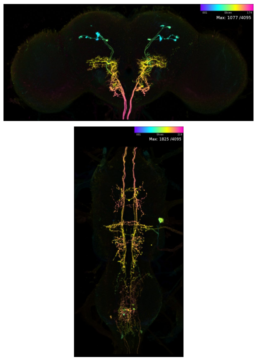

SS65484 - DNa04, DNa05, DNa09, DNa16

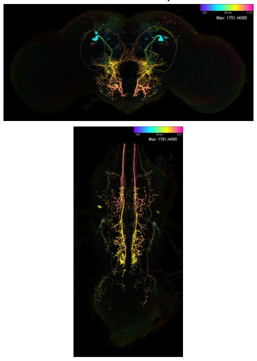

SS52376 - DNa04, DNa05, DNb08

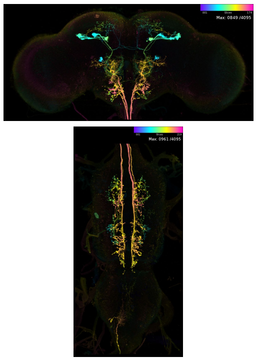

SS55578 - DNa04, DNa11

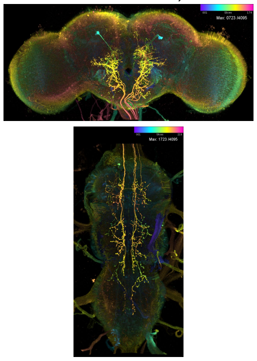

SS27191 - DNa04, DNa15

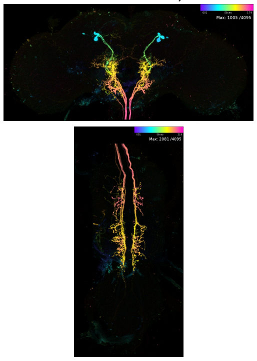

SS55584 - DNa04, DNb08

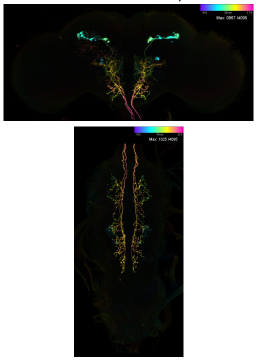

SS68369 - DNa04, DNb08

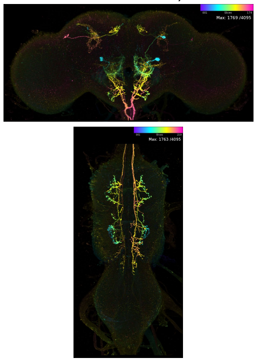

SS53405 - DNa05

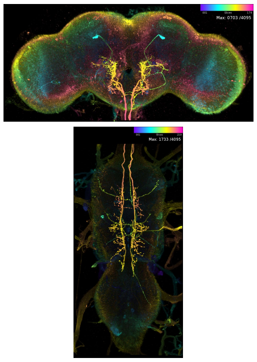

SS95335 - DNa05

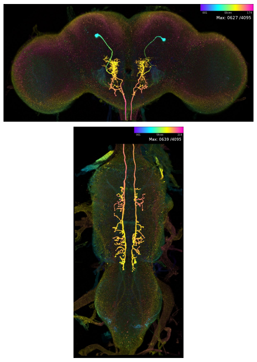

SS95343 - DNa05

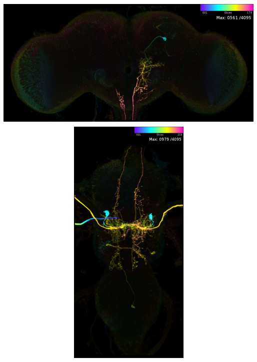

SS52447 - DNa05, DNp20

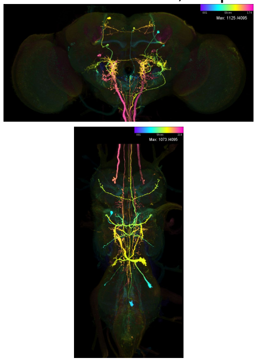

SS66156 - DNa06

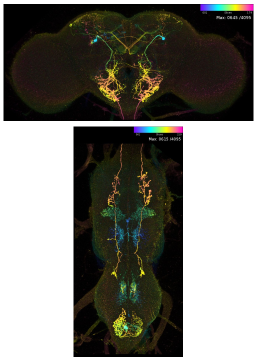

SS65482 - DNa06, DNa09

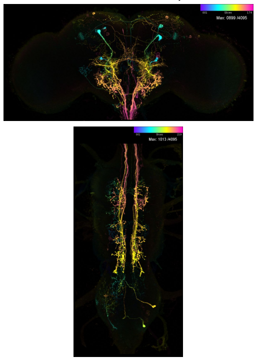

SS52370 - DNa07

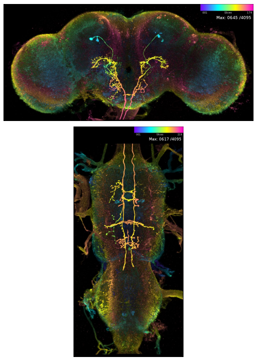

SS55576 - DNa07

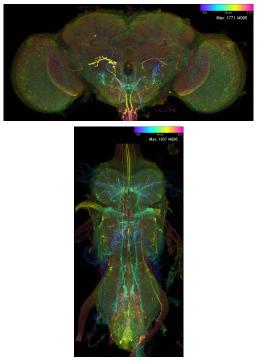

SS55579 - DNa07

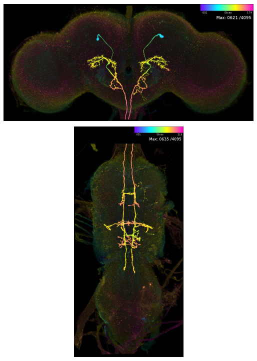

SS88193 - DNa07

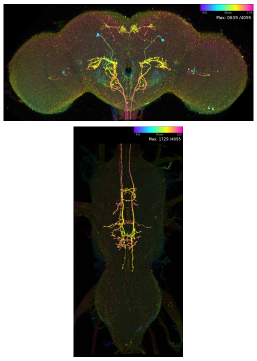

SS89587 - DNa07

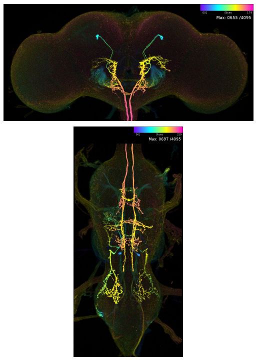

SS94749 - DNa07

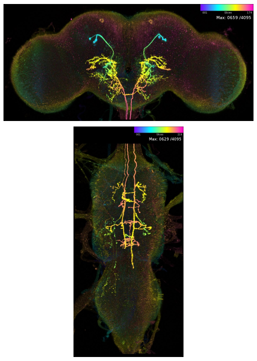

SS64887 - DNa09

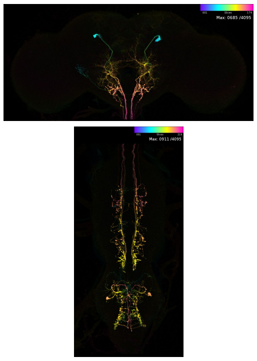

SS65480 - DNa09

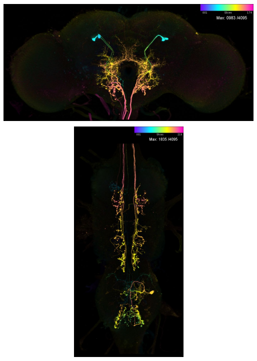

SS65486 - DNa09

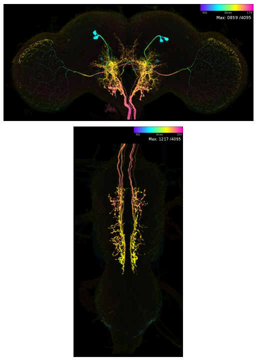

SS65487 - DNa09, DNa16

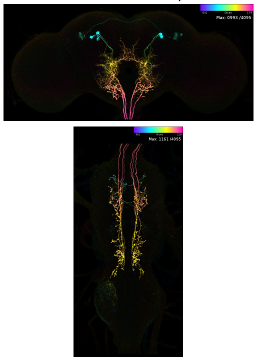

SS65540 - DNa10

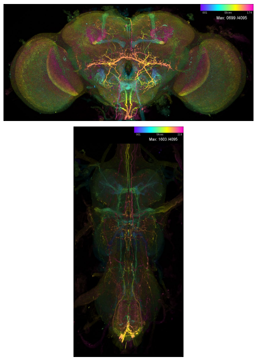

SS66183 - DNa10

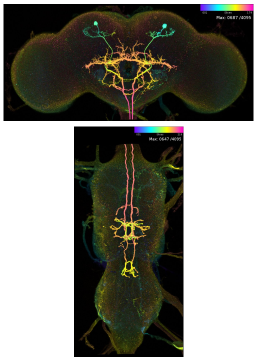

SS70534 - DNa10

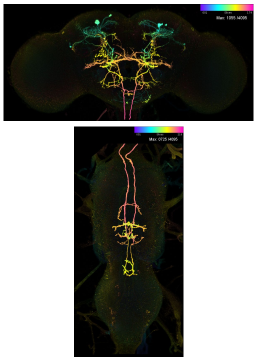

SS70607 - DNa10

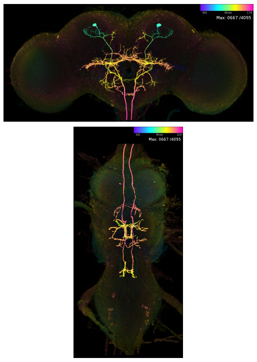

SS95287 - DNa11

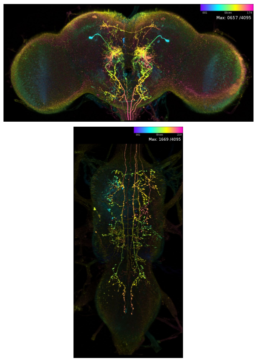

SS41452 - DNa13

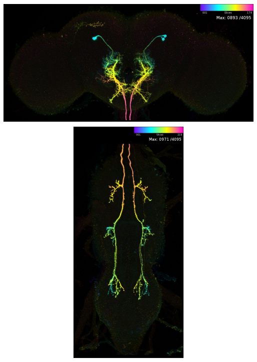

SS58308 - DNa13

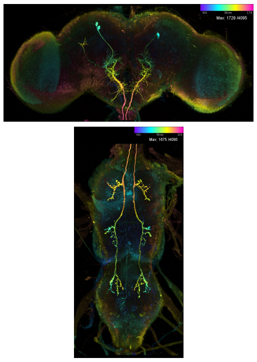

SS95327 - DNa13

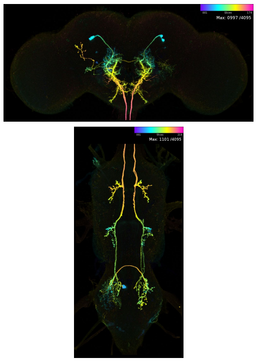

SS63697 - DNa16

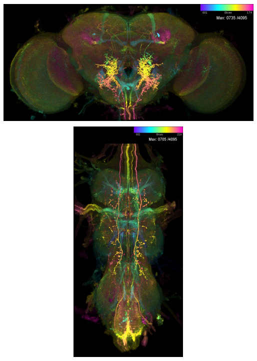

SS65483 - DNa16

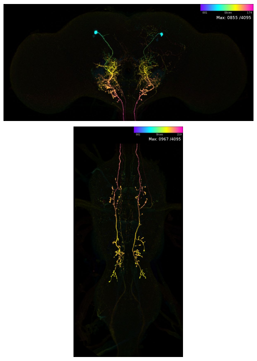

SS65544 - DNa16

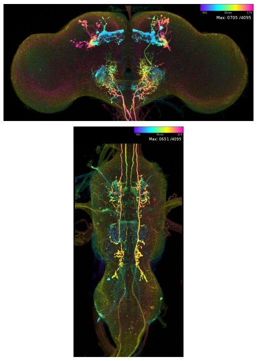

SS72170 - DNb01

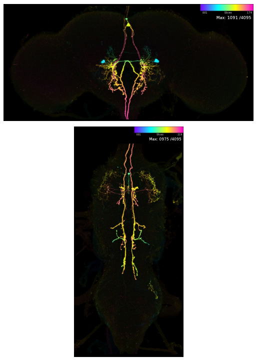

SS72179 - DNb01

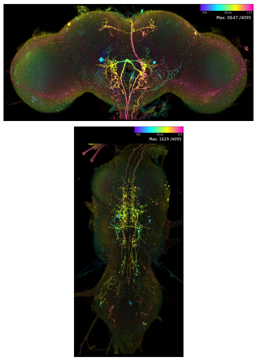

SS73097 - DNb01

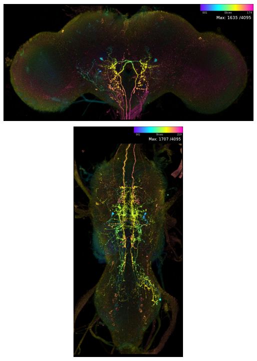

SS86130 - DNb03

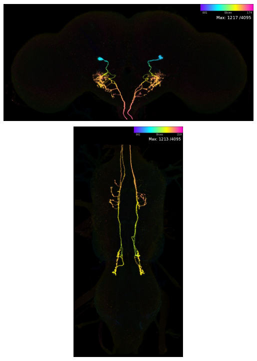

SS86231 - DNb03

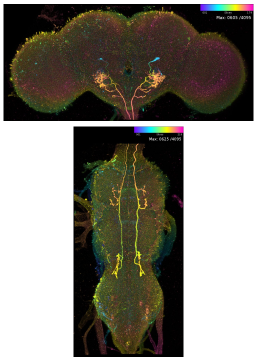

SS88143 - DNb03

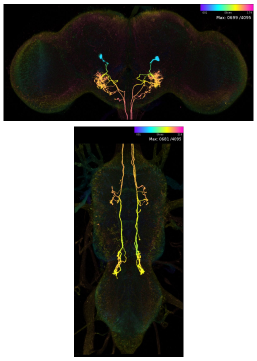

SS92786 - DNb03

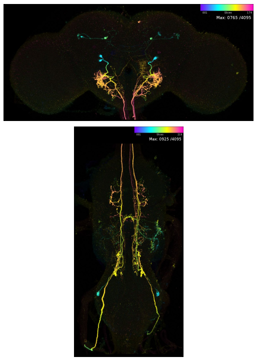

SS25635 - DNb04

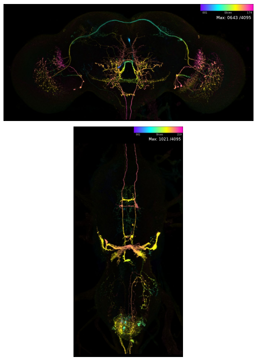

SS57817 - DNb04

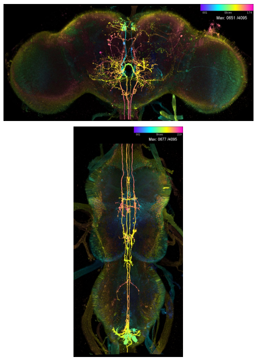

SS72972 - DNb04

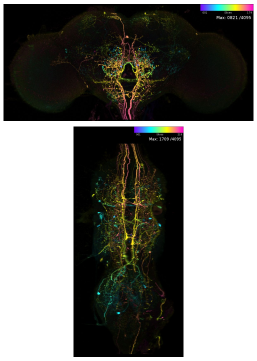

SS73069 - DNb04

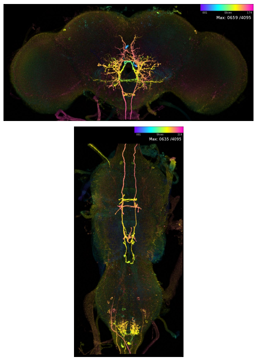

SS74475 - DNb04

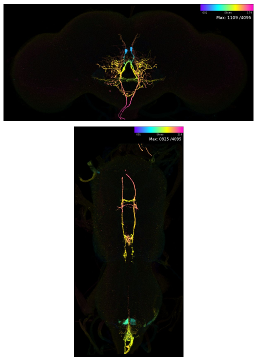

SS53402 - DNb05

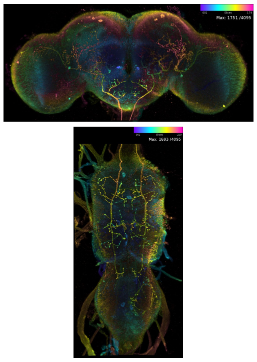

SS53436 - DNb05

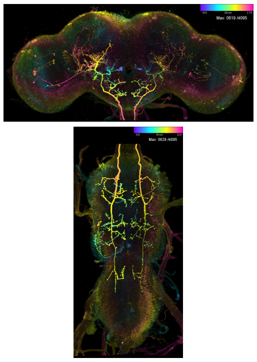

SS94697 - DNb05

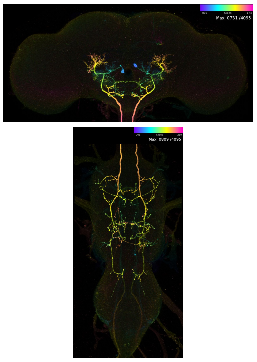

SS94698 - DNb05

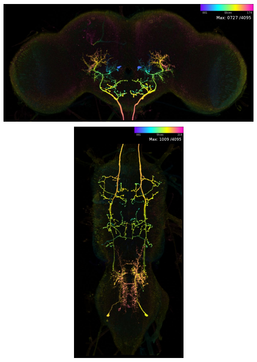

SS94700 - DNb05

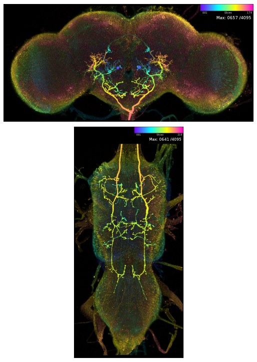

SS64589 - DNb08

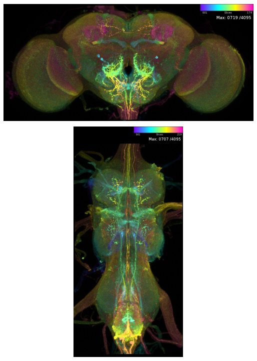

SS64604 - DNb08

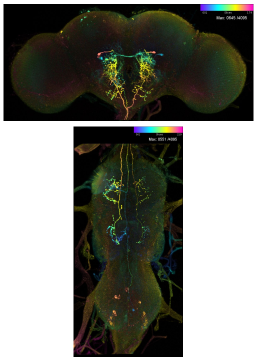

SS64607 - DNb08

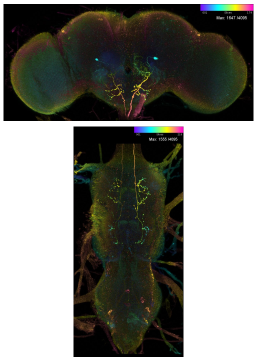

SS70590 - DNb08

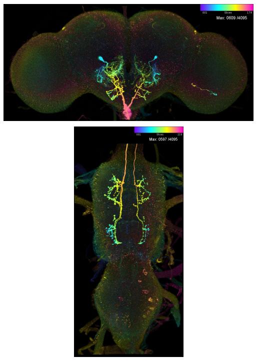

SS70620 - DNb08

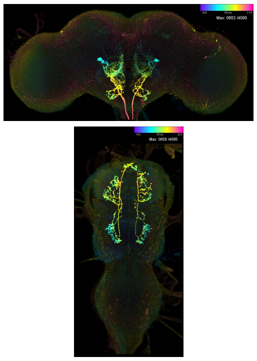

SS86129 - DNb08

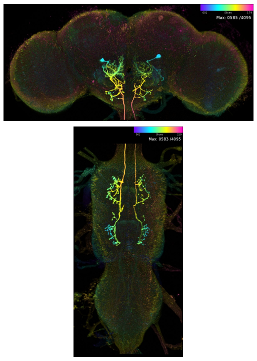

SS86213 - DNb08

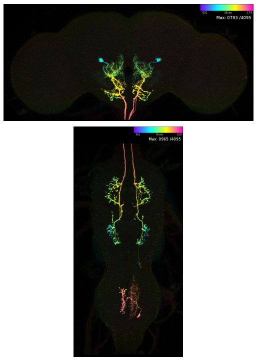

SS86277 - DNb08

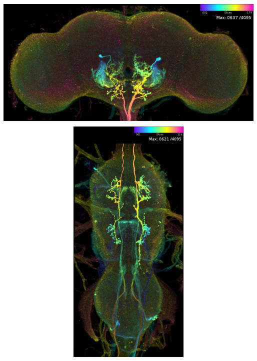

SS95330 - DNb09

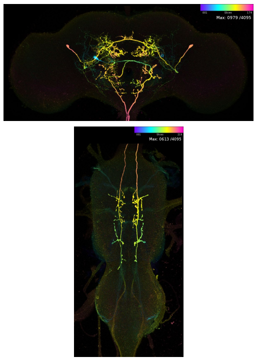

SS95408 - DNb09

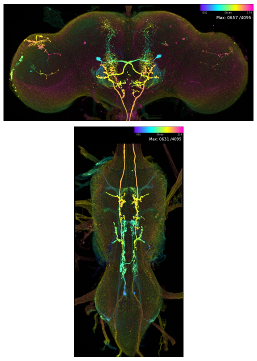

SS56194 - DNc01, DNc02

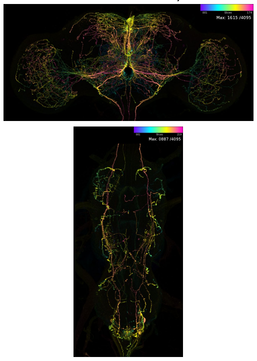

SS56196 - DNc01, DNc02

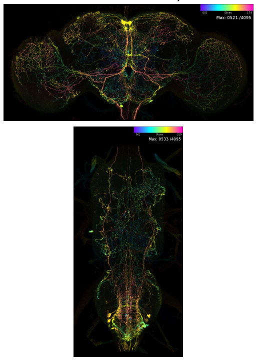

SS95441 - DNc01, DNc02

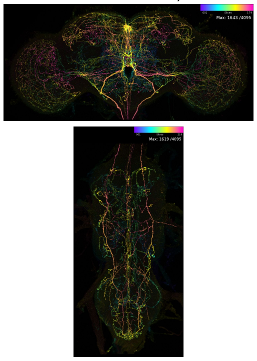

SS95458 - DNc01, DNc02

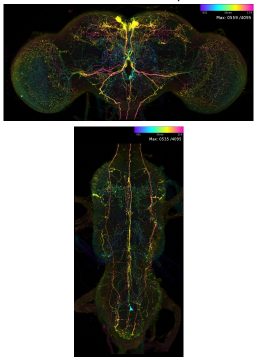

SS73077 - DNd01

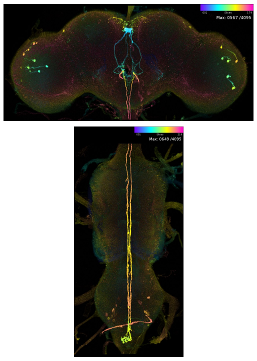

SS73441 - DNd01

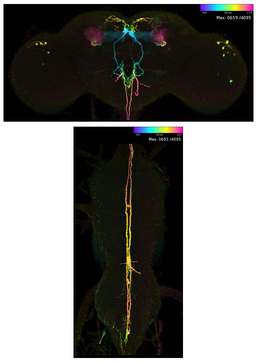

SS73444 - DNd01

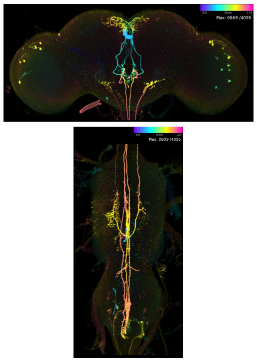

SS86097 - DNd01

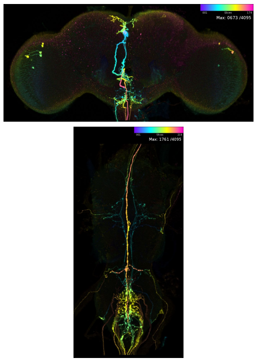

SS86098 - DNd01

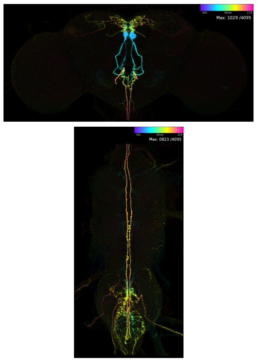

SS86283 - DNd01

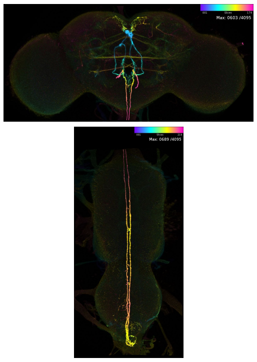

SS86341 - DNd01

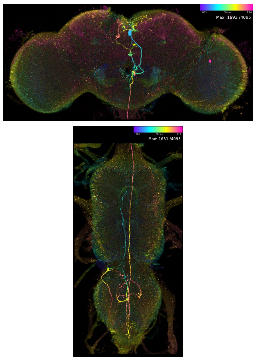

SS57753 - DNg01

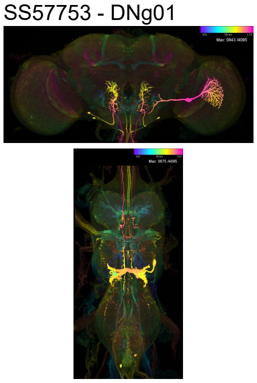

SS57798 - DNg01

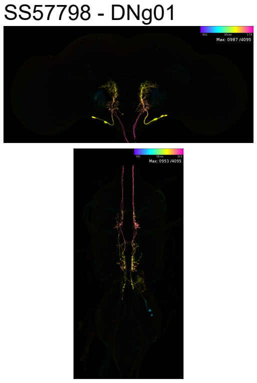

SS90870 - DNg01

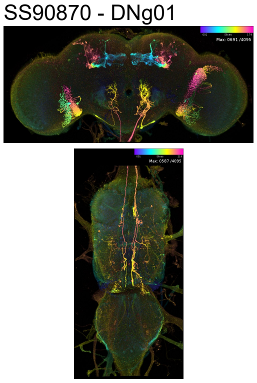

SS90918 - DNg01

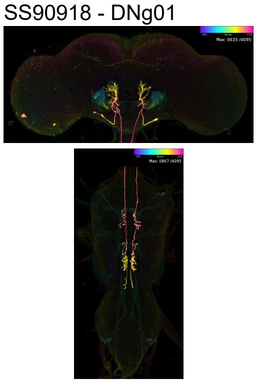

SS05120 - DNg01, DNg02

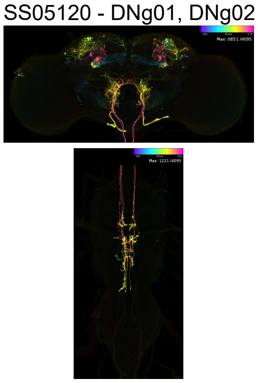

SS90856 - DNg01, DNg03

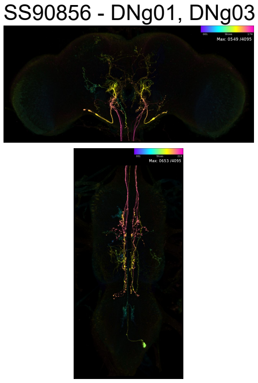

SS77445 - DNg01, DNg04, DNg82

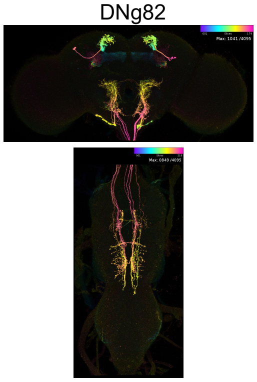

SS65577 - DNg01, DNg82

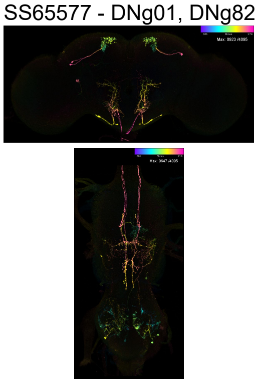

SS67125 - DNg01, DNg82

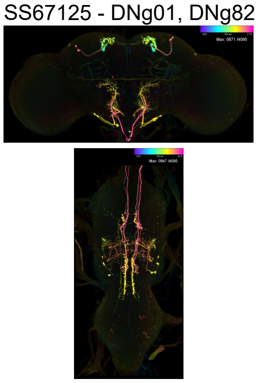

SS90868 - DNg01, DNg82

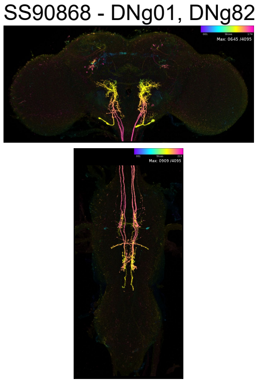

SS95356 - DNg01, DNg82

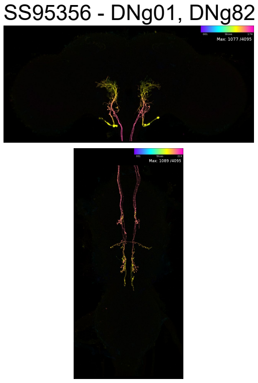

SS57754 - DNg01, DNg82, DNg91

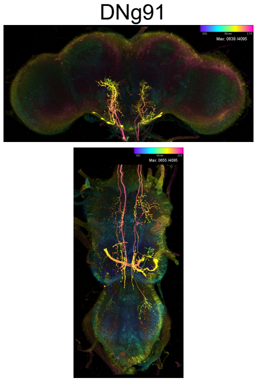

SS58304 - DNg01, DNg91

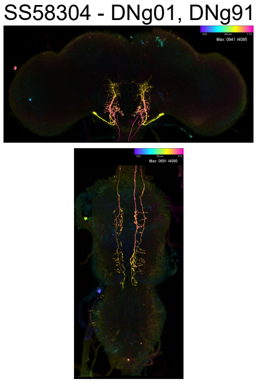

SS02380 - DNg02

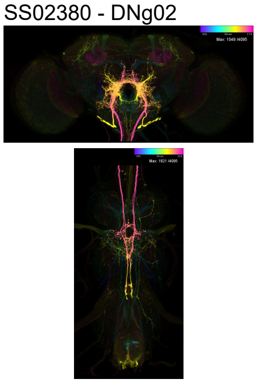

SS02543 - DNg02

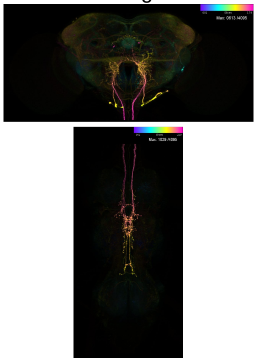

SS02549 - DNg02

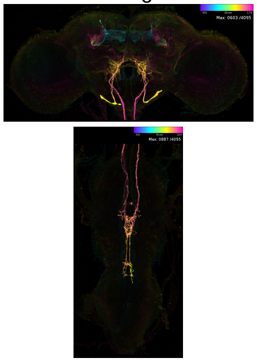

SS05121 - DNg02

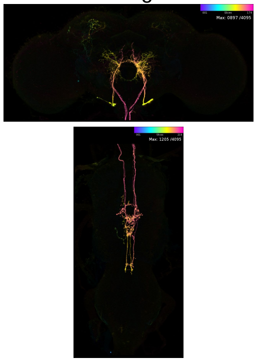

SS05135 - DNg02

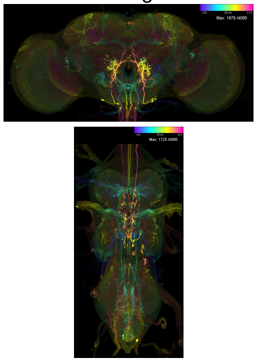

SS05136 - DNg02

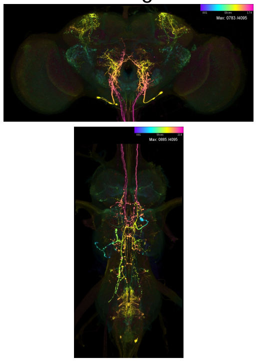

SS55460 - DNg02

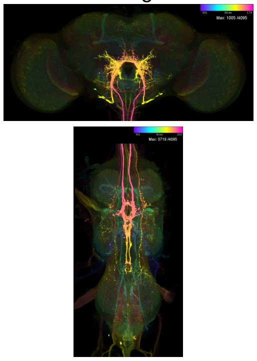

SS55954 - DNg02

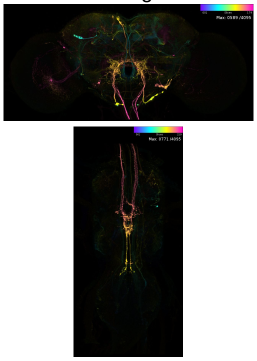

SS55956 - DNg02

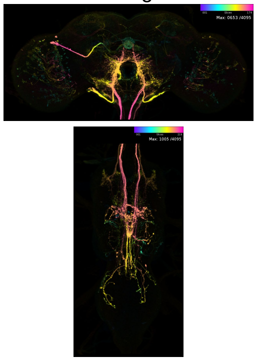

SS56995 - DNg02

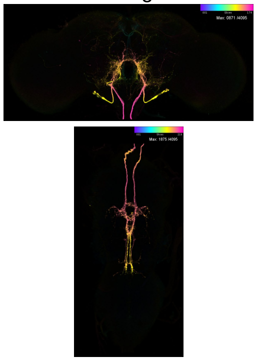

SS64637 - DNg02

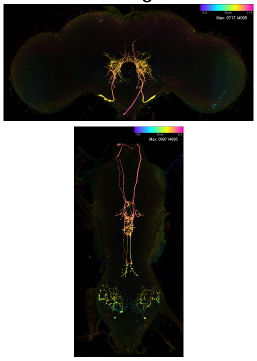

SS64638 - DNg02

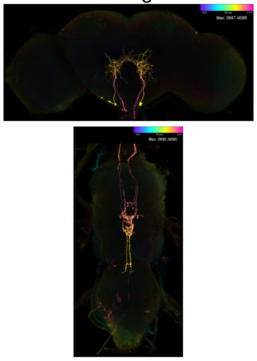

SS64680 - DNg02

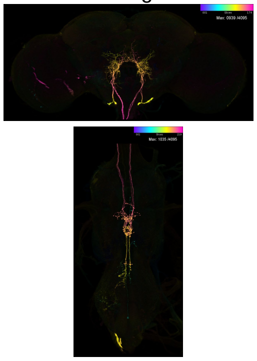

SS64721 - DNg02

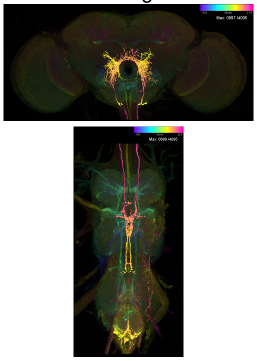

SS68365 - DNg02

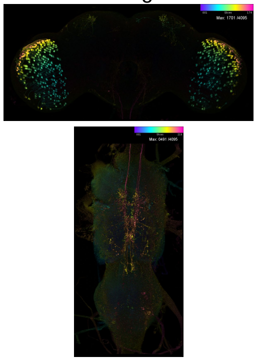

SS69379 - DNg02

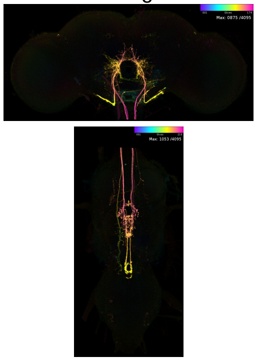

SS89574 - DNg02

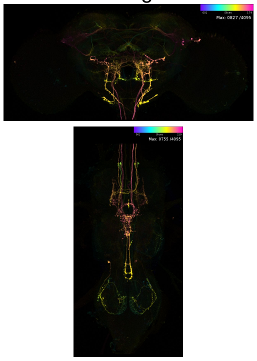

SS89626 - DNg02

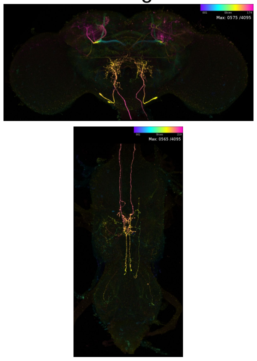

SS89637 - DNg02

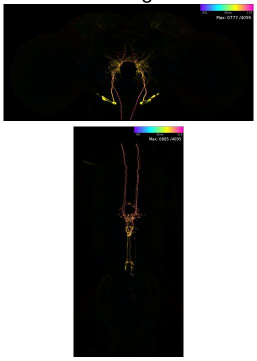

SS89648 - DNg02

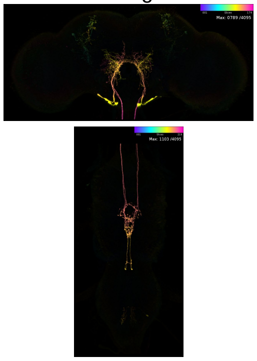

SS89691 - DNg02

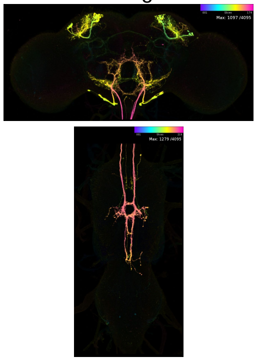

SS89712 - DNg02

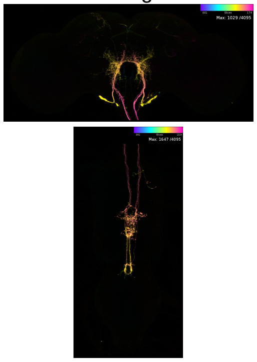

SS95557 - DNg02

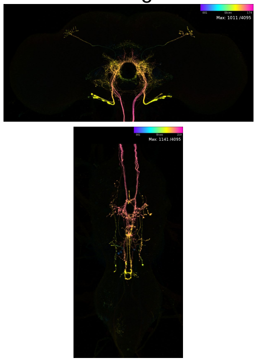

SS55459 - DNg02, DNg03

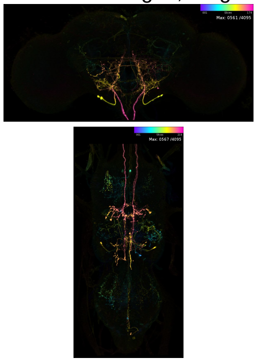

SS89710 - DNg02, DNg03

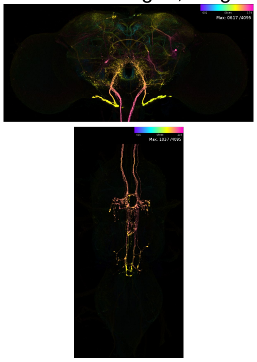

SS05098 - DNg03

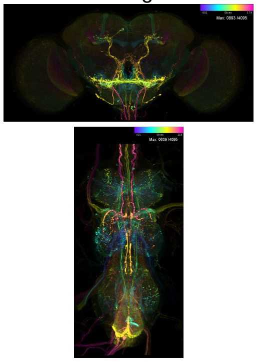

SS05108 - DNg03

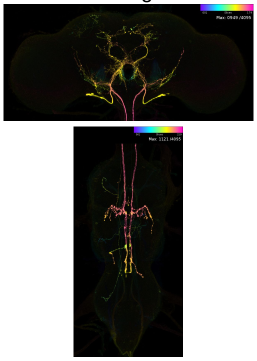

SS55951 - DNg03

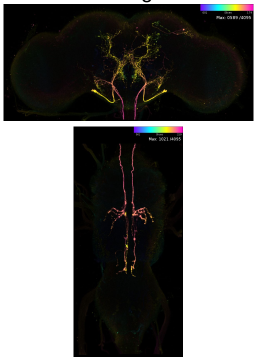

SS89559 - DNg03

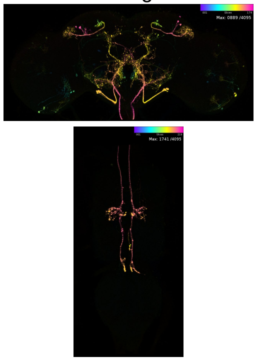

SS89561 - DNg03

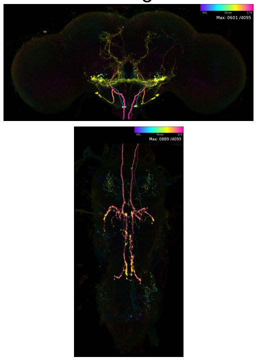

SS89562 - DNg03

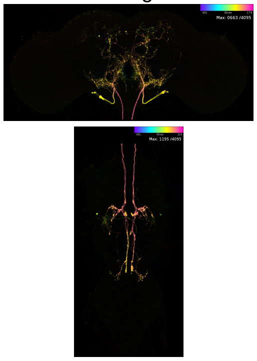

SS89564 - DNg03

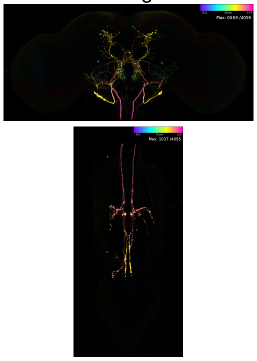

SS89565 - DNg03

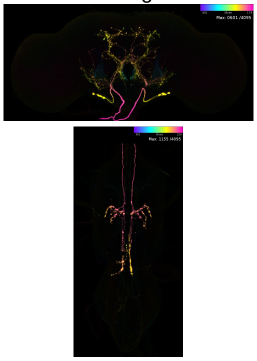

SS89566 - DNg03

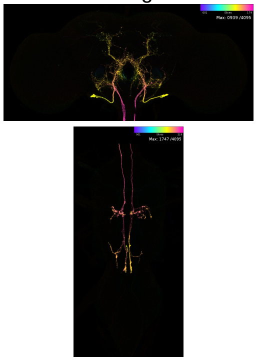

SS89568 - DNg03

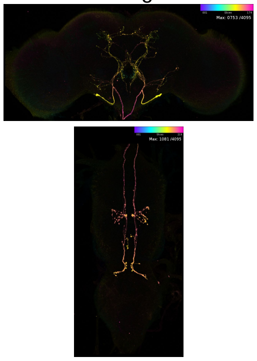

SS89572 - DNg03

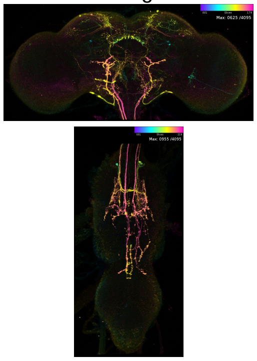

SS89598 - DNg03

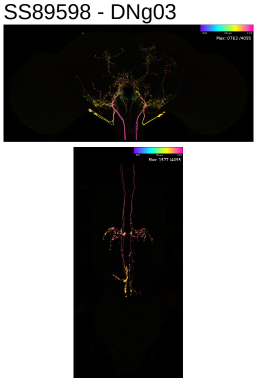

SS89602 - DNg03

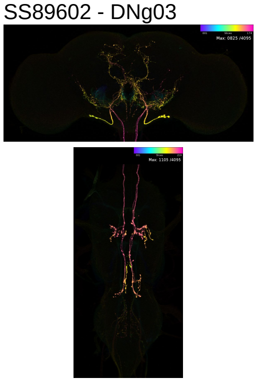

SS89603 - DNg03

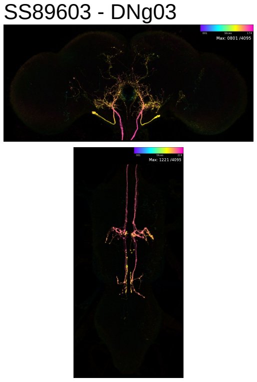

SS89611 - DNg03

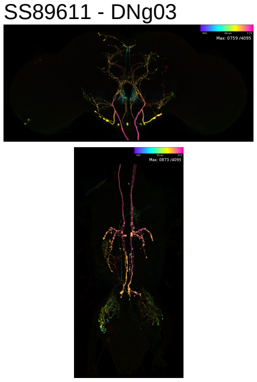

SS89640 - DNg03

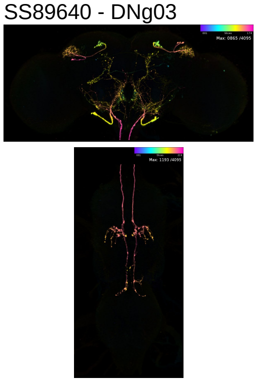

SS89644 - DNg03

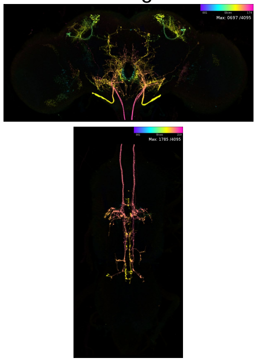

SS89645 - DNg03

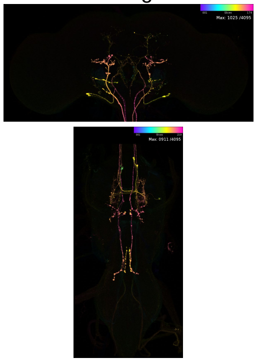

SS89650 - DNg03

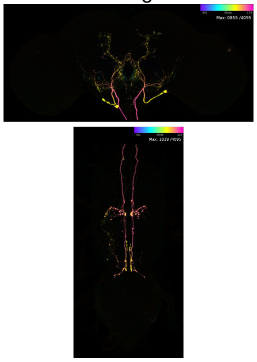

SS89651 - DNg03

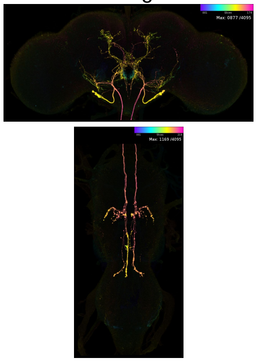

SS89654 - DNg03

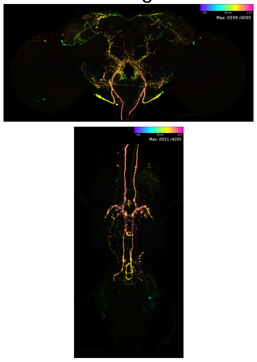

SS89656 - DNg03

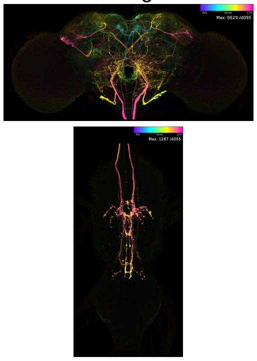

SS89657 - DNg03

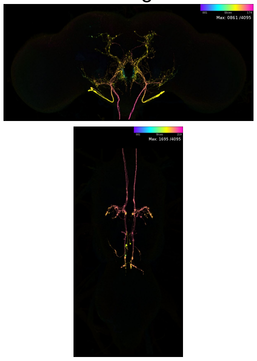

SS89661 - DNg03

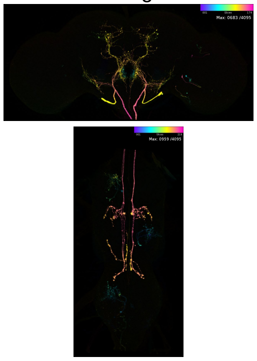

SS89669 - DNg03

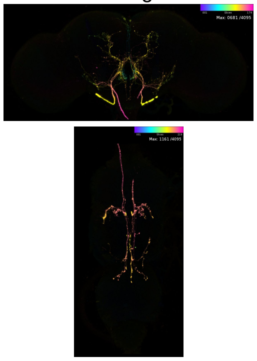

SS89674 - DNg03

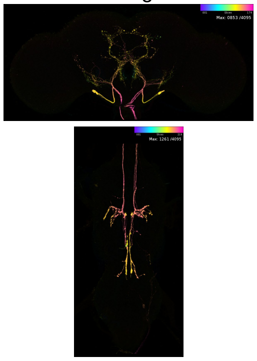

SS89676 - DNg03

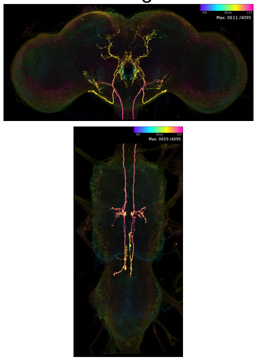

SS89684 - DNg03

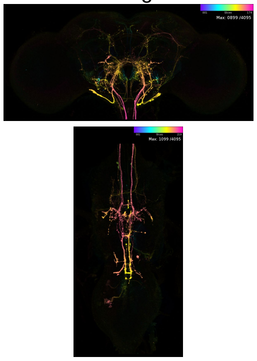

SS89711 - DNg03

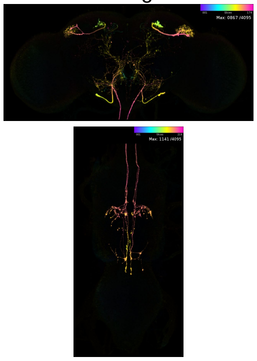

SS90854 - DNg03

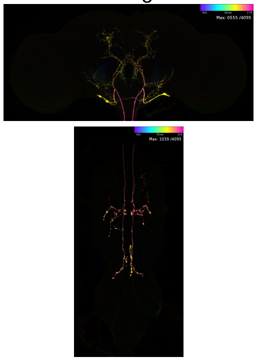

SS90917 - DNg03

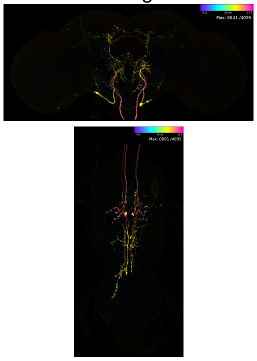

SS90919 - DNg03

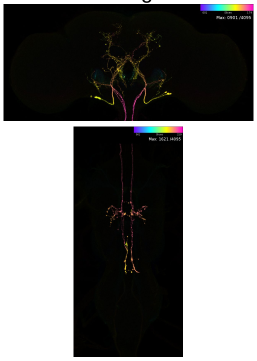

SS90922 - DNg03

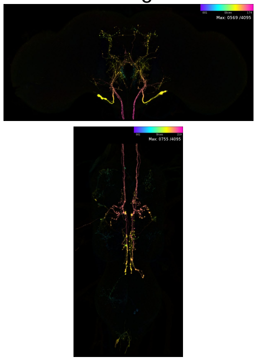

SS64679 - DNg03, DNg05

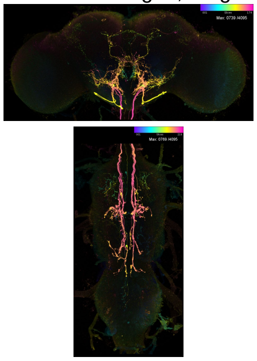

SS74390 - DNg04

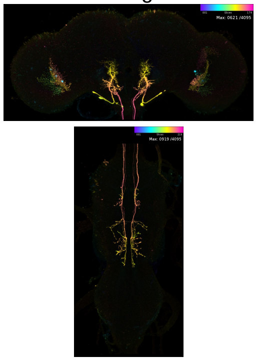

SS74434 - DNg04

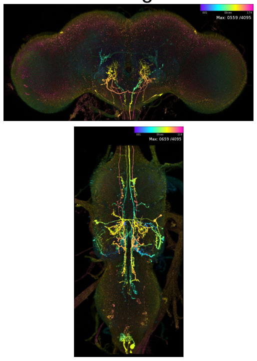

SS48205 - DNg05

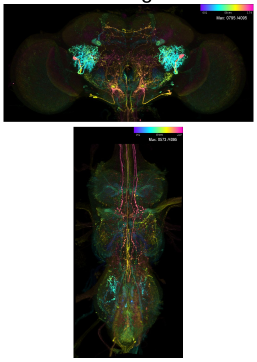

SS64574 - DNg05

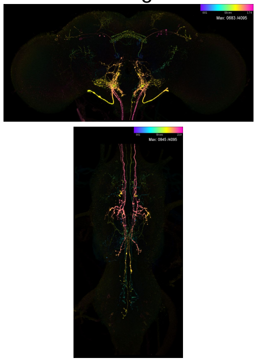

SS73455 - DNg05

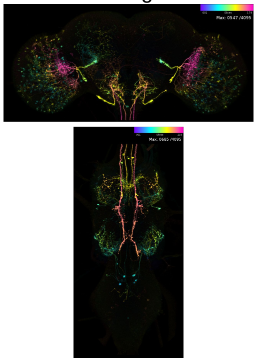

SS95560 - DNg06

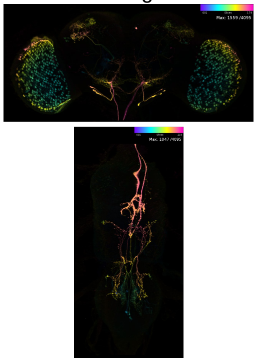

SS01586 - DNg07

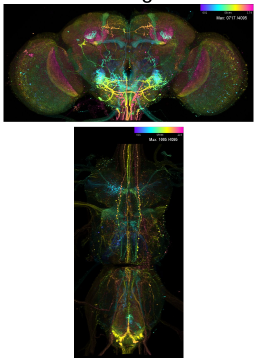

SS85845 - DNg07

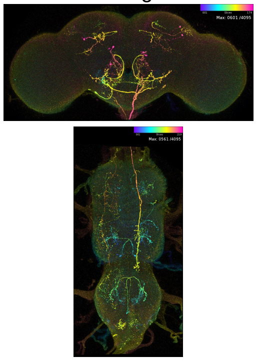

SS85901 - DNg07

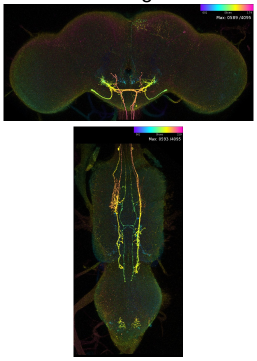

SS95796 - DNg07

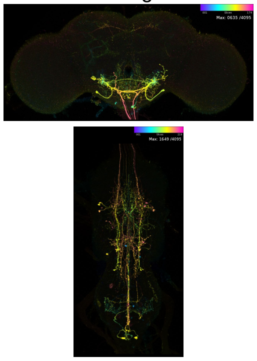

SS92805 - DNg08

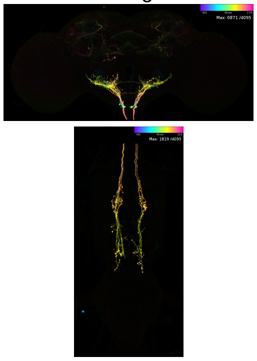

SS92759 - DNg08, DNg122

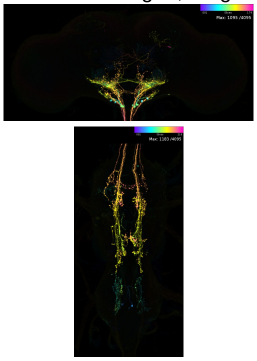

SS68388 - DNg09

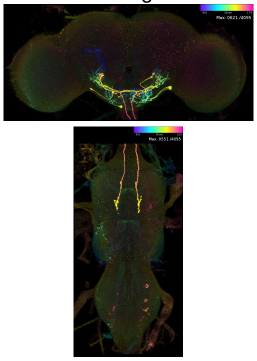

SS96100 - DNg09

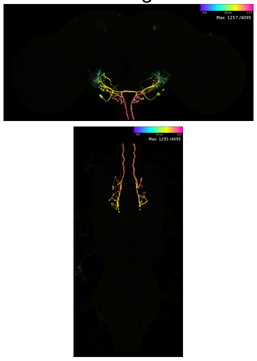

SS04518 - DNg10

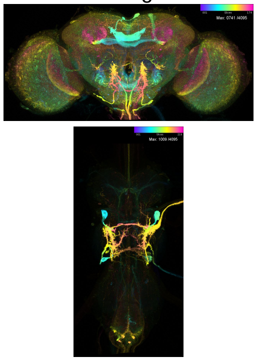

SS02389 - DNg10, DNg18

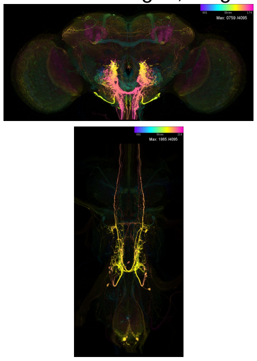

SS02390 - DNg10, DNg18

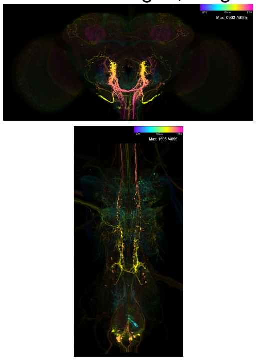

SS92722 - DNg10, DNp32

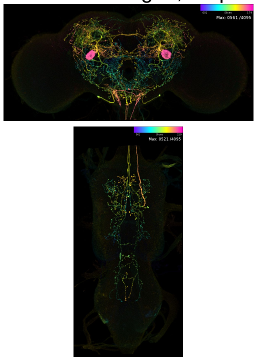

SS44071 - DNg11

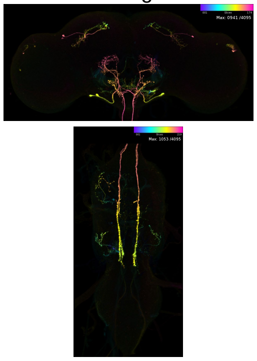

SS92297 - DNg12

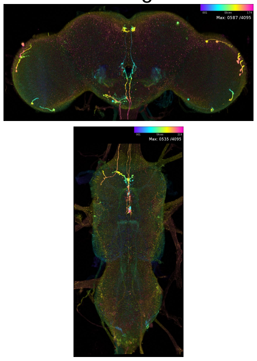

SS82228 - DNg13, DNp26

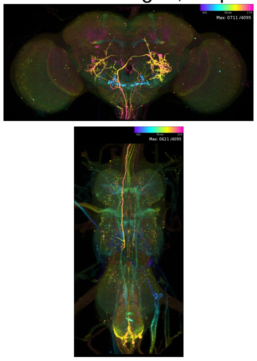

SS01055 - DNg14, DNg22

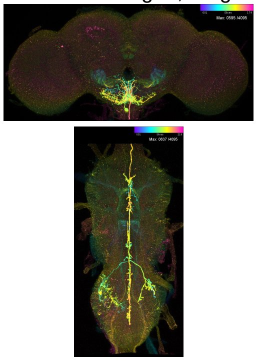

SS88264 - DNg14, DNg78

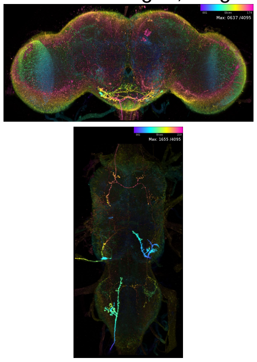

SS53432 - DNg16

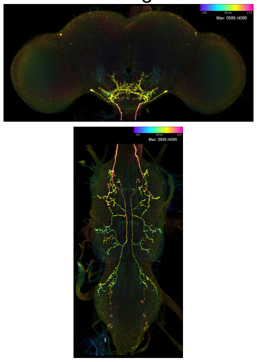

SS86205 - DNg16

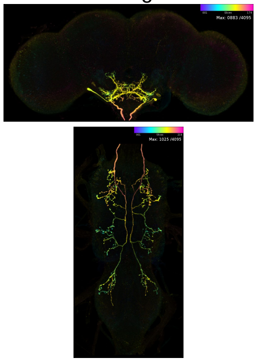

SS92616 - DNg16

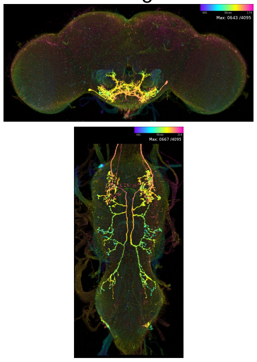

SS93749 - DNg16

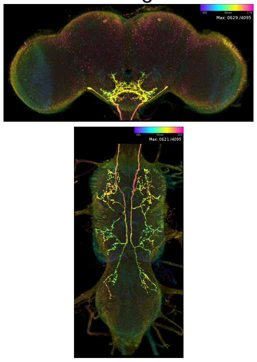

SS93750 - DNg16

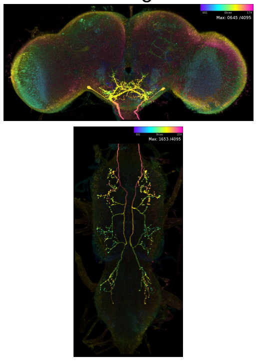

SS93767 - DNg16

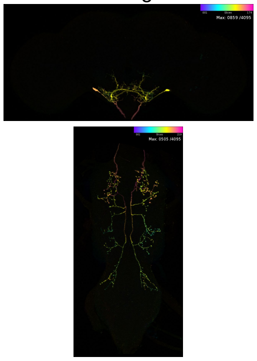

SS93771 - DNg16

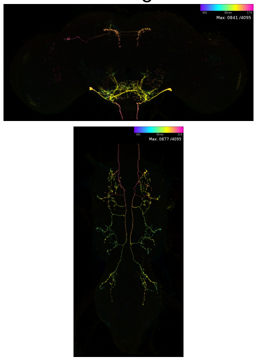

SS67063 - DNg16, DNg78

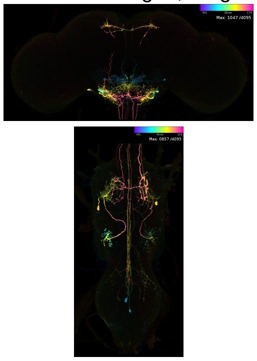

SS73433 - DNg16, DNg78

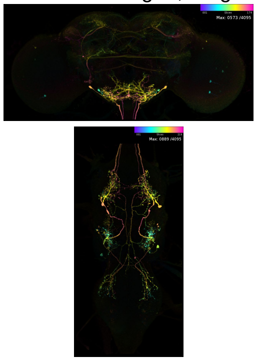

SS63693 - DNg18

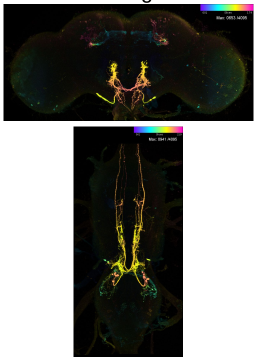

SS67107 - DNg18

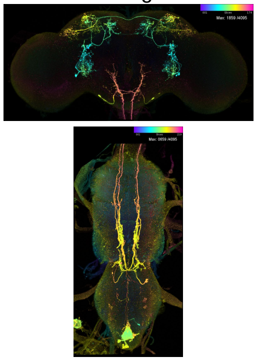

SS67131 - DNg18

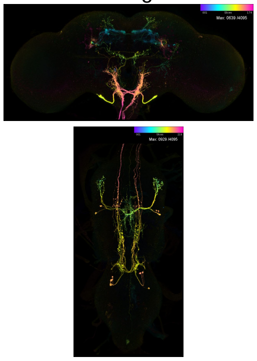

SS67174 - DNg18

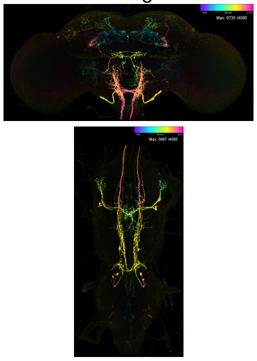

SS72104 - DNg18

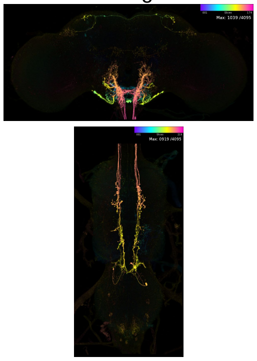

SS73446 - DNg18

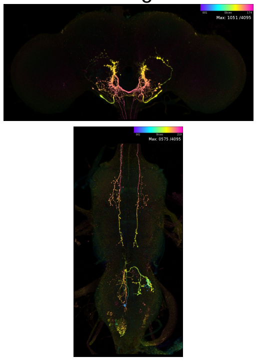

SS73447 - DNg18

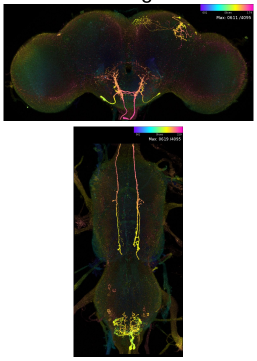

SS65581 - DNg20

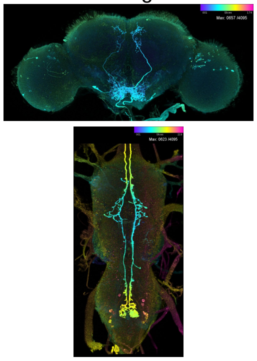

SS66270 - DNg20

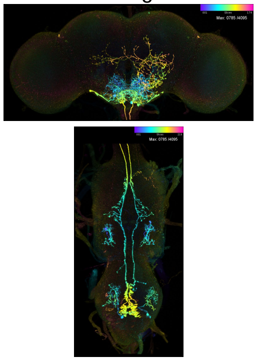

SS73102 - DNg20

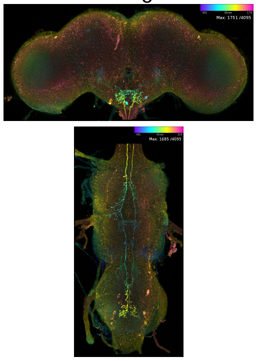

SS63678 - DNg21

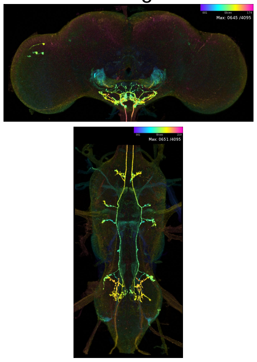

SS63720 - DNg21

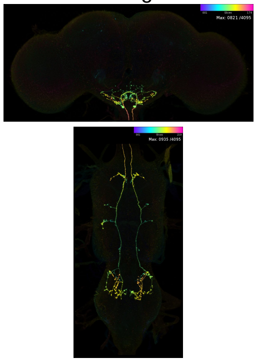

SS66220 - DNg21

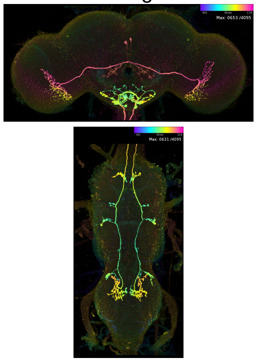

SS66264 - DNg21

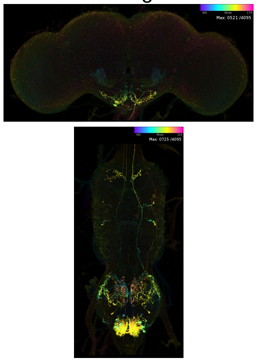

SS66271 - DNg21, DNg62

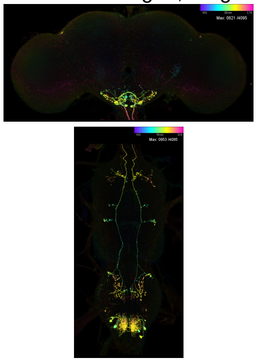

SS81919 - DNg21, DNg62

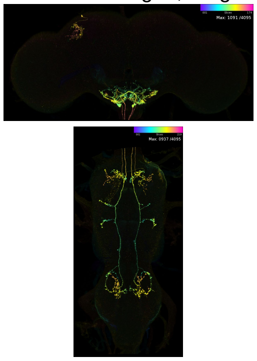

SS64673 - DNg22

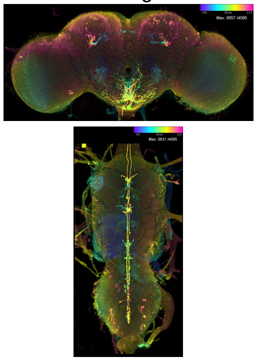

SS64702 - DNg22

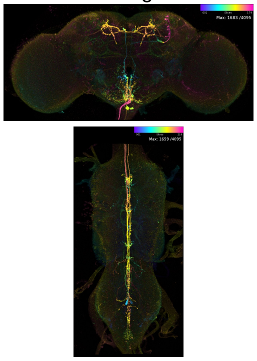

SS74393 - DNg22

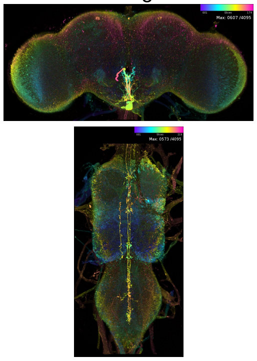

SS74442 - DNg22

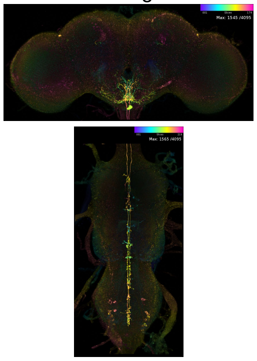

SS81923 - DNg22

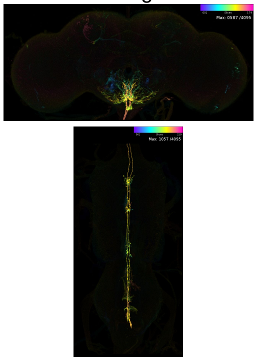

SS93166 - DNg22

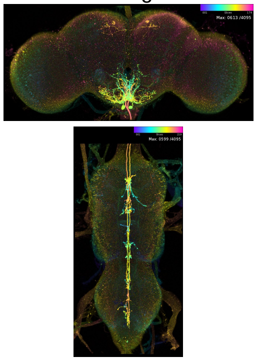

SS95368 - DNg22

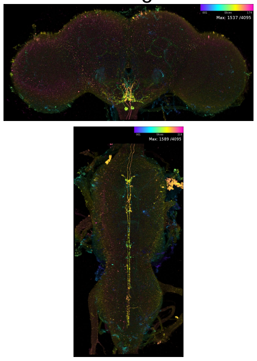

SS99387 - DNg24, DNp25

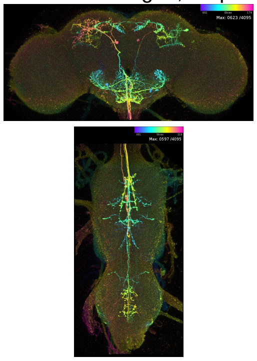

SS96111 - DNg26

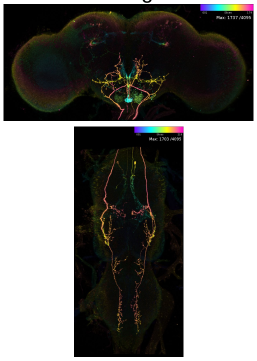

SS05129 - DNg28

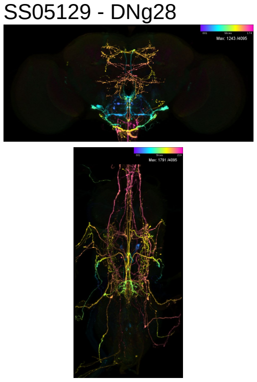

SS45805 - DNg28

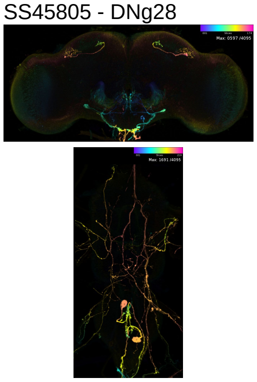

SS57822 - DNg29

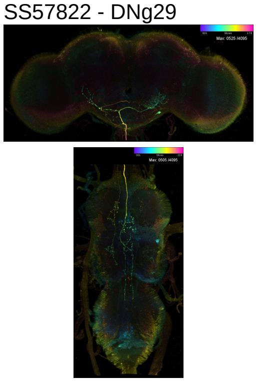

SS83529 - DNg29

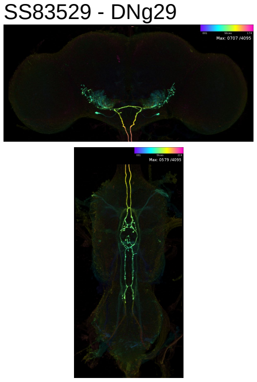

SS30397 - DNg29, DNg72

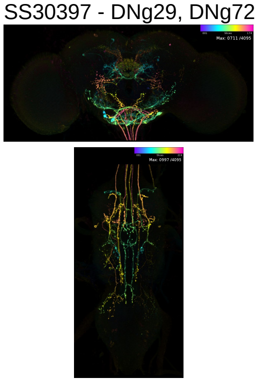

SS83514 - DNg29, DNp27

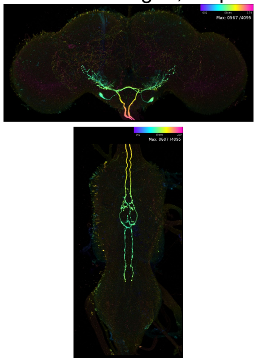

SS49825 - DNg31

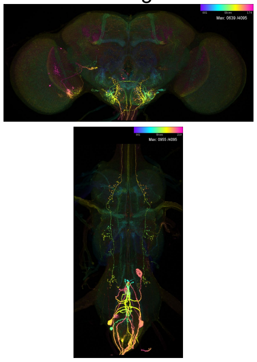

SS49838 - DNg31

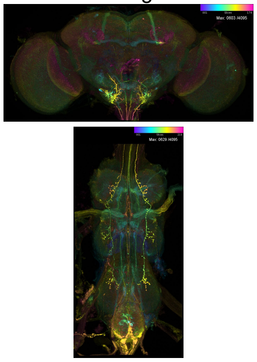

SS53429 - DNg31, DNg59, DNg113

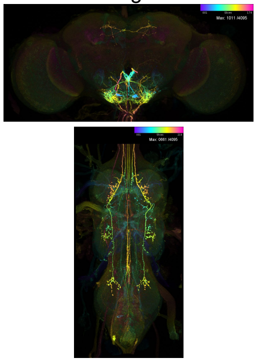

SS81943 - DNg31, DNg59, DNg113

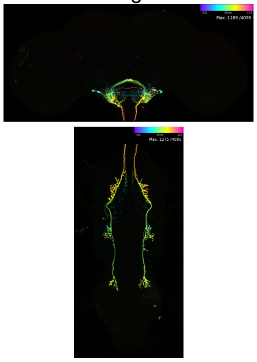

SS81942 - DNg31, DNg79

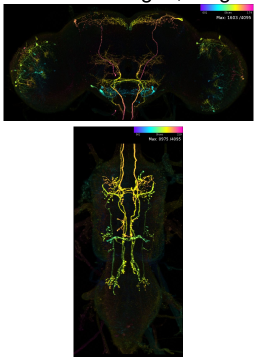

SS64068 - DNg33

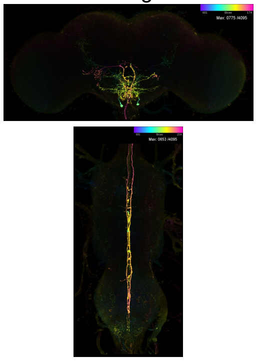

SS53439 - DNg34

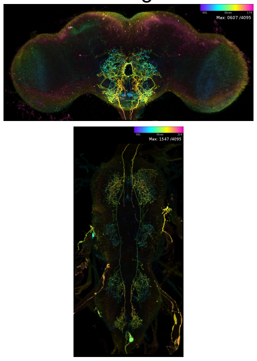

SS58292 - DNg34

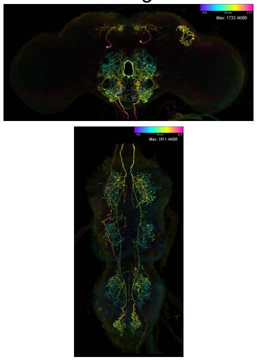

SS59935 - DNg34

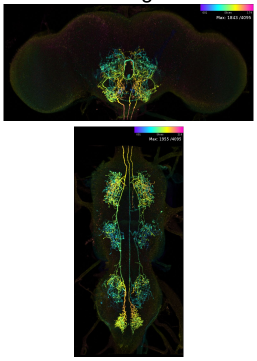

SS72208 - DNg34

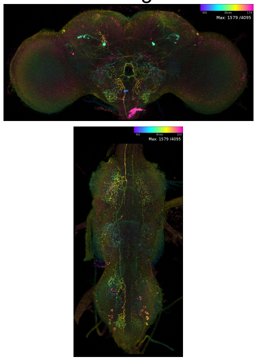

SS72207 - DNg34, DNg114

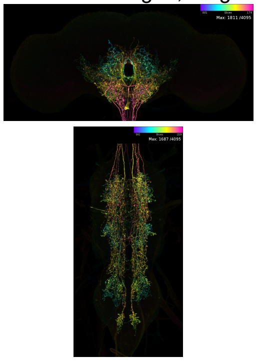

SS57787 - DNg34, DNp33

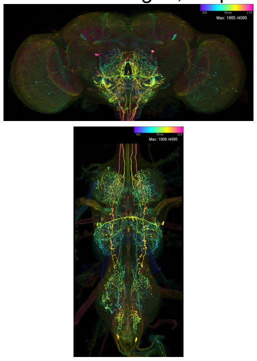

SS57828 - DNg35

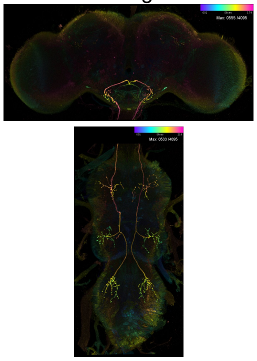

SS63217 - DNg35

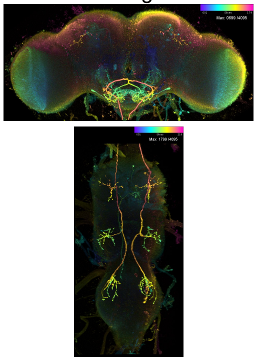

SS63264 - DNg35

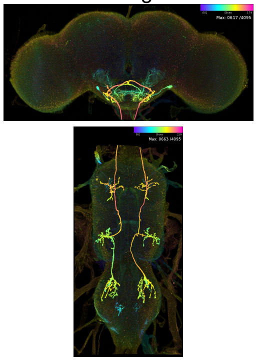

SS63305 - DNg35

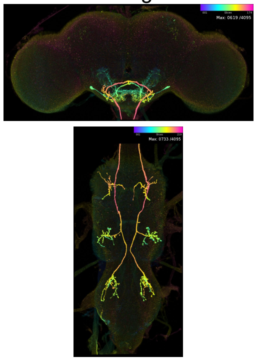

SS74458 - DNg35

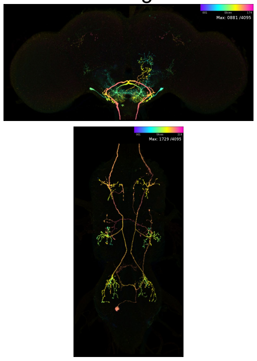

SS92268 - DNg36

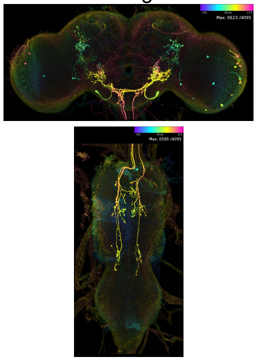

SS63295 - DNg37

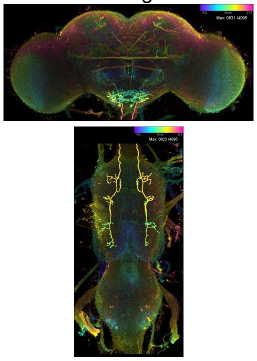

SS63296 - DNg37

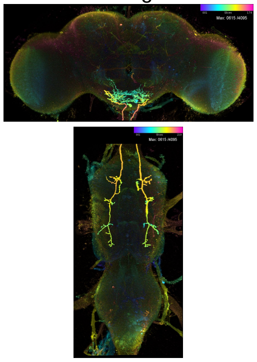

SS72975 - DNg37

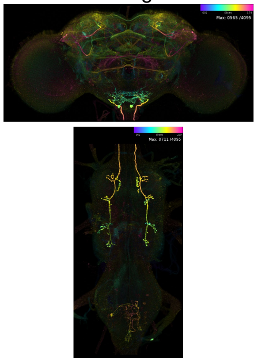

SS73035 - DNg37

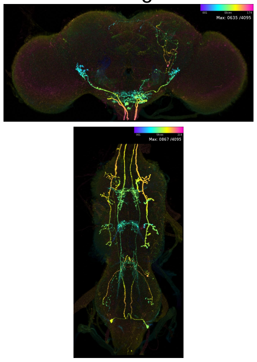

SS73420 - DNg37

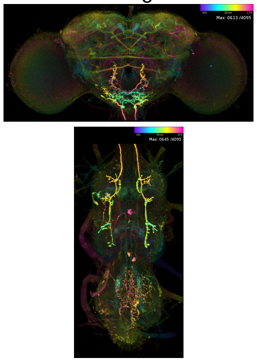

SS73421 - DNg37

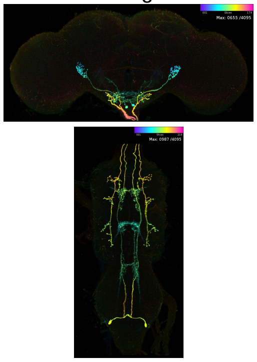

SS63737 - DNg38

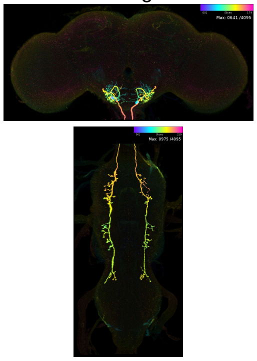

SS57737 - DNg39

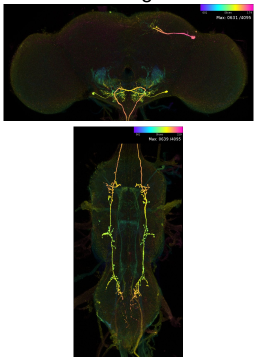

SS64882 - DNg39

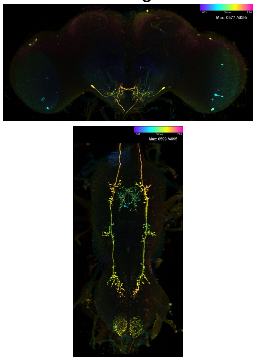

SS70644 - DNg39

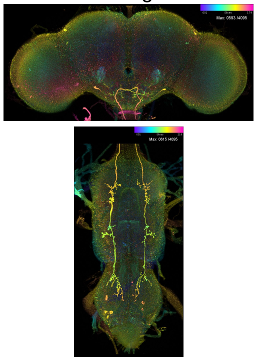

SS70655 - DNg39

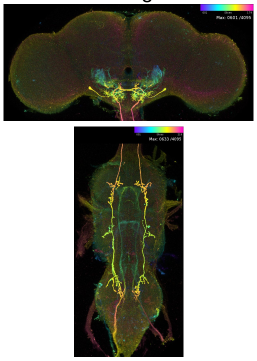

SS70659 - DNg39

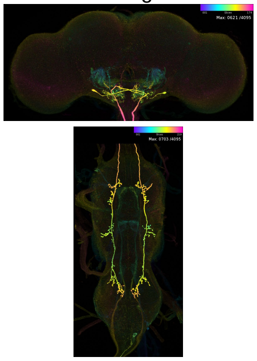

SS56718 - DNg41

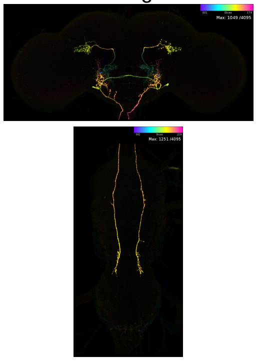

SS72087 - DNg41

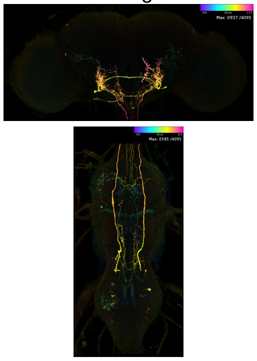

SS64896 - DNg43

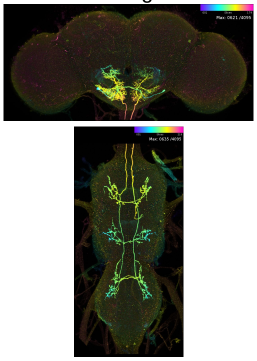

SS66182 - DNg43

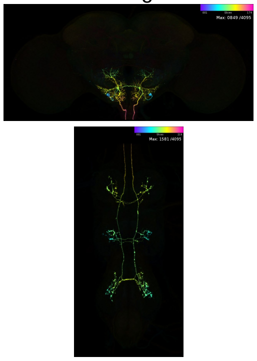

SS66224 - DNg43

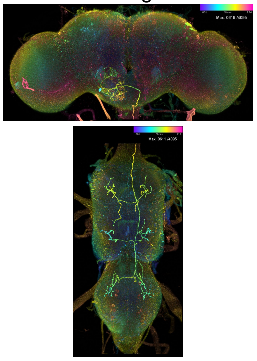

SS68371 - DNg43

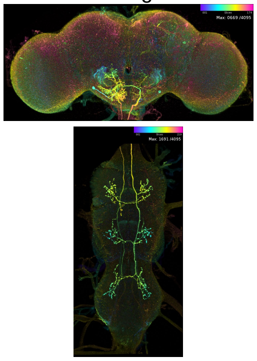

SS53380 - DNg45

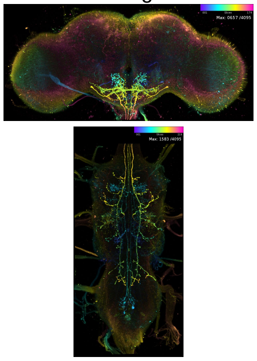

SS64037 - DNg45

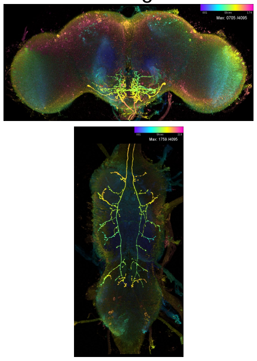

SS64584 - DNg45

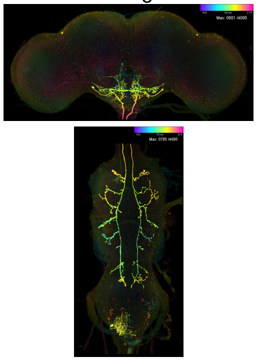

SS64640 - DNg45

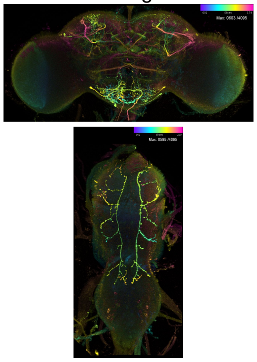

SS64641 - DNg45

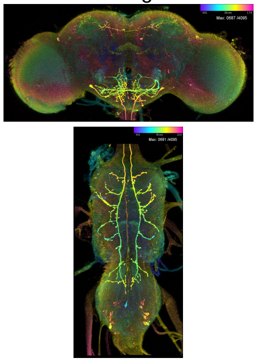

SS64642 - DNg45

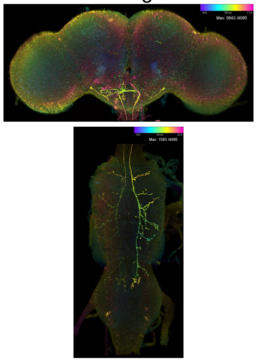

SS70544 - DNg45

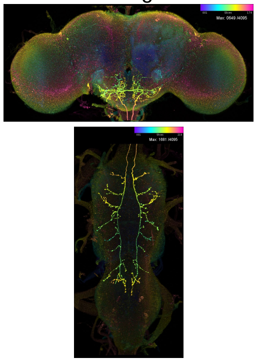

SS81824 - DNg45

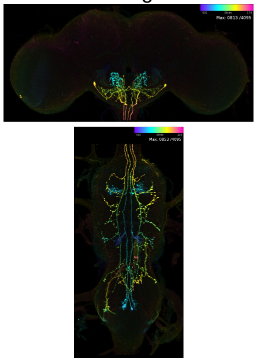

SS81829 - DNg45

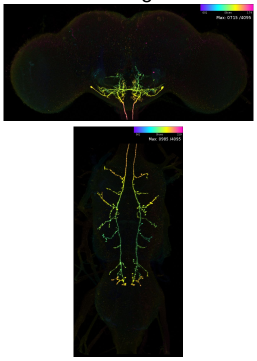

SS81914 - DNg45

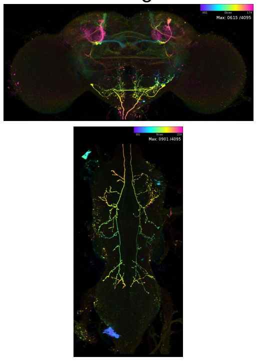

SS90832 - DNg45

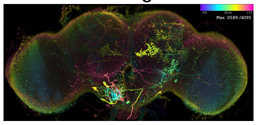

SS88050 - DNg46

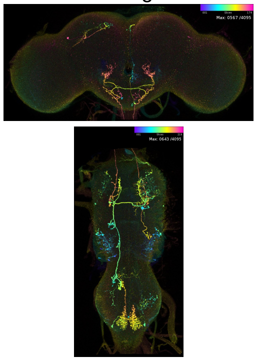

SS67049 - DNg48

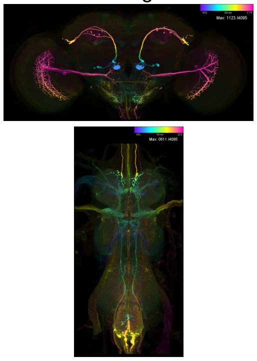

SS67173 - DNg48

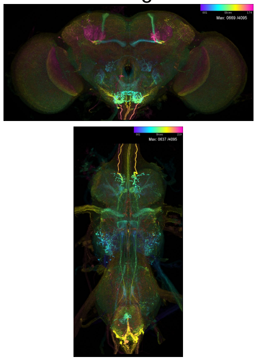

SS94158 - DNg48

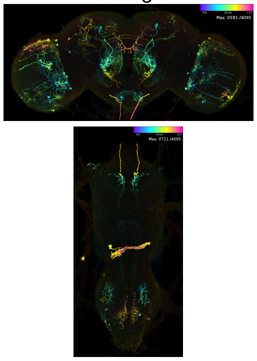

SS66214 - DNg49

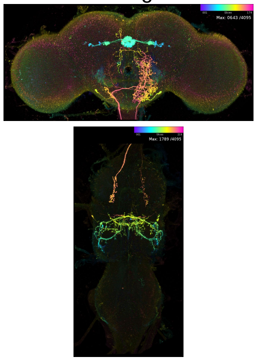

SS67126 - DNg49

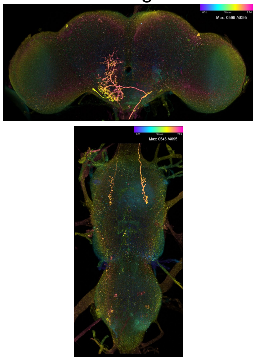

SS68377 - DNg49

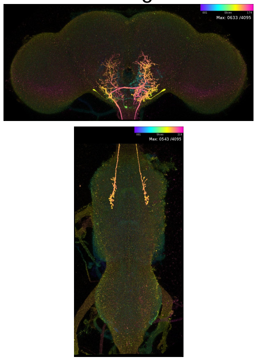

SS83462 - DNg49

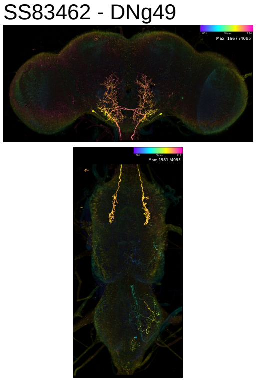

SS83552 - DNg49

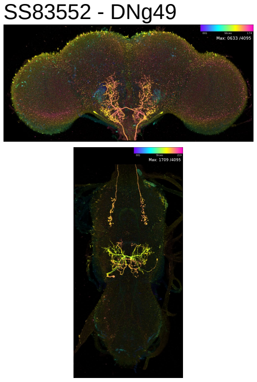

SS83554 - DNg49

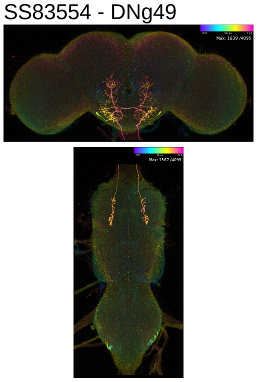

SS83573 - DNg49

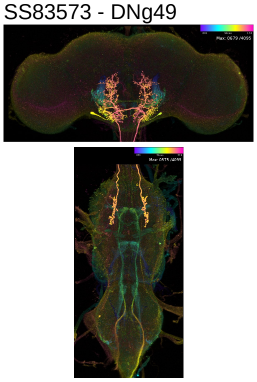

SS83625 - DNg49

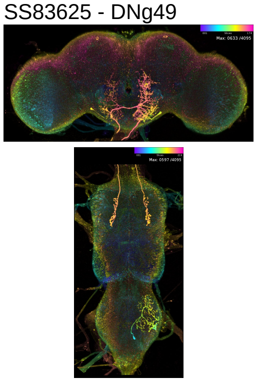

SS83627 - DNg49

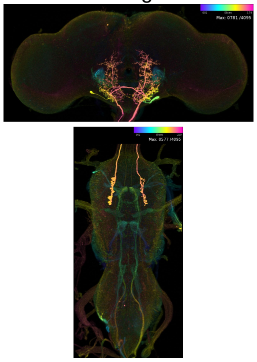

SS64577 - DNg50

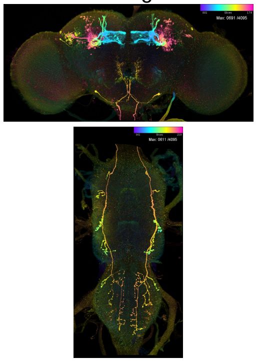

SS83468 - DNg50

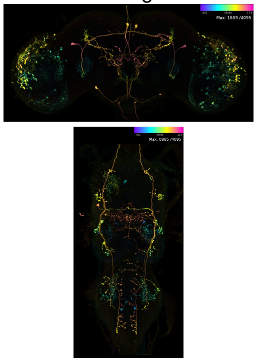

SS95552 - DNg50

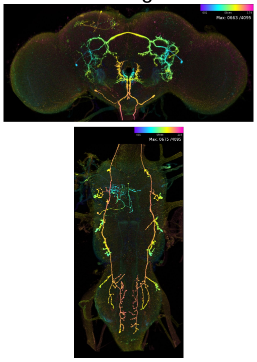

SS95555 - DNg50

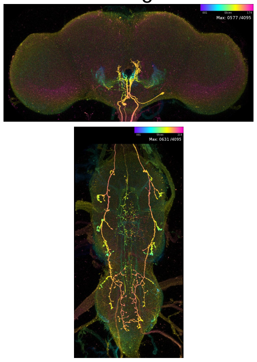

SS81873 - DNg51

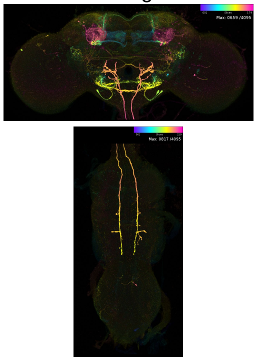

SS83497 - DNg51

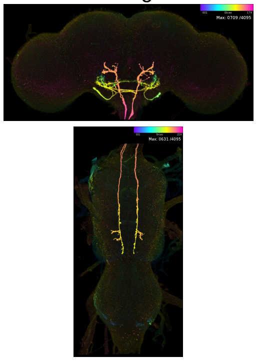

SS83634 - DNg51

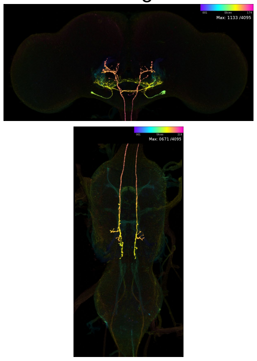

SS88178 - DNg51

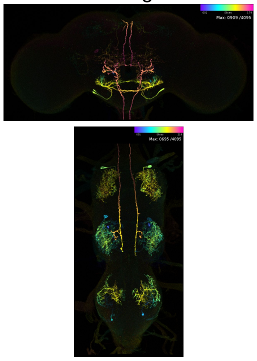

SS88179 - DNg51

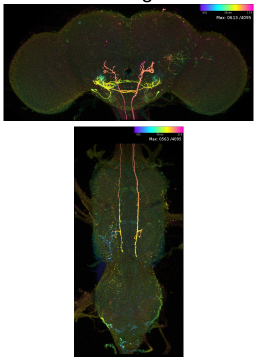

SS88180 - DNg51

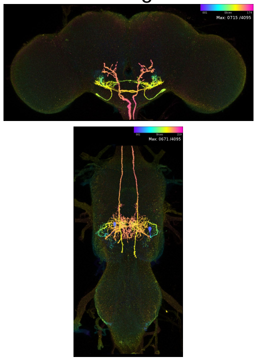

SS67133 - DNg52

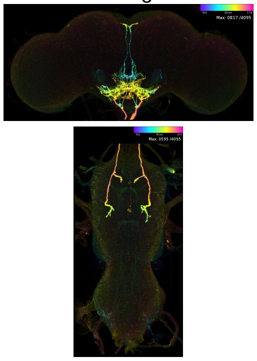

SS92766 - DNg53

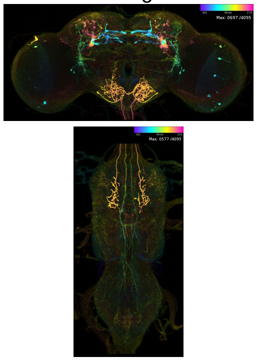

SS81934 - DNg57

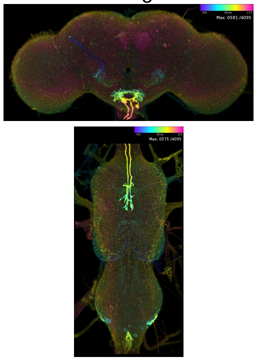

SS81935 - DNg57

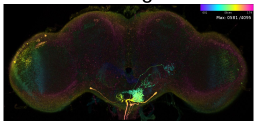

SS67102 - DNg62

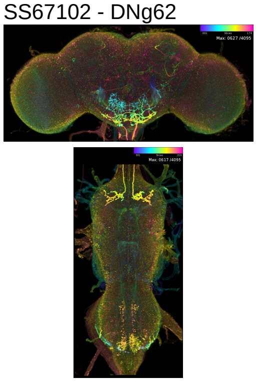

SS81883 - DNg62

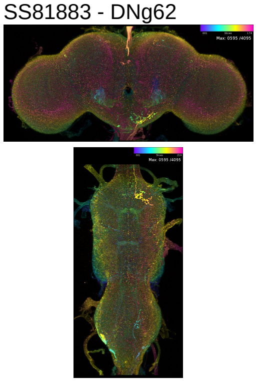

SS62943 - DNg63

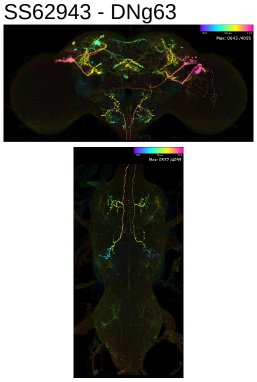

SS74446 - DNg65

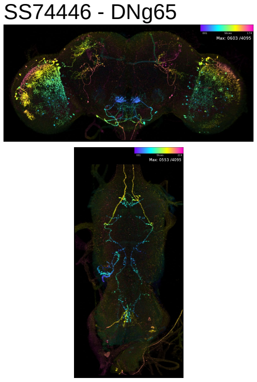

SS64570 - DNg69

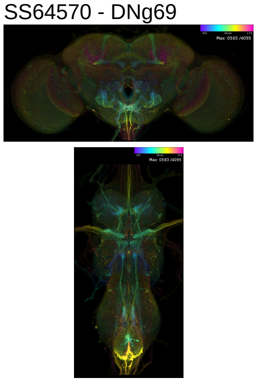

SS64627 - DNg69

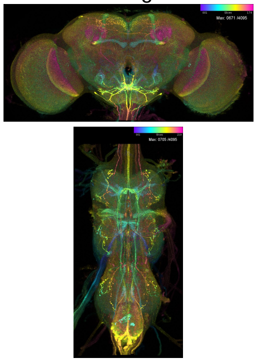

SS64662 - DNg69

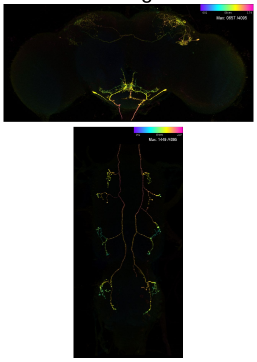

SS64664 - DNg69

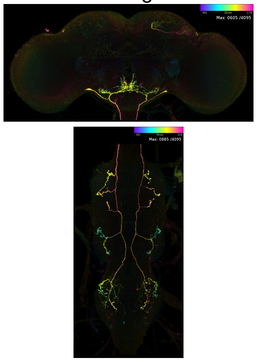

SS64675 - DNg69

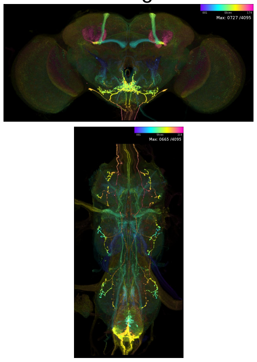

SS64676 - DNg69

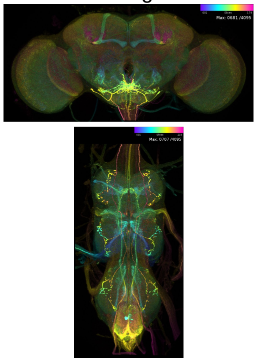

SS85819 - DNg69

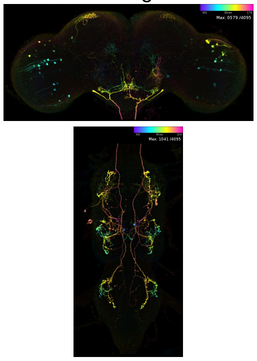

SS92183 - DNg69

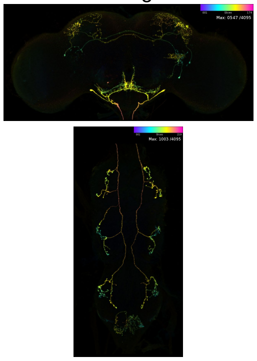

SS92196 - DNg69

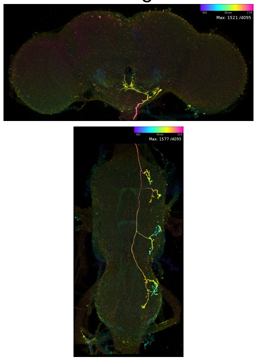

SS92200 - DNg69

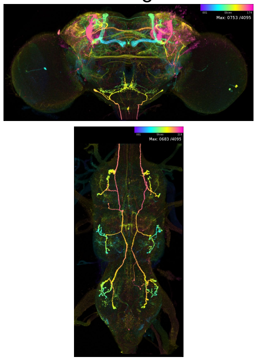

SS92652 - DNg69

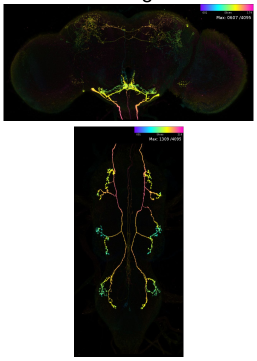

SS92658 - DNg69

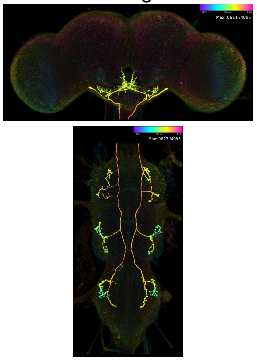

SS92852 - DNg69

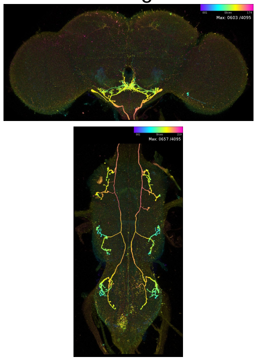

SS92895 - DNg69

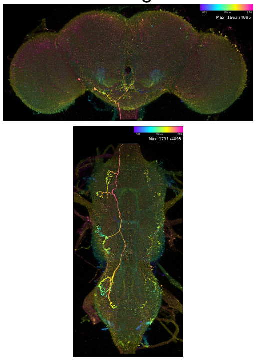

SS93272 - DNg69

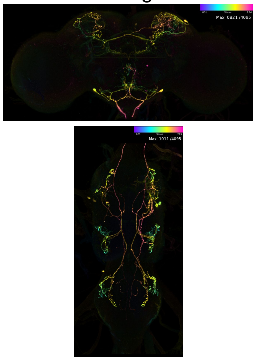

SS53371 - DNg72

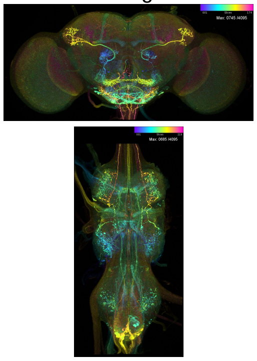

SS66234 - DNg72

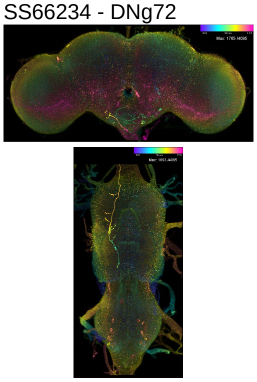

SS66262 - DNg72

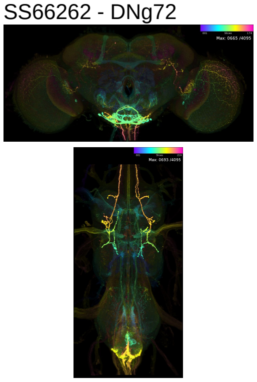

SS81946 - DNg72

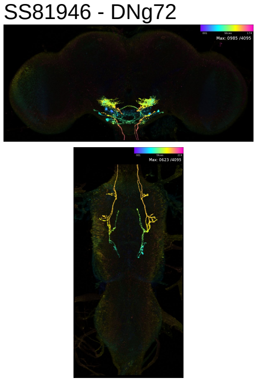

SS88190 - DNg74

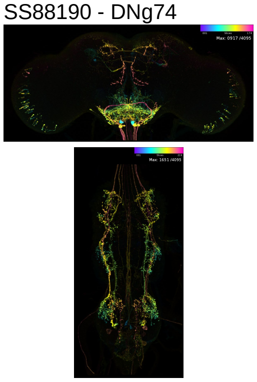

SS90782 - DNg74

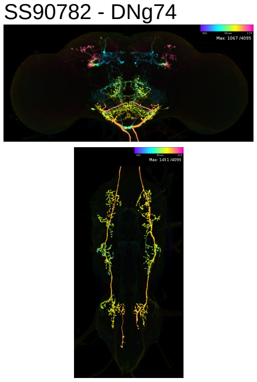

SS67217 - DNg76

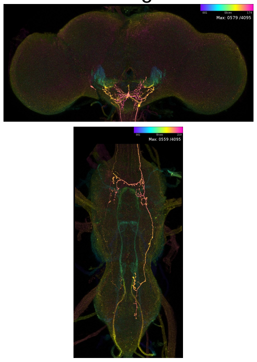

SS72267 - DNg76

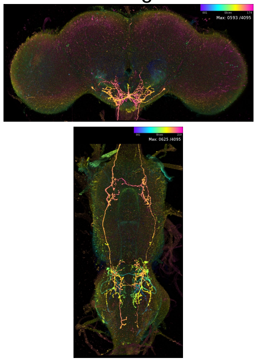

SS93746 - DNg76

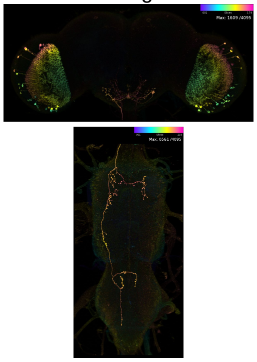

SS93751 - DNg76

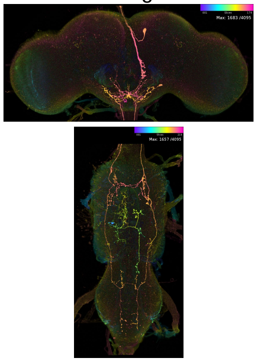

SS81928 - DNg76, DNp19

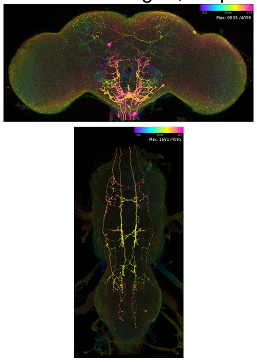

SS31716 - DNg77

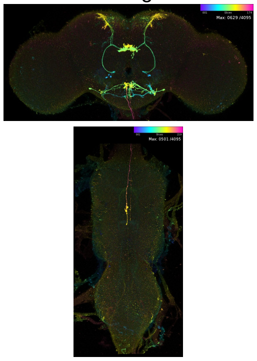

SS86249 - DNg77

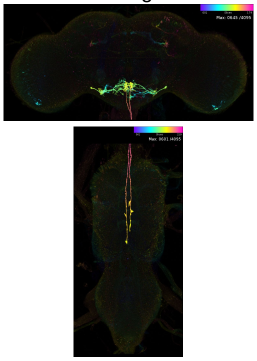

SS92649 - DNg77

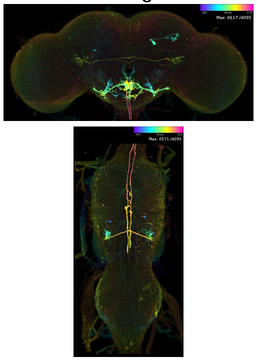

SS92893 - DNg77

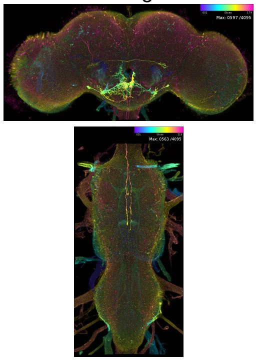

SS93646 - DNg77

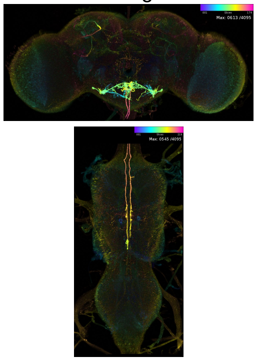

SS13273 - DNg78

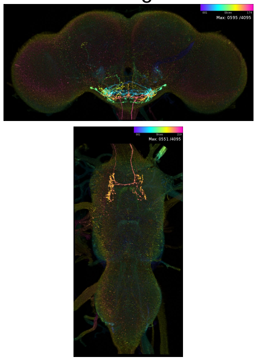

SS13337 - DNg78

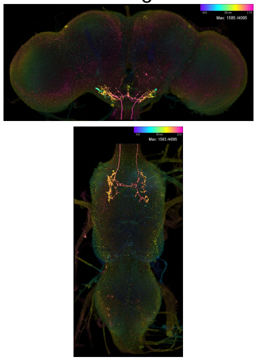

SS66261 - DNg78

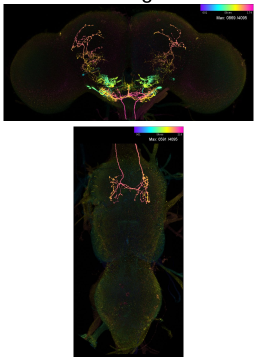

SS67141 - DNg78

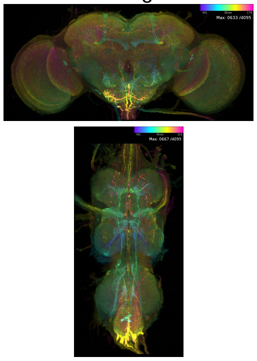

SS72084 - DNg78

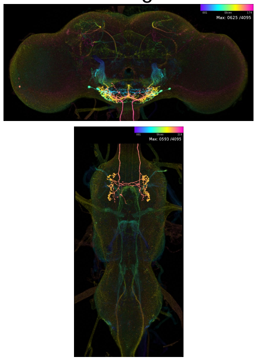

SS73431 - DNg78

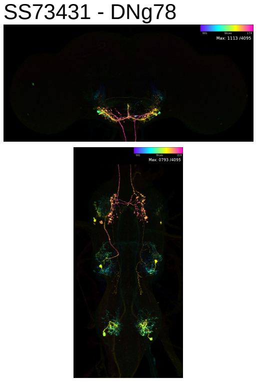

SS95049 - DNg78

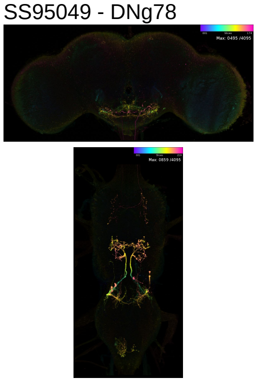

SS95074 - DNg78

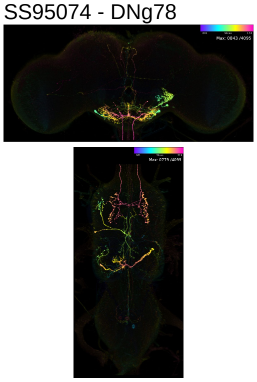

SS57731 - DNg80

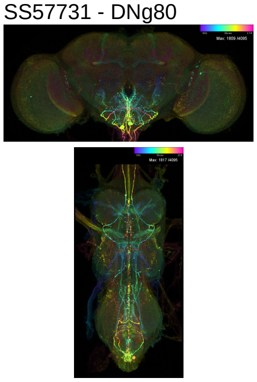

SS57781 - DNg80

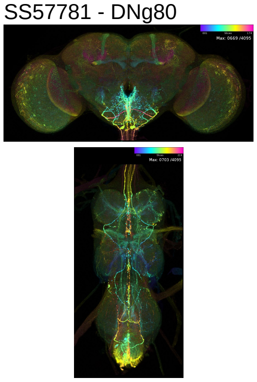

SS73484 - DNg80

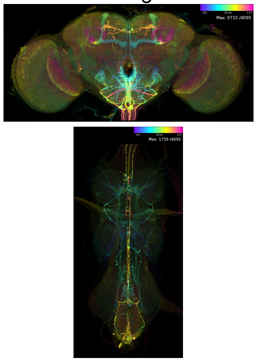

SS93643 - DNg80

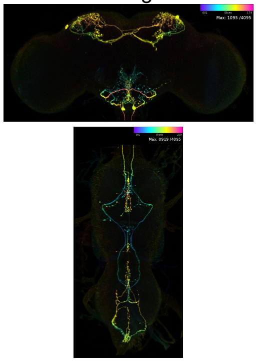

SS56192 - DNg80, DNp27

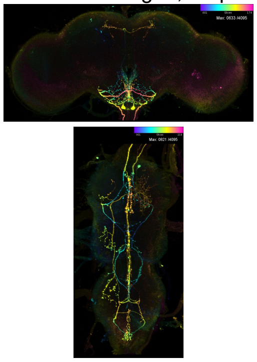

SS95034 - DNg82, DNg91

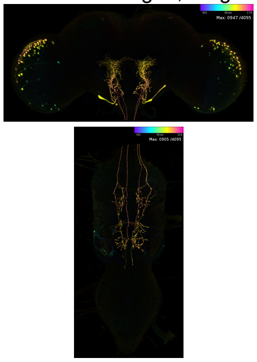

SS95406 - DNg84

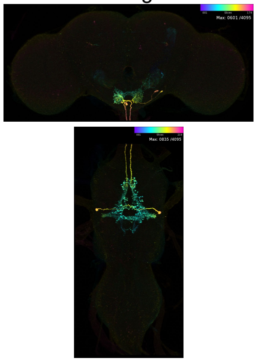

SS64051 - DNg87

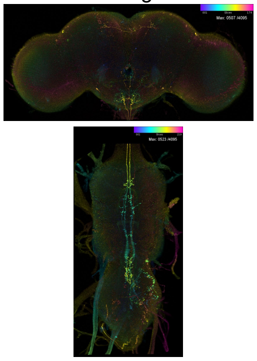

SS94997 - DNg91

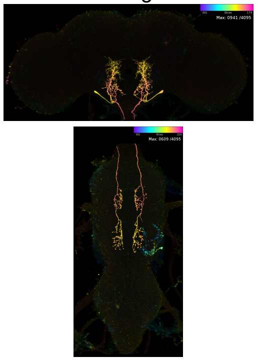

SS57751 - DNg92

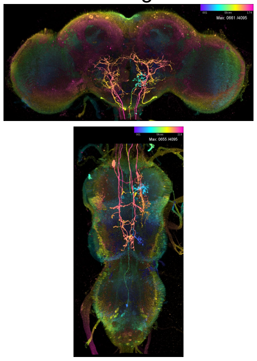

SS65576 - DNg92

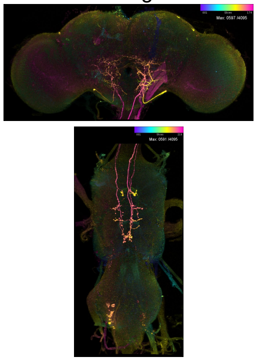

SS85855 - DNg93

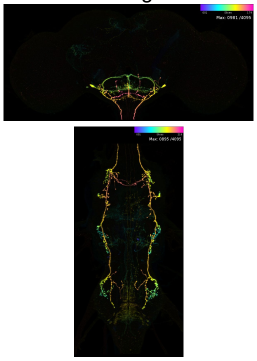

SS86207 - DNg93

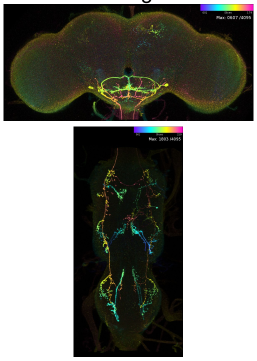

SS88170 - DNg93

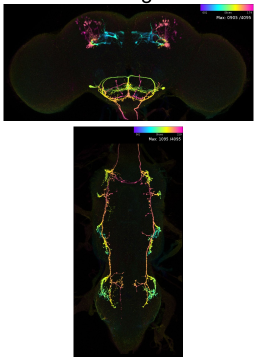

SS90781 - DNg93

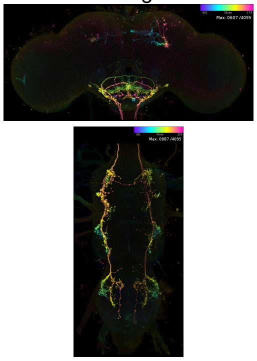

SS92255 - DNg93

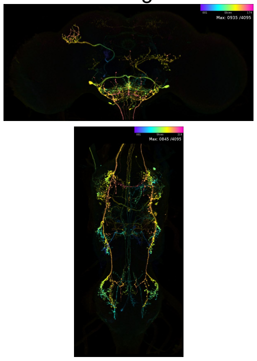

SS92312 - DNg93

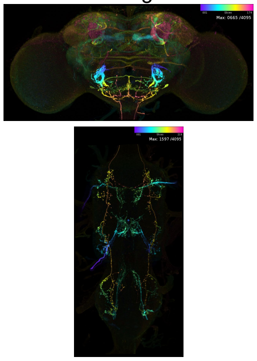

SS93715 - DNg93

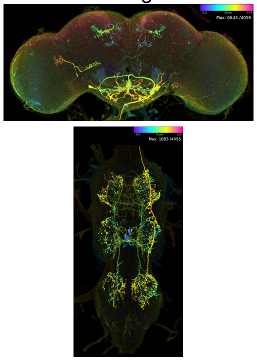

SS95522 - DNg93

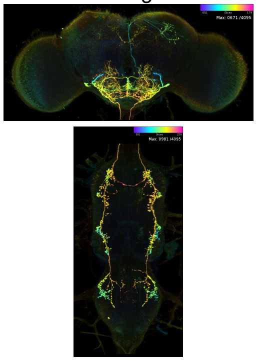

SS95523 - DNg93

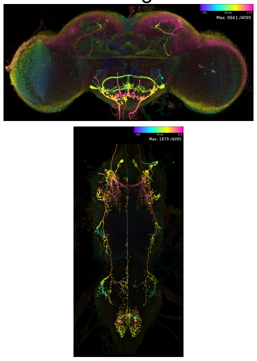

SS88151 - DNg100, DNg108

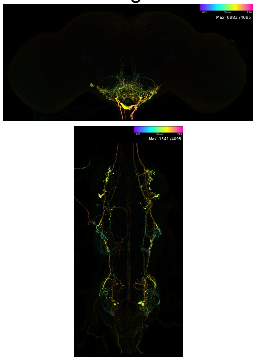

SS67050 - DNg100, DNp41, DNp51

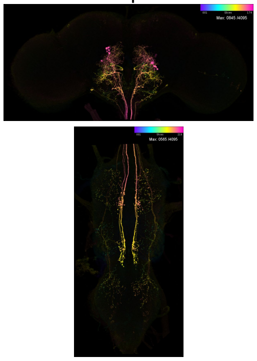

SS52378 - DNg106

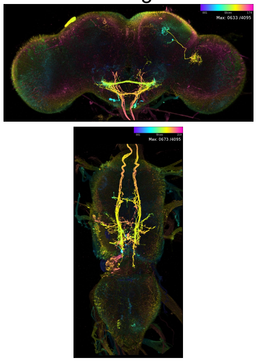

SS98615 - DNg106

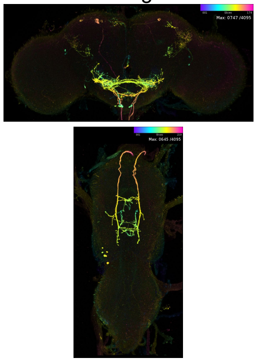

SS98621 - DNg106

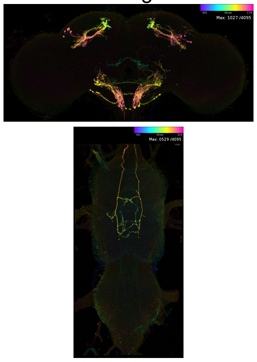

SS98622 - DNg106

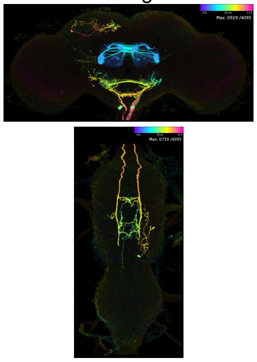

SS64046 - DNg112

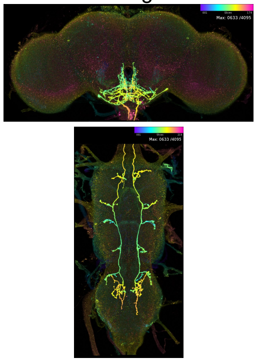

SS64047 - DNg112

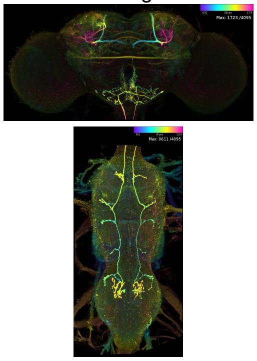

SS64582 - DNg112

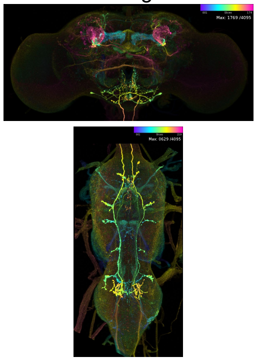

SS64648 - DNg112

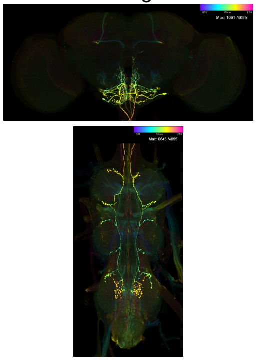

SS57831 - DNg114

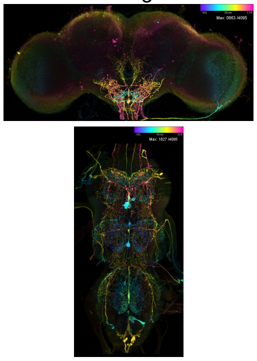

SS55962 - DNg115

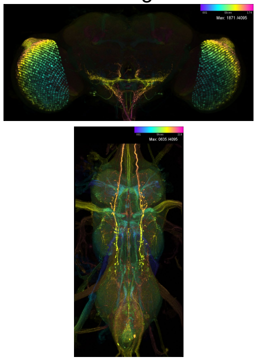

SS67175 - DNg116

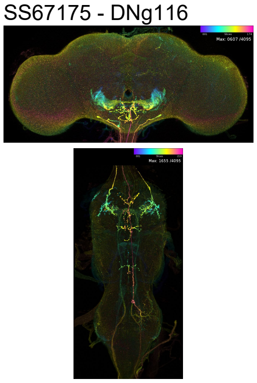

SS73466 - DNg117

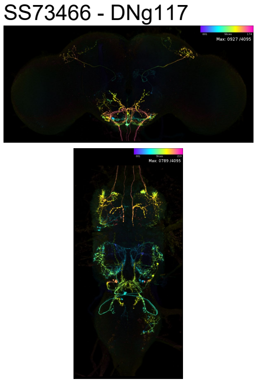

SS88171 - DNg118

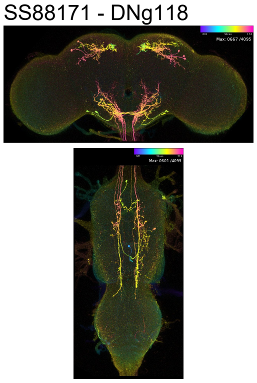

SS94744 - DNg118

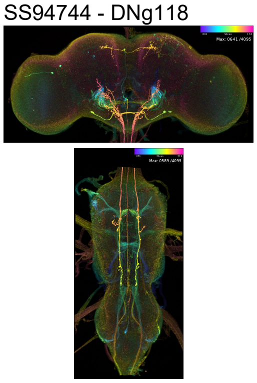

SS92768 - DNg119

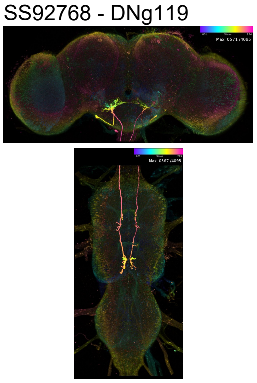

SS67124 - DNg119, DNg126

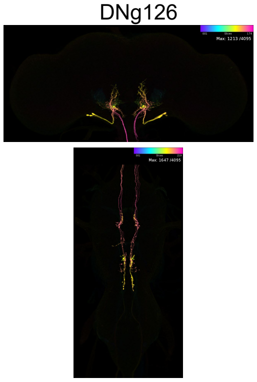

SS67162 - DNg119, DNg126

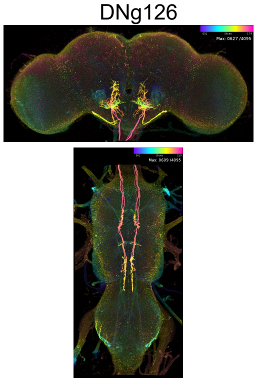

SS93264 - DNg120

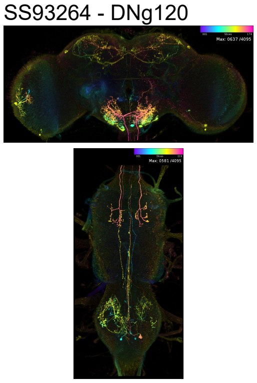

SS96088 - DNg121

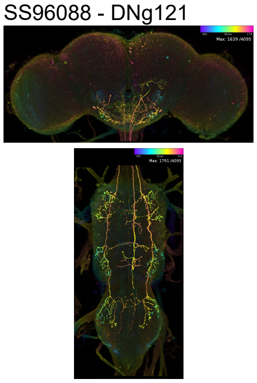

SS92798 - DNg122

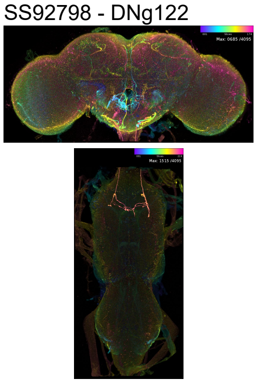

SS96120 - DNg123

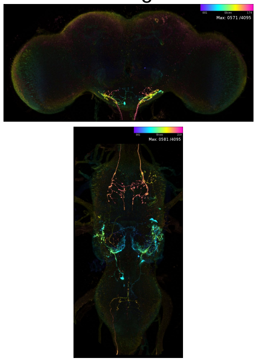

SS67209 - DNg126

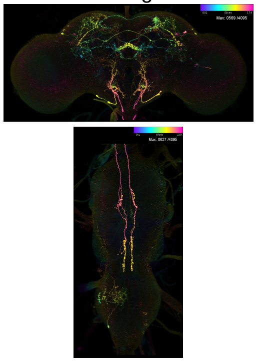

SS67129 - DNp01, DNp74

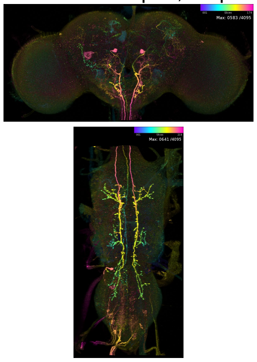

SS49056 - DNp02, DNp06

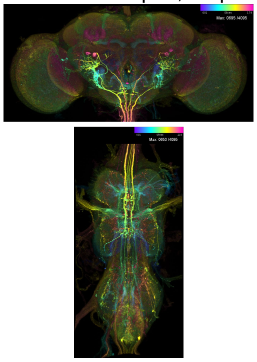

SS56230 - DNp03

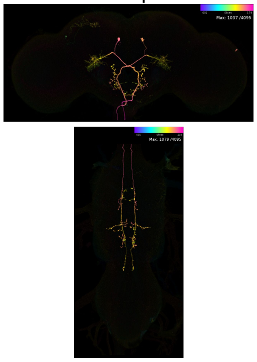

SS95781 - DNp07

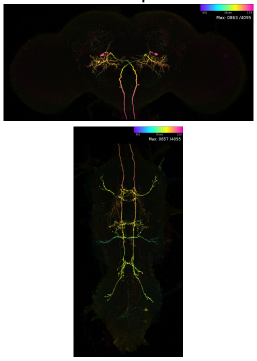

SS72971 - DNp08

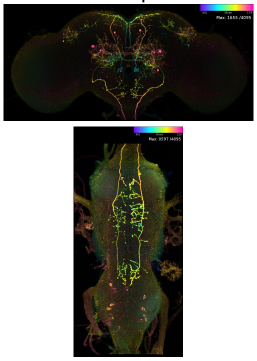

SS74433 - DNp08

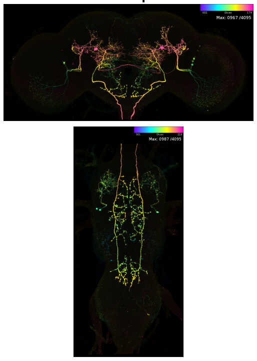

SS74436 - DNp08

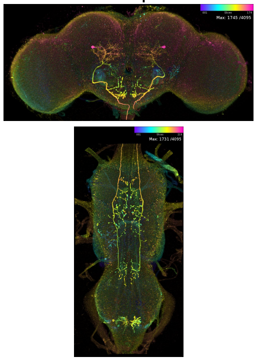

SS74438 - DNp08

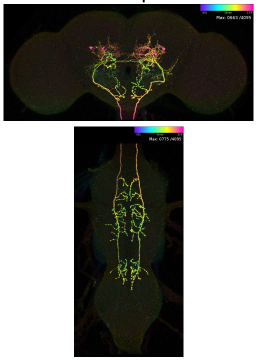

SS74441 - DNp08

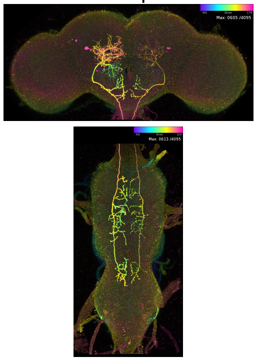

SS81930 - DNp08

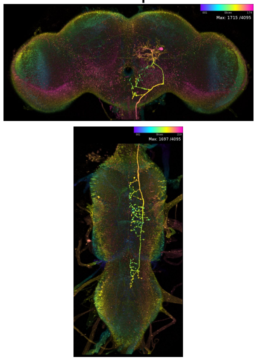

SS83512 - DNp08

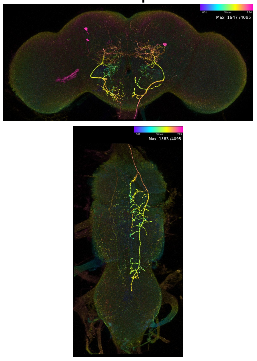

SS83515 - DNp08

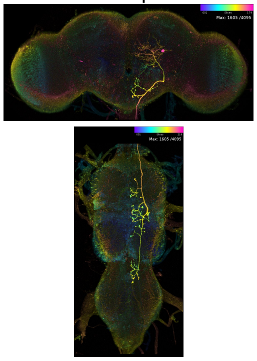

SS83614 - DNp08

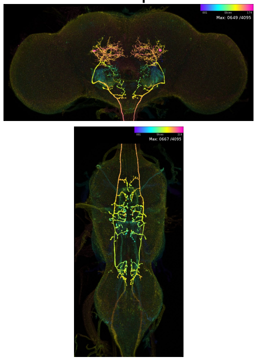

SS83628 - DNp08

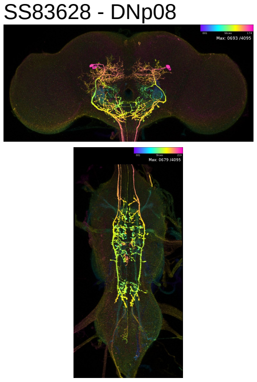

SS45758 - DNp09

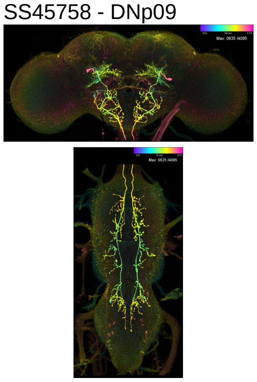

SS04525 - DNp11

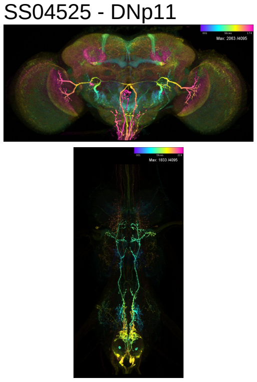

SS49051 - DNp11

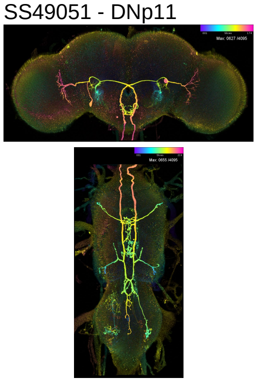

SS95373 - DNp11

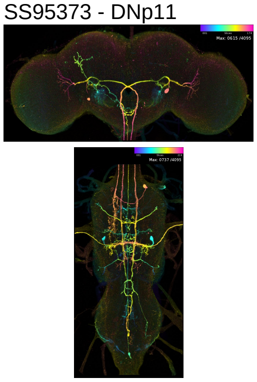

SS81905 - DNp12

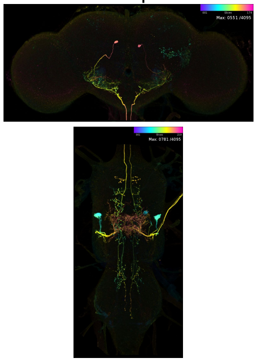

SS56181 - DNp13

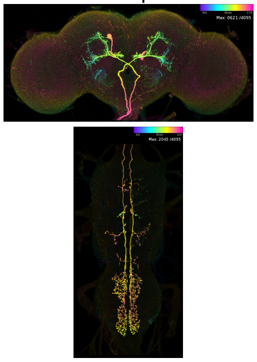

SS92713 - DNp13

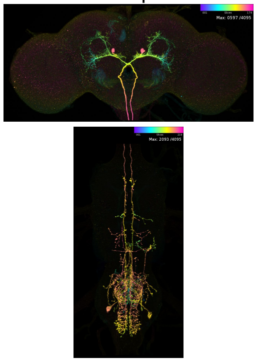

SS57757 - DNp14

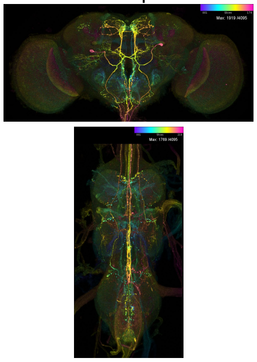

SS64605 - DNp14

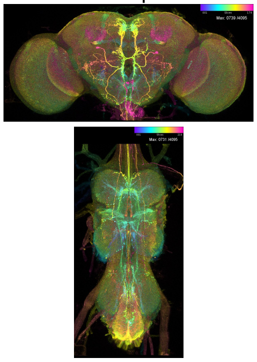

SS72989 - DNp14

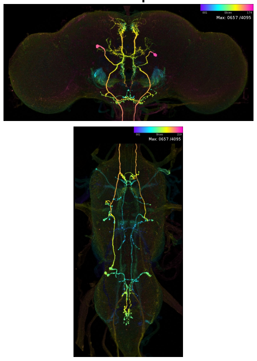

SS73011 - DNp14

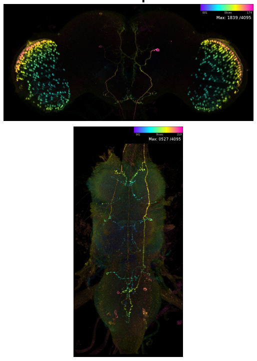

SS74396 - DNp14

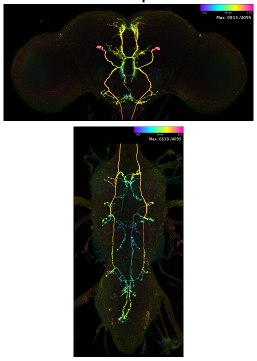

SS12653 - DNp15, DNp19

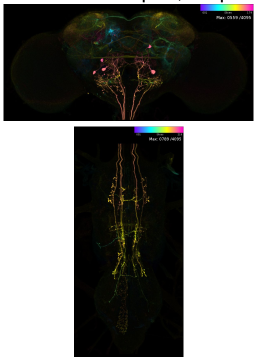

SS86301 - DNp16

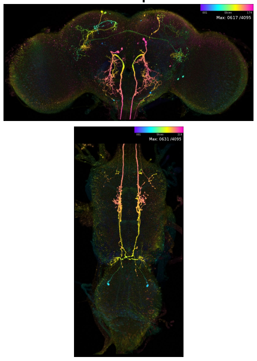

SS86302 - DNp16

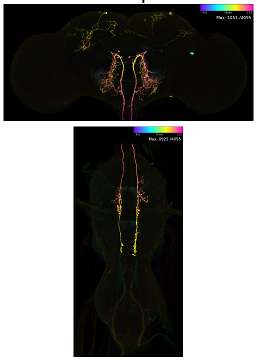

SS97219 - DNp16

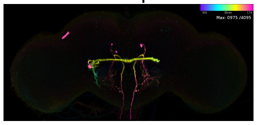

SS35956 - DNp17

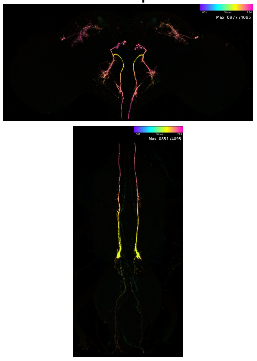

SS85849 - DNp17

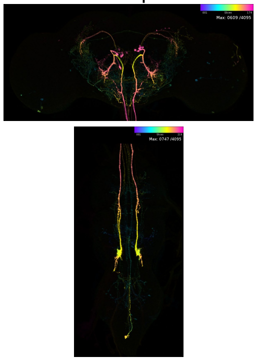

SS85858 - DNp17

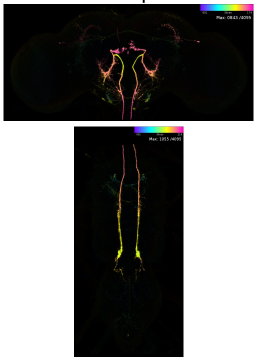

SS85873 - DNp17

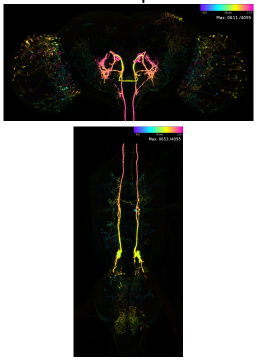

SS85878 - DNp17

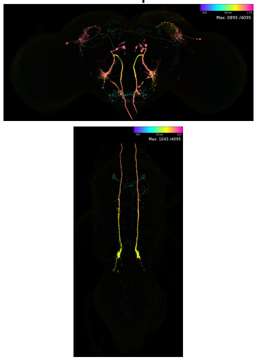

SS85883 - DNp17

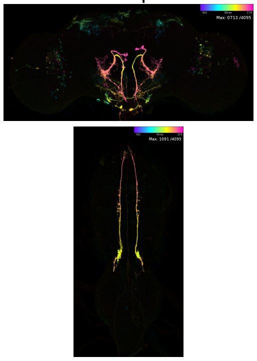

SS85903 - DNp17

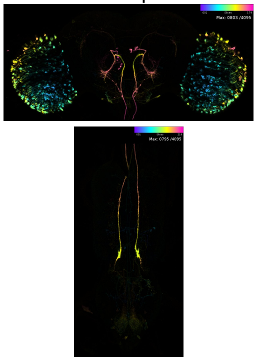

SS85912 - DNp17

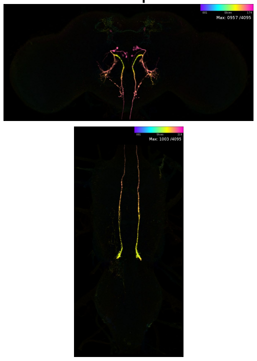

SS86184 - DNp17

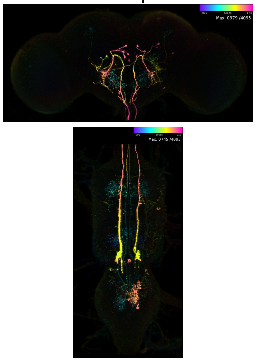

SS86186 - DNp17

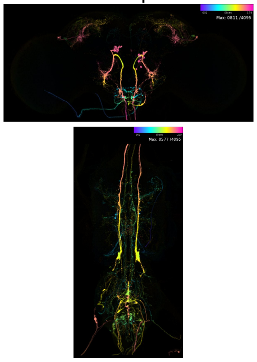

SS88183 - DNp17

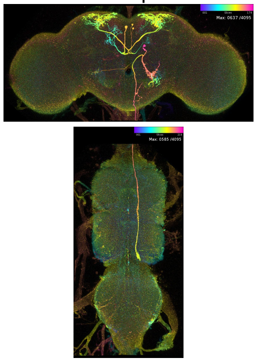

SS88197 - DNp17

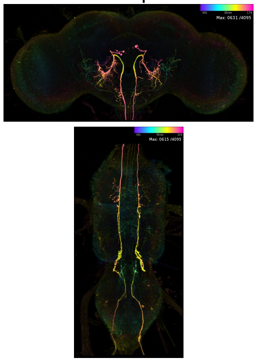

SS97180 - DNp17

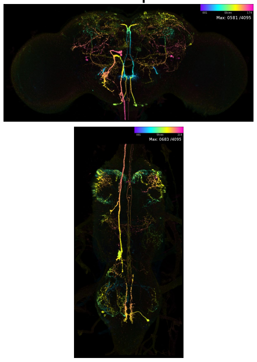

SS97181 - DNp17

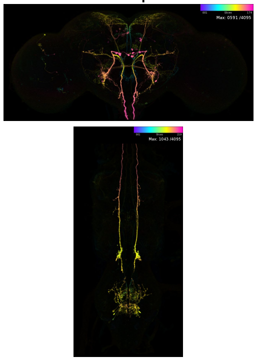

SS97206 - DNp17

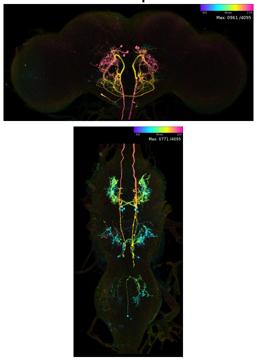

SS81965 - DNp19

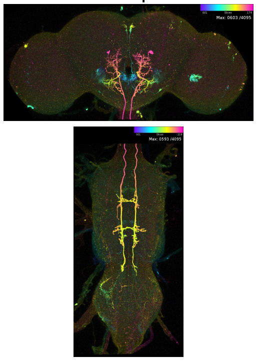

SS81956 - DNp20

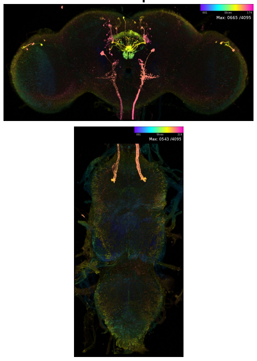

SS69336 - DNp22

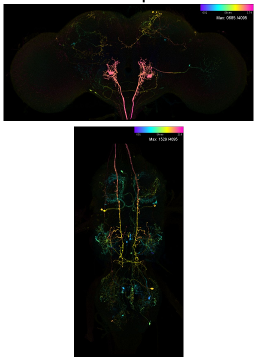

SS63716 - DNp24, DNp34

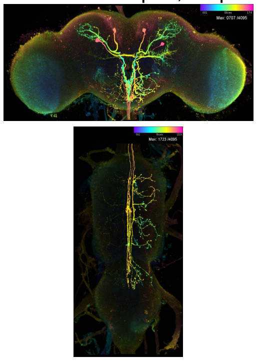

SS31564 - DNp26

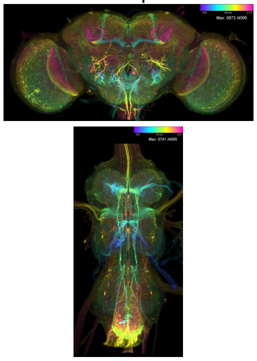

SS45741 - DNp26

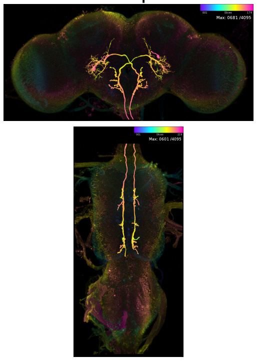

SS45762 - DNp26

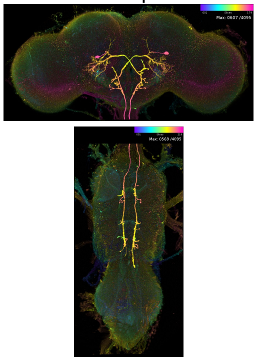

SS81818 - DNp26

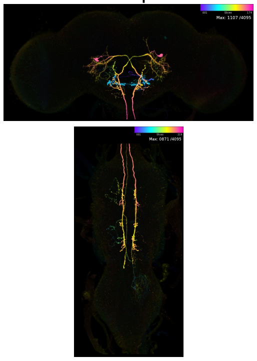

SS81821 - DNp26

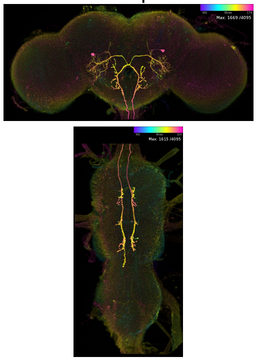

SS82248 - DNp26

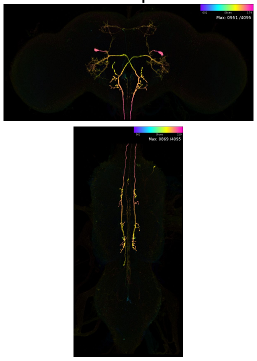

SS35976 - DNp30, DNp62

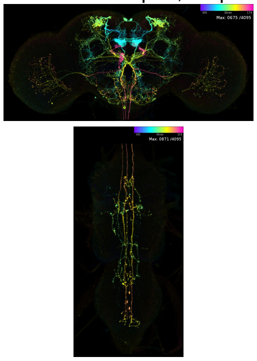

SS63665 - DNp30, DNp62

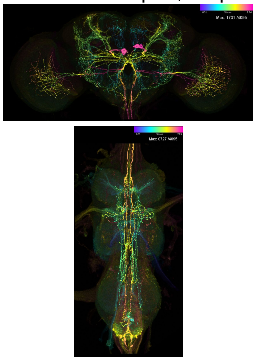

SS63699 - DNp30, DNp62

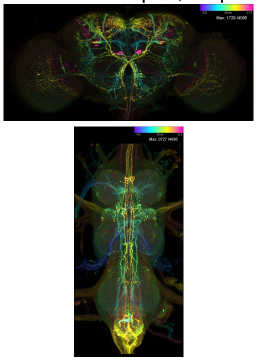

SS63700 - DNp30, DNp62

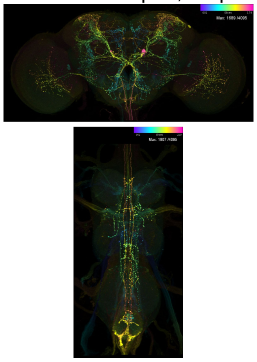

SS67122 - DNp30, DNp62

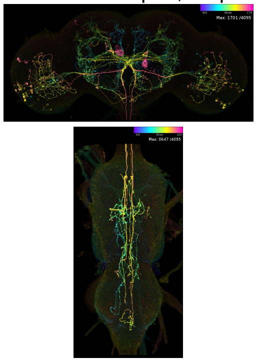

SS67193 - DNp30, DNp62

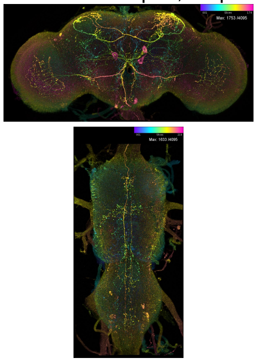

SS67194 - DNp30, DNp62

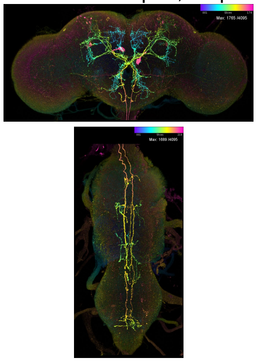

SS74148 - DNp30, DNp62

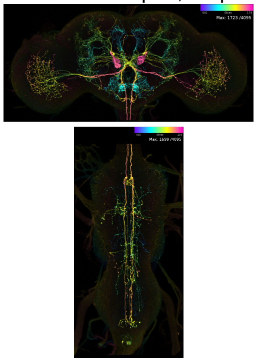

SS83544 - DNp30, DNp62

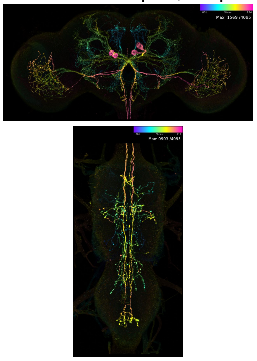

SS92748 - DNp30, DNp62

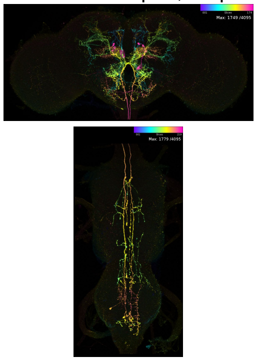

SS92762 - DNp30, DNp62

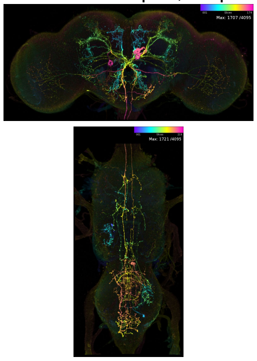

SS95362 - DNp31

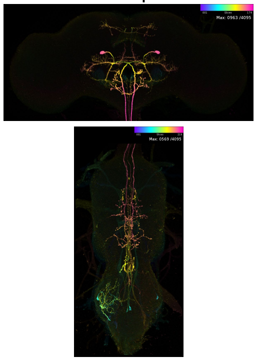

SS95379 - DNp31

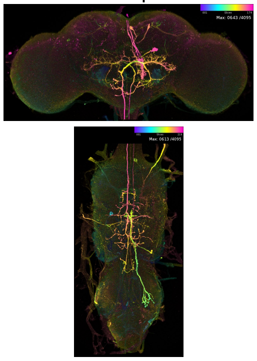

SS95508 - DNp31

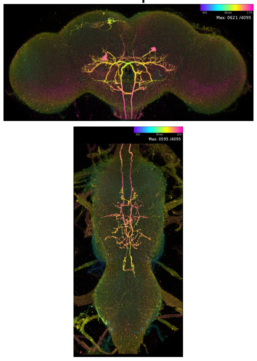

SS95510 - DNp31

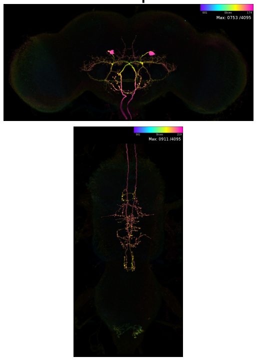

SS95780 - DNp31

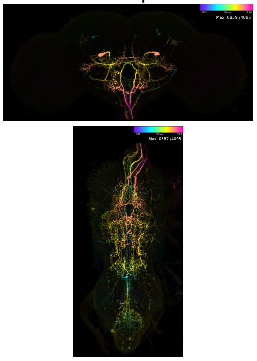

SS92858 - DNp32

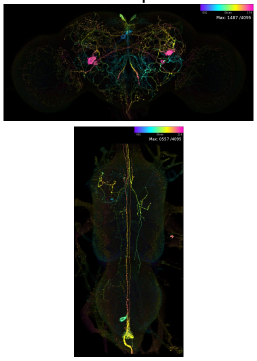

SS55960 - DNp33

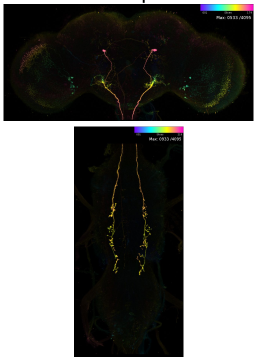

SS58316 - DNp33

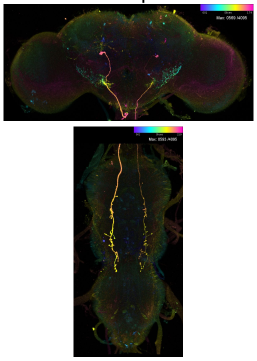

SS58318 - DNp33

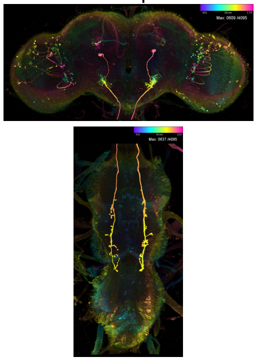

SS58354 - DNp33

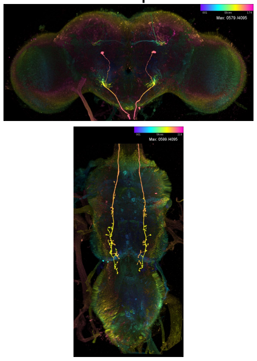

SS64070 - DNp33

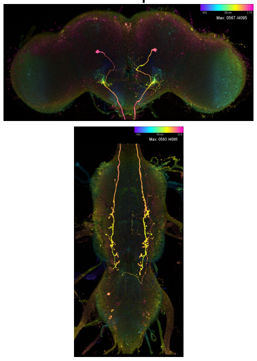

SS57026 - DNp34, DNp56

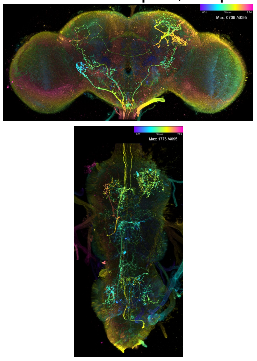

SS63993 - DNp35

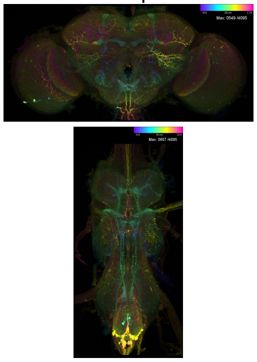

SS81912 - DNp37

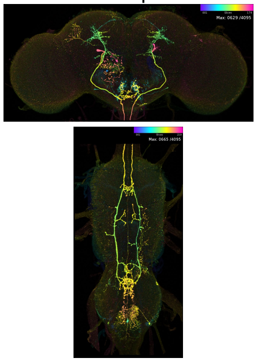

SS81962 - DNp37

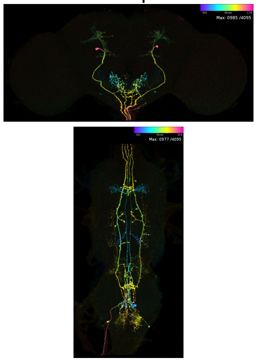

SS95006 - DNp37

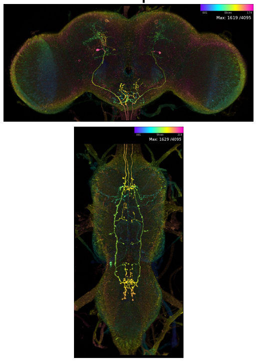

SS95004 - DNp37, DNp58

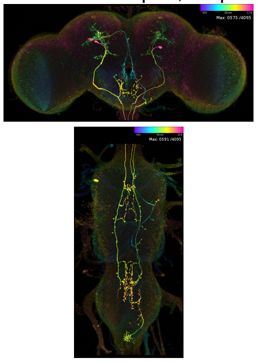

SS48301 - DNp38

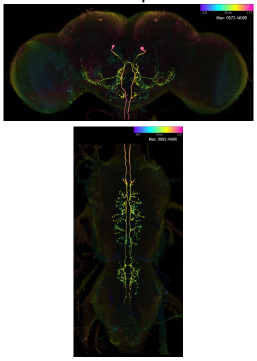

SS82242 - DNp41, DNp51

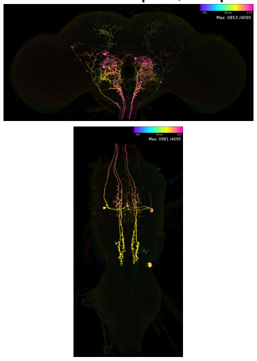

SS57049 - DNp42

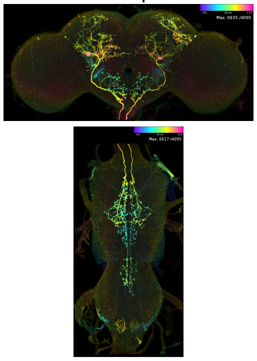

SS57742 - DNp42

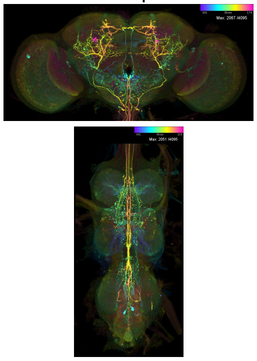

SS65441 - DNp43

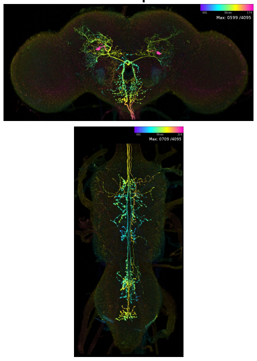

SS66153 - DNp44

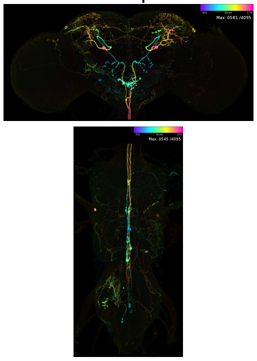

SS81831 - DNp45

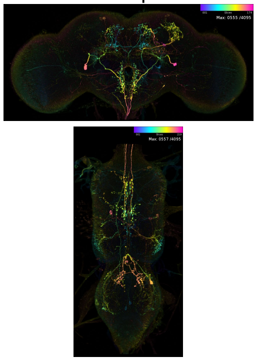

SS81903 - DNp46

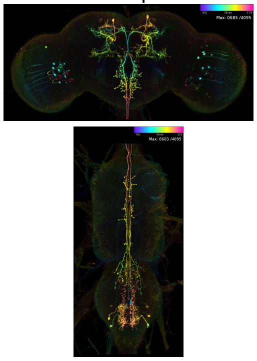

SS95399 - DNp46

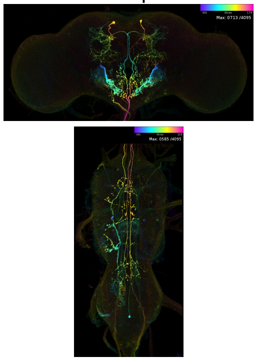

SS64601 - DNp48

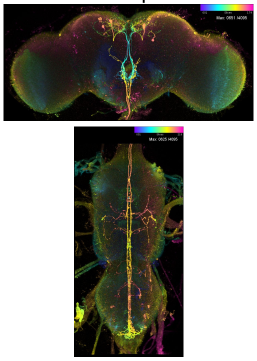

SS83557 - DNp48

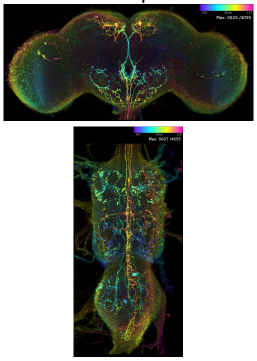

SS83607 - DNp48

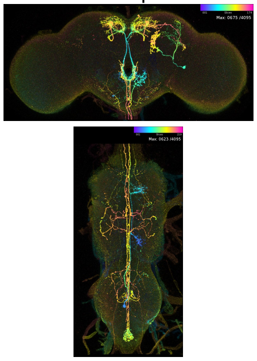

SS83608 - DNp48

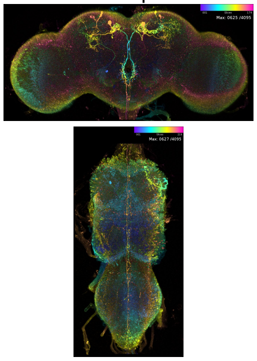

SS66163 - DNp49

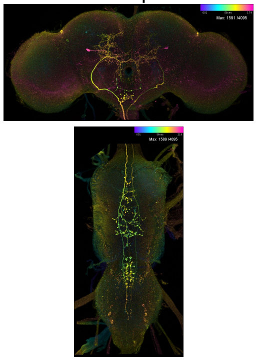

SS81968 - DNp51

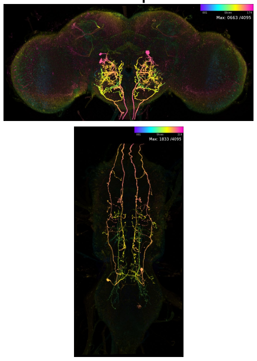

SS81969 - DNp51

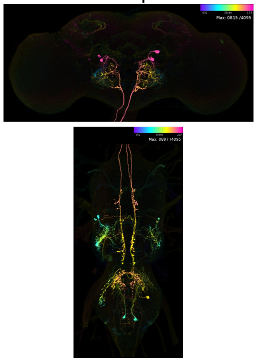

SS96062 - DNp54

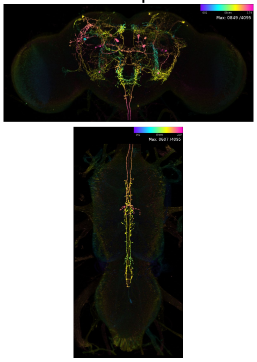

SS88245 - DNp56

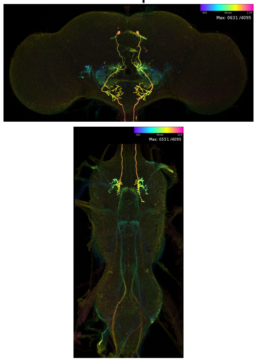

SS74386 - DNp57

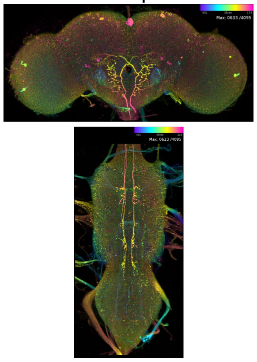

SS83495 - DNp62

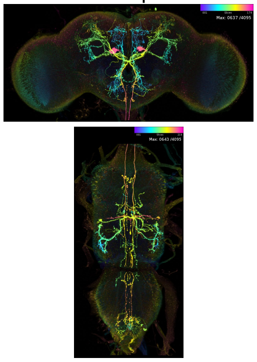

SS83496 - DNp62

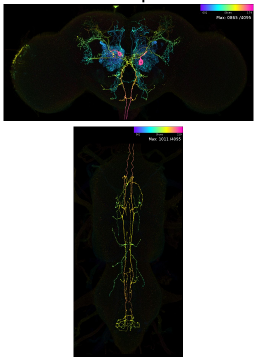

SS83545 - DNp62

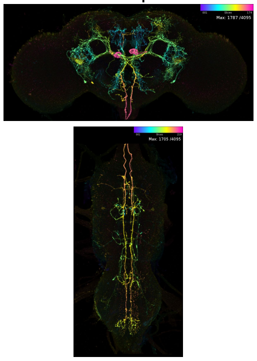

SS83568 - DNp62

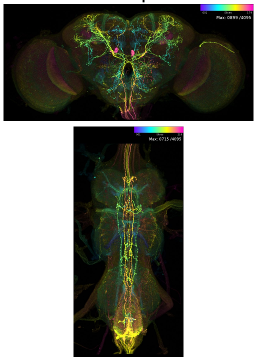

SS83615 - DNp62

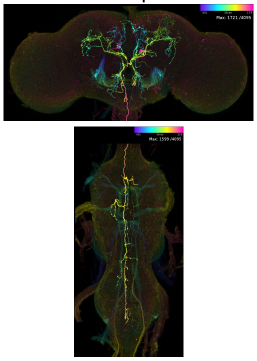

SS92749 - DNp62

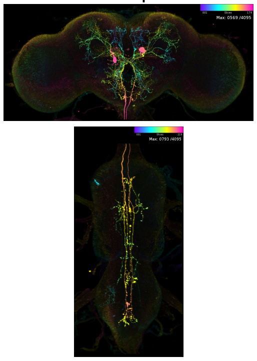

SS92910 - DNp62

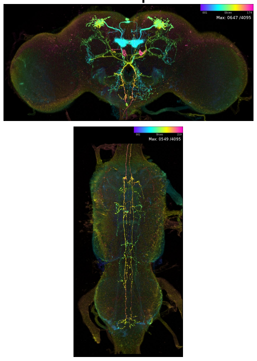

SS92911 - DNp62

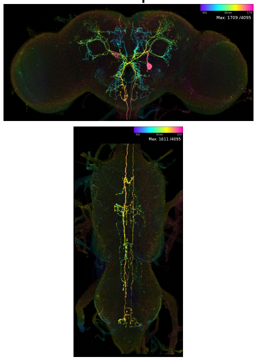

SS67048 - DNp65

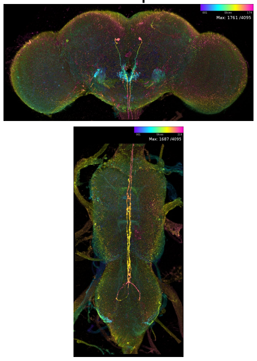

SS88242 - DNp65

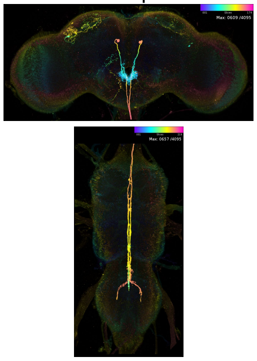

SS88246 - DNp65

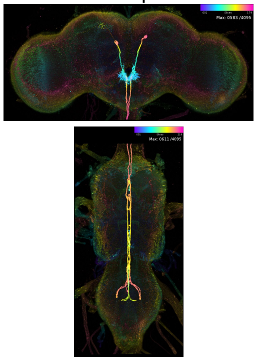

SS90829 - DNp71

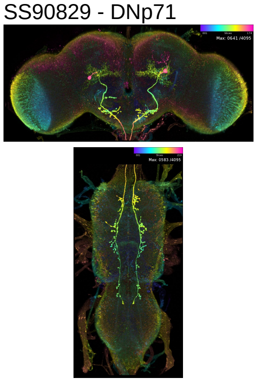

SS81847 - DNp71, DNp103

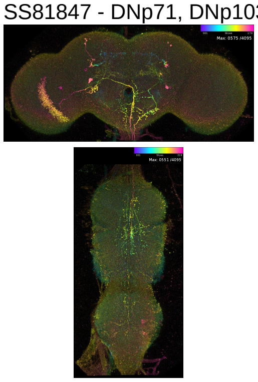

SS98644 - DNp75

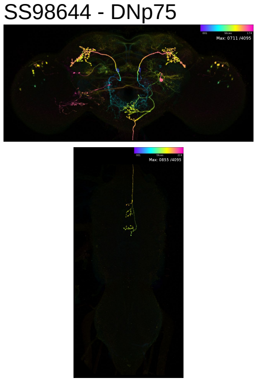

SS72185 - DNx01

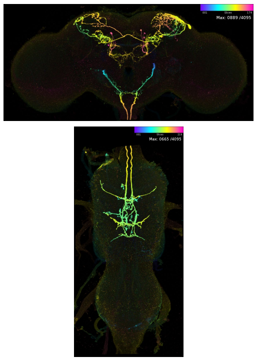

SS73430 - DNx01

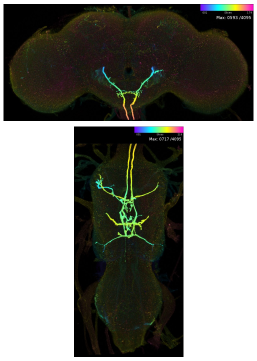
